# Lifestyle adaptations of *Rhizobium* from rhizosphere to symbiosis

**DOI:** 10.1101/2020.05.07.082560

**Authors:** Rachel M. Wheatley, Brandon L. Ford, Li Li, Samuel T. N. Aroney, Hayley E. Knights, Raphael Ledermann, Alison K. East, Vinoy K. Ramachandran, Philip S. Poole

**Author notes:** These authors contributed equally to this work (Rachel M. Wheatley, Brandon L. Ford, Li Li). Corresponding authors: Prof. Philip Poole, Department of Plant Sciences, University of Oxford, South Parks Road, Oxford, OX1 3RB, UK.; or Dr Vinoy Ramachandran, Department of Plant Sciences, University of Oxford, South Parks Road, Oxford, OX1 3RB, UK.

## Abstract

By analyzing successive lifestyle stages of a model *Rhizobium*-legume symbiosis using mariner-based transposon insertion sequencing (INSeq), we have defined the genes required for rhizosphere growth, root colonization, bacterial infection, N_2_-fixing bacteroids and release from legume (pea) nodules. While only 27 genes are annotated as *nif* and *fix* in *Rhizobium leguminosarum*, we show 603 genetic regions (593 genes, 5 tRNAs and 5 RNA features) are required for the competitive ability to nodulate pea and fix N_2_. Of these, 146 are common to rhizosphere growth through to bacteroids. This large number of genes, defined as rhizosphere-progressive, highlights how critical successful competition in the rhizosphere is to subsequent infection and nodulation. As expected, there is also a large group (211) specific for nodule bacteria and bacteroid function. Nodule infection and bacteroid formation require genes for motility, cell envelope restructuring, nodulation signalling, N_2_ fixation, and metabolic adaptation. Metabolic adaptation includes urea, erythritol and aldehyde metabolism, glycogen synthesis, dicarboxylate metabolism and glutamine synthesis (GlnII). There are separate lifestyle adaptations specific to rhizosphere growth (17) and root colonization (23), distinct from infection and nodule formation. These results dramatically highlight the importance of competition at multiple stages of a *Rhizobium*-legume symbiosis.

**Significance:** Rhizobia are soil-dwelling bacteria that form symbioses with legumes and provide biologically useable nitrogen as ammonium for the host plant. High-throughput DNA sequencing has led to a rapid expansion in publication of complete genomes for numerous rhizobia, but analysis of gene function increasingly lags gene discovery. Mariner-based transposon insertion sequencing (INSeq) has allowed us to characterize the fitness contribution of bacterial genes and determine those functionally important in a *Rhizobium*-legume symbiosis at multiple stages of development.

## Introduction

Biological N_2_ fixation carried out by bacteria provides ~65% of the biosphere’s available nitrogen (1), with the single greatest contribution coming from the symbioses between soil-dwelling bacteria of the *Rhizobiaceae* family and plants of the legume family (2–4). Rhizobia provide the plant with ammonium (NH_4_^+^) and receive carbon sources as an energy supply in return (5). This effectively functions as a bio-fertilizer for the legume host, increasing plant growth without the need for exogenously applied nitrogen (4–6). During symbiosis, rhizobia must adapt to a number of different lifestyles. These range from free-living growth in the rhizosphere, through root attachment and colonization, to passage along infection threads, differentiation into bacteroids that fix N_2_ and, finally, bacterial release from nodules at senescence.

Several decades of intense research have gone into understanding the mechanistic details of signalling between rhizobia and legumes leading to nodulation (7–9). Rhizobia respond to plant flavonoids by synthesizing lipochitooligosaccharides that are detected by plant LysM receptors to initiate nodule morphogenesis and concomitant rhizobial infection (9–12). Rhizobia typically, but not always, attach to the tips of developing root hairs, triggering root hair curling and entrapment of bacteria in an infection pocket (8). The plant cell plasma membrane invaginates to form an infection thread, down which bacteria grow towards the base of the root hair cell through continuous division at the leading edge. Infection is usually a clonal event (13), selecting one *Rhizobium* and allowing it to multiply to high numbers within the ramifying threads. Infection threads deliver rhizobia to the root cortex of the developing nodule. In symbiosis between *Rhizobium leguminosarum* bv. *viciae* and pea, bacteria are internalized by nodule cells and undergo terminal differentiation into N_2_-fixing bacteroids (10). Nodules provide a protective microaerobic environment to maintain oxygen-labile nitrogenase (7). In exchange for ammonia (NH_4_^+^) and alanine, the legume host provides carbon sources to fuel this process, primarily as dicarboxylic acids (14–15).

However, nodule infection is only one stage of the lifestyle of rhizobia and they spend much of their time surviving in the rhizosphere, the zone of soil immediately surrounding roots (16). Plants can release up to a fifth of their photosynthate via their roots (17), making the rhizosphere a nutritionally-rich zone, but one where there is strong selective pressure and competition for resources. Attachment and subsequent colonization of roots by rhizobia is almost certainly important to their long-term survival, but it is unclear how important these steps are to infection of legumes (18). Research over several decades has enabled identification of around 8 *nif* and 27 *fix* genes required directly for N_2_ fixation, as well as genes essential for recognition by the plant, differentiation and energization of N_2_ fixation. However, it has long been speculated these probably represent a fraction of genes needed by rhizobia for successful survival and growth in competition with other bacteria.

Rhizobiaceae have large genomes that probably reflect their adaptation to the heterogenous soil environment and dramatic lifestyle and developmental changes undergone in symbiosis (19–20). The 7.75 Mb genome of *R. leguminosarum* bv. *viciae* Rlv3841 consists of a circular chromosome (4,788 genes) and six plasmids: pRL12 (790 genes), pRL11 (635 genes), the symbiosis plasmid pRL10 (461 genes), pRL9 (313 genes), pRL8 (141 genes), and pRL7 (189 genes). Here we perform a multi-stage mariner transposon insertion sequencing (INSeq) genetic screen of Rlv3841 engaging in symbiosis with pea (*Pisum sativum*) at four different symbiotic lifestyle stages: 1) growth in the rhizosphere, 2) root colonization, 3) undifferentiated nodule bacteria, and 4) terminally differentiated bacteroids. This has allowed us to identify the genetic requirements for competitiveness at multiple developmental stages during symbiosis.

## Results and Discussion

### Mariner library construction, sequencing, and analysis

Rlv3841 mariner insertion mutants (input1) were inoculated onto 7-day-old pea seedlings and subsequently recovered at 5 days post inoculation (dpi) from; 1) the rhizosphere and 2) roots (Fig. 1). These samples (input1, rhizosphere, and root) were used to form libraries for INSeq analysis. From pea plants at 28 dpi (inoculated with input2), 150,000 nodules were picked from approximately 1,500 plants and two further libraries for INSeq analysis constructed; 3) undifferentiated nodule bacteria, generated by growing bacteria from crushed nodules (nodule bacteria that are not terminally differentiated along with those recently released from infection threads) for 12 h in rich medium before extracting DNA, and 4) bacteroids, generated from total nodule bacterial DNA (Fig. 1). These six libraries were analyzed (input1 and input2, together with the four test conditions; rhizosphere, root, nodule bacteria and bacteroids), by a Hidden Markov Model (HMM) (21). Each of the 7,357 Rlv3841 genes which contain TA insertion sites (99.9% of the genome), was classified as essential (ES), defective (DE), neutral (NE) or advantaged (AD) (*SI Appendix*, Dataset S1). Genes classified as NE in the respective input library (6,429 for input1 and 6,656 for input2) were considered for each test condition, with genes classified as either ES or DE (ES/DE) regarded as required at that stage of symbiosis and described thus throughout this manuscript.

**Fig. 1.**
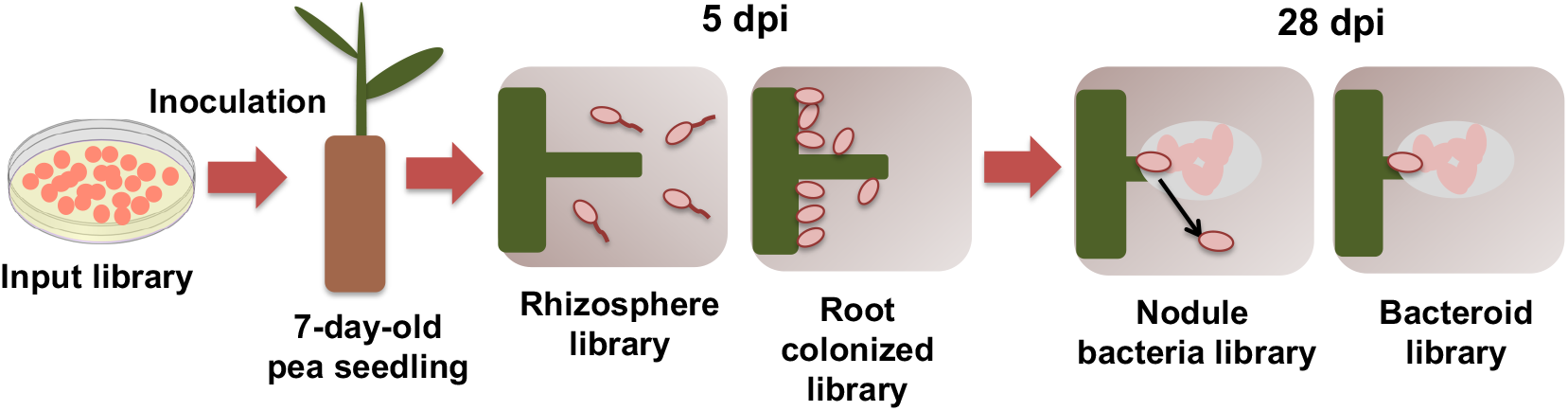
Collection steps following inoculation of pea plants with the INSeq input libraries. Bacterial DNA was purified from: 1) the rhizosphere (5 dpi), 2) colonized roots (5 dpi) following inoculation with input1 and from 3) nodule bacteria (28 dpi), and 4) bacteroids (28 dpi) following inoculation with input2. These four samples, together with DNA purified from input1 and input2, were used to make libraries for INSeq analysis.

Bacteria with a mutation which results in out-competition at any stage of early nodule formation will occupy fewer nodules on peas, and lead to an ES/DE classification. However, many mutants unable to fix N_2_, such as those defective in *nifH* (encoding Fe protein of nitrogenase), do not alter the competitiveness for nodule infection. Nonetheless, plant sanctioning results in the formation of smaller nodules (22). We therefore predicted at the outset of this work, and subsequently confirm, that mutants impaired specifically in N_2_ fixation but not nodule formation, the result is an ES/DE classification for the gene in bacteroids.

### Classification of symbiotic fitness determinants

Across all stages of symbiosis, INSeq reveals a total of 603 genetic regions (593 genes, 5 tRNAs and 5 RNA features) are required to competitively nodulate peas (Fig. 2). A total of 17 are required only in the rhizosphere (*SI Appendix*, Table S1), while 146 are classified as rhizosphere-progressive (*SI Appendix*, Table S2), required not only in the rhizosphere but also at every subsequent stage. Seven genes are required for both rhizosphere and root colonization (rhizosphere and root) (*SI Appendix*, Table S3), with 23 genes specifically required for root colonization (root) (*SI Appendix*, Table S4). Genes classified as rhizosphere or root required are important for fitness and persistence in those particular environments, but not linked to nodule infection, *per se*. There are 33 genes required from root colonization to nodule (root-progressive) (*SI Appendix*, Table S5), while 211 genes are required by both nodule bacteria and bacteroids (nodule-general) (*SI Appendix*, Table S6). There are an additional 142 specific for nodule bacteria i.e. required only for regrowth of bacteria from crushed nodules (*SI Appendix*, Table S7), with 24 specific for bacteroids (direct isolation from nodules without regrowth) (*SI Appendix*, Table S8) (Fig. 2).

**Fig. 2.**
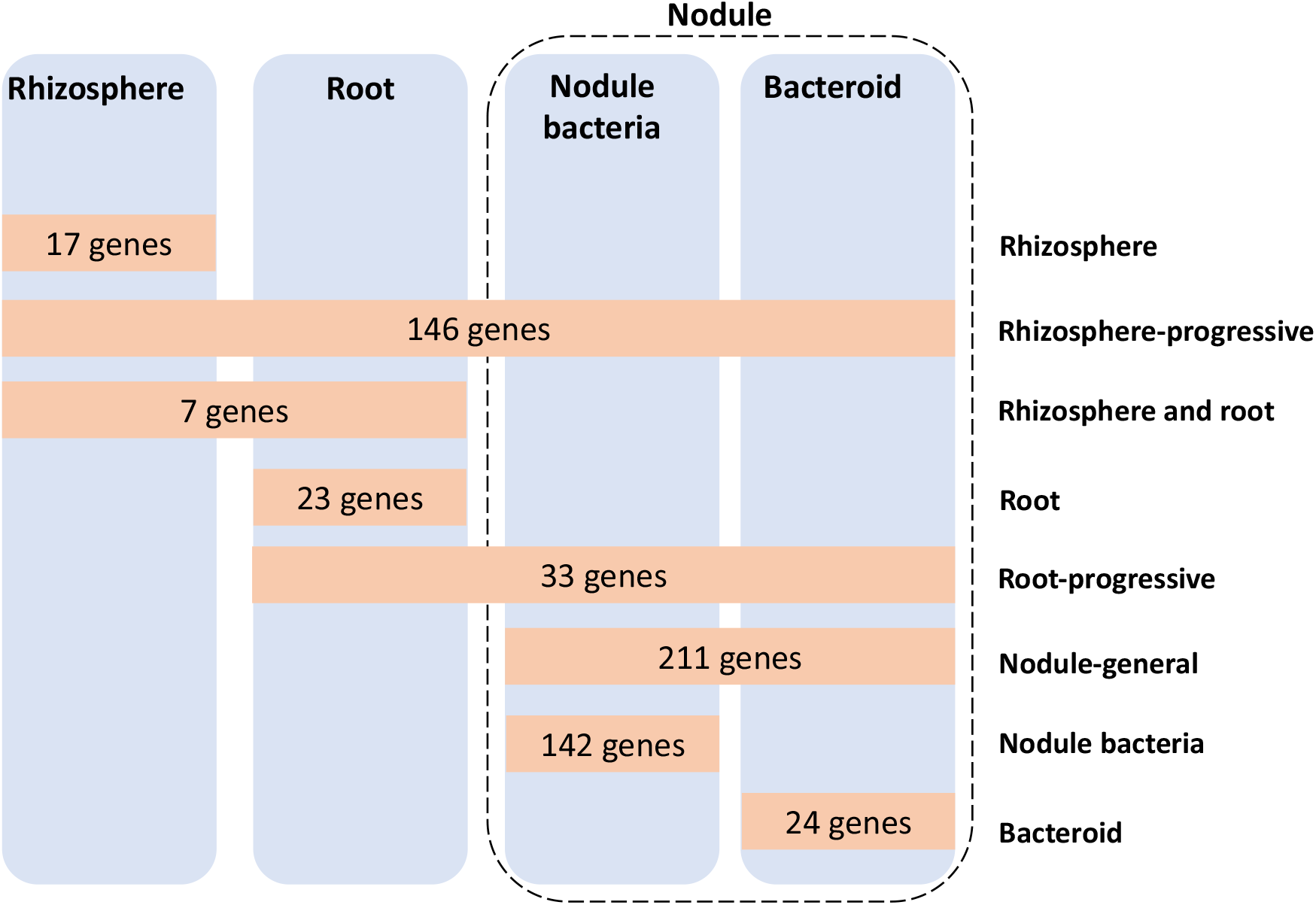
Rlv3841 genes required during the stages of symbiosis with pea. Orange boxes show the number at each symbiotic stage; rhizosphere (*SI Appendix*, Table S1), rhizosphere-progressive (*SI Appendix*, Table S2), rhizosphere and root (*SI Appendix*, Table S3), root (*SI Appendix*, Table S4), root-progressive (*SI Appendix*, Table S5), nodule-general (*SI Appendix*, Table S6), nodule bacteria (*SI Appendix*, Table S7), and bacteroid (*SI Appendix*, Table S8). All genes shown were classified neutral (NE) in the respective input library.

### Rhizosphere genes

The rhizosphere, the zone immediately surrounding plant roots, is an oxidative environment formed through the release of reactive oxygen species (23–24). Mutation of the 17 genetic regions required specifically in the rhizosphere (*SI Appendix*, Table S1) did not impair subsequent stages of root colonization or nodulation (Fig. 2). Rhizosphere-specific genes involved in the oxidative stress response, important for survival in this environment, encode a putative oxidoreductase (pRL110539), a superoxide dismutase (RL1340, *sodB*), and RNA-binding protein Hfq (RL2284) involved in control of post-transcriptional gene regulation. Persistence in the rhizosphere also required RL2964 (*cspA3*), encoding a cold shock protein. This rhizosphere-specific group also includes a single tRNA (tRNA^Gln^) and five genes predicted to encode metabolic or biosynthetic functions for; pRL80090, a fucose operon protein, pRL90136, a glycosyl transferase, pRL90204, an amidase, pRL100145, an acyl-CoA dehydrogenase, and RL3335, a lysophospholipase.

### Rhizosphere-progressive genes

The 146 rhizosphere-progressive genes (Fig. 2, *SI Appendix*, Table S2), required in the rhizosphere but also at subsequent stages, include genes for the biosynthesis of tryptophan (RL0021 (*trpB*), RL0022 (*trpA*), RL2493 (*trpD*), RL2494 (*trpC*), and RL3521 (*trpE*)), riboflavin (RL1621 (*ribG*) and RL1622 (*ribC*)), leucine (RL3513 (*leuA1*), RL4707 (*leuB*), and RL4732 (*leuS*)), isoleucine (RL1803 (*ilvD1*), RL3205 (*ilvC*), and RL3245 (*ilvI*)), and cobalamin (pRL110632 (*cobF*), RL2781 (*cobU*), RL2781A (*cobV*), RL2829 (*cobO*), RL4348 (*cobT*), and RL4349 (*cobS*)). There may be a specific requirement for these biosynthetic genes in the rhizosphere, with mutants subsequently uncompetitive at later stages of symbiosis, or alternatively, they may be required throughout all symbiotic stages. Riboflavin availability influences rhizobial survival in the rhizosphere and root colonization. An *R. leguminosarum* mutant in *ribN* (encoding putative riboflavin transporter protein RibN) showed lower nodule occupancy in pea plants (25).

The outer surface of rhizobia, composed of a variety of polysaccharides, including lipopolysaccharide (LPS) and exopolysaccharide (EPS) (18), is critical for both attachment to root hairs and host-specific recognition. Rhizosphere-progressive genes include those encoding putative proteins; O-antigen ligase (pRL90053), exopolysaccharide production protein (pRL110389, with high homology (97% identity (id)) to PssF of *R. leguminosarum* TA1 involved in polysaccharide export (26), and O-antigen transporter (RL0821). In addition, both RL3651 (*pssG*) and RL3664 (*pssN*) have well-defined roles in EPS biosynthesis (26), and their requirement, alongside pRL110389 (*pssF*) and RL1600 (*ppx*), highlight the importance of correct EPS processing not only in nodule infection but in all stages of interaction with the plant. The transcriptional regulator RL1379 (*rosR*), classified as rhizosphere-progressive, has a role in nodulation competitiveness (27). RosR regulates *pss* genes encoding EPS and *leguminosarum rosR* mutants produce only one third the EPS of wild type (28).

Other requirements include; RL1589 (*ropB*) and RL2775 (*ropA*) which have been identified to encode outer membrane amyloid fibrils in *R. leguminosarum* (29). These are conserved across numerous rhizobia (30–31), and are possibly required for either root attachment or transport. RL2256 (*ntrB*) and RL2257 (*ntrC*), encoding the NtrB/NtrC two component regulatory system, control expression of genes in response to nitrogen-limitation. While in this work pea seedlings were grown in nitrogen-free rooting solution to enable nodulation, this pathway is likely to be important in a natural soil environment where nitrogen availability is low. RL0422 (*rpoN*), the gene encoding RNA polymerase sigma factor 54, is classified as rhizosphere-progressive. Interestingly, this indicates that while essential for *nif* gene expression and dicarboxylate transport (32) (processes which occur in nodules), it may also be required for processes in the rhizosphere. Bacterial uptake of solutes is important, shown by the requirement for genes encoding ATP-binding cassette (ABC) transporters of nitrogen-containing ions (RL4400-4402), and ferric cations (RL4583, *fbpB*). Another ABC system is required (RL1003-4), but the solute transported is unknown. RL3557 (*bacA*) encoding the BacA transporter protein is well-established to be required for bacteroid development (33). BacA protects rhizobia against the antimicrobial activity of nodule-specific cysteine-rich (NCR) peptides produced by the host pea (34, 35), but clearly it is also important at earlier stages of interaction with the plant. The gene encoding the stringent response protein RelA (RL1506 (*relA*), which synthesizes ppGpp) was similarly required across all symbiotic stages, indicating its central importance in response to stress.

### Root colonization genes

The root library, formed by collecting bacteria from the entire pea root, consists of colonizing bacteria attached over the whole epidermal surface. The genes specific for rhizosphere and root (7 genes, *SI Appendix*, Table S3) are required for both bacterial rhizosphere growth and root colonization, while 23 genes are required only for root colonization (designated root, Fig. 2) (*SI Appendix*, Table S4). This latter group is important for attachment to roots (9, 18, 36) but not involved in nodule formation, indicating a pathway of root colonization, distinct from that of nodulation.

Root-specific genes include those encoding putative cell-envelope related functions for; pRL90052, polysaccharide biosynthesis O-antigen related protein, and RL3663 (*pssO*), a polysaccharide transport outer membrane protein. These cell-envelope related functions may actively facilitate adherence to the root surface or may be important for general cell fitness during colonization. RL2512, encoding a putative transmembrane protein, was specifically required for root colonization. RL2512 encodes a putative SecG which forms part of the general protein secretion (Sec) system in bacteria. The protein encoded by RL2512 shows homology to SecG in *Escherichia coli* (39% id over the N-terminal 71 amino acids) and *Agrobacterium tumefaciens* (73% id over 175 amino acids). Potential orthologues of the Sec system have been identified in Rlv3841 by homology with *E. coli* proteins (37). However, these chromosomal genes were not located in operons or even close to one another. SecG forms a protein-conducting channel in the SecYEG complex (37). Both the putative Rlv3841 genes encoding SecE (RL1759) and SecY (RL1794) were essential in the input libraries as well as across all stages of symbiosis. The different phenotype observed for RL2512 suggests there may be a second gene encoding a SecG-like function in Rlv3841 (although no gene was found with > 26% id).

### Root-progressive genes

Thirty-three root-progressive genes affect all stages from root colonization onwards (Fig. 2, *SI Appendix*, Table S5). They include RL2382 (*nodM/glmS*) encoding a fructose-6-phosphate aminotransferase that forms glucosamine-6-phosphate. There are two homologous *nodM* genes in *R. leguminosarum*, RL2382 and pRL100180, encoded on the chromosome and symbiosis plasmid, respectively (38). Biochemical assays demonstrated either protein provides sufficient glucosamine-6-phosphate (G6P) for lipochitooligosaccharide (LCO) Nod factor synthesis (38). A double *nodM* mutant in *R. leguminosarum* no longer nodulated pea or vetch (38). The two genes fall into different categories in this study; pRL100180 (*nodM*) is required only by nodule bacteria, while RL2382 (*nodM/glmS*) is root-progressive. Their different INSeq phenotypes suggest differential expression during symbiosis, consistent with plant-flavonoid induction of *nodM*.

This root-progressive group also includes two genes encoding putative cell-surface constituents for; RL0818, a lipopolysaccharide biosynthesis protein, and RL3677 (*ispL*), a UDP-glucuronate 5’-epimerase, required for glycoconjugation in EPS biosynthesis (39). This root-progressive designation suggests these two cell-surface related genes are required to produce the correct bacterial cell-surface needed for root colonization, as well as in subsequent steps, potentially root hair attachment, infection thread ramification or bacteroid formation in nodules. An ABC transporter (RL4538-39, *ccmB cycZ*) showing homology with a system exporting heme-binding proteins to the periplasm in *R. etli* (40) is required, together with RL4540 (*cycY*) a cytochrome *c* biogenesis protein, responsible for interchange of thiol:disulfide bonds. This suggests novel cytochrome biosynthesis and export is important during initial root colonization.

### Nodule-general genes

A total of 211 genes are classified nodule-general (Fig. 2, *SI Appendix*, Table S6). These are genes whose mutation impaired recovery from both nodule bacteria (i.e. regrown bacteria) and bacteroids (i.e. isolated from total nodule rhizobacterial DNA without regrowth). This group includes nine *nod* genes, encoding components for the biosynthesis and signalling pathways of LCO nodulation (Nod) factors (41). LCOs are diffusible molecules and *nod* mutants are still able to initiate symbiosis if co-inoculated 1:1 with wild type. However, in mass competition, such as this INSeq experiment, *nod* mutants may either have delayed establishment or reduced cell numbers within nodules, one or both of which could result in a nodule-general classification. The genes are organized in three operons on pRL10; pRL100178-pRL100180 (*nodTNM*), pRL100181-pRL100183 (*nodLEF*) and pRL100189-pRL100184 (*nodABCIJ-nodD*), and in the latter two clusters, every gene (9 in total) is classified as nodule-general (*SI Appendix*, Table S6). These genes include biosynthesis of Nod factor as well as export by the ABC transporter encoded by pRL100188 (*nodI*) and pRL100189 (*nodJ*).

The nitrogenase complex is encoded by genes pRL100162-pRL100158 (*nifHDKEN*) and pRL100196-pRL100195 (*nifAB*). These genes are classified as nodule-general (*nifDK* and *nifN*) (*SI Appendix*, Table S6) and bacteroid (*nifH, nifE*, and *nifAB*) (*SI Appendix*, Table S8). Although the classification of these latter four as bacteroid specific suggests that their mutation reduces N_2_-fixing bacteroids but not the level of undifferentiated bacteria within nodules, as these classifications are closely related, the division of functionally related clusters into separate groups should not be over-interpreted.

Electron transfer flavoproteins (ETF) transfer electrons to nitrogenase by bifurcation (42). Genes *fixABCX* (pRL100200-pRL100197) encoding ETF FixABCX, were required in the nodule, although, as above, they fall into two categories; nodule-general (*fixAB*) (*SI Appendix*, Table S6) and bacteroid (*fixCX*) (*SI Appendix*, Table S8). Again, we might tentatively conclude that while all four genes are needed for symbiotic N_2_ fixation, mutation of *fixC* and *fixX* do not alter the number of viable nodule bacteria. Mutation of ETF proteins abolishes N_2_ fixation in several rhizobia-legume symbioses (43).

The microaerobic environment required for the efficient functioning of the nitrogenase complex is achieved by the combined activities of plant leghemoglobin and the bacteroid’s high affinity *cbb*3 cytochrome complex. Rlv3841 has two clusters of *fix* genes that encode the components of *cbb*3, one on the symbiosis plasmid *fixN1O1Q1P1G1H1I1S1* (pRL100205-pRL100211) and the other on pRL9 (pRL90012-pRL90020, *fixN2O2Q2P2G2H2I2S2KL*). Only *fixN1* and *fixN2* were required (*SI Appendix*, Tables S6 and S2), suggesting the other duplicated components of the *cbb*3 complex can compensate for each other and, possibly, that the level of FixN limits the activity of the complex.

Differentiation of rhizobia into bacteroids requires significant reprogramming of the LPS and outer membrane (44). Genes with a predicted role in cell envelope functions formed an important group of nodule-general genes. These include RL3649 (*pssI*) and RL3650 (*pssH*), encoding putative glycosyltransferases, and RL3820, encoding a putative exopolysaccharide production protein. These proteins may restructure the cell surface during infection and bacteroid differentiation. O-antigen biosynthesis also appears to play a role in these structural changes, with pRL110056, encoding a putative UDP-N-acetyl-D-glucosamine 6-dehydrogenase involved in O-antigen biosynthesis classified as nodule-general. As with growth in the rhizosphere and root colonization, stress and defence response genes are also required in nodules. These include RL0824, encoding a putative oxidoreductase, RL0962, encoding a putative dioxygenase, RL0994 and RL1545, encoding putative reductases.

In terms of metabolism, nodule-general genes include pRL100433-pRL100434, which are predicted to encode allophanate hydrolase, catalyzing the hydrolysis of allophanate yielding ammonium and CO_2_. Allophanate can be either derived from cyanuric acid degradation or be an alternative to urease in degradation of urea (45), in which urea is first carboxylated to allophanate. Indeed, pRL100436 and pRL100437 (nodule bacteria-specific, *SI Appendix*, Table S6) are annotated to encode a biotin-dependent carboxylase, which may be a urea carboxylase. Urea carboxylase belongs to the biotin-dependent carboxylase family of enzymes, which has 40% id to urea carboxylase of *Saccharomyces cerevisiae* and *Oleomonas sagaranensis* (46). Overall, this suggests urea catabolism is important in nodule bacteria and bacteroids.

Genes pRL100246 and pRL100247, encoding a putative aldehyde dehydrogenase and adjacent conserved hypothetical protein, were nodule-general. This suggests aldehyde metabolism is important for optimal fitness in both nodule bacteria and bacteroids. Soybean nodules contain aldehydes (31 to 53 nM), which are converted to acetate by aldehyde dehydrogenase (47, 48). Erythritol utilization is also required in nodules; pRL120205 (*eryB*), encoding erythritol phosphate dehydrogenase and pRL120206 (*eryC*), encoding erythrulose 4-phosphate dehydrogenase are nodule-general. This agrees with the known reduction in nodule competitiveness of *ery* mutants (49). RL4117 (*glgA*) encoding glycogen synthase is nodule-general. Glycogen is an important carbon store in rhizobia and its disruption impairs growth on glucose-containing molecules (1).

Various transport systems are required in nodules. Genes required include those encoding pRL110047 (permease of ABC exporter), pRL90262 (ABC of a hydrophobic amino acid transporter (HAAT) ABC uptake system), RL0186-7 (permease components of peptide transporter (PepT) ABC uptake system), RL0226 and RL0228 (permease and solute binding protein (SBP) of a second PepT family) and RL3333 (ABC of an unclassified transporter). The importance of NCR peptides in development of mature bacteroids within N_2_-fixing nodules is well known (50). While we speculate that the two PepT uptake systems are involved in delivery of plant-derived peptides crucial for plant controlled bacteroid development, uptake by systems transporting unknown solutes and export of unwanted compounds is clearly also necessary.

### Nodule bacteria genes

Although most genes with known or obvious roles in nodule infection are classified as nodule-general or bacteroid, there are an additional 142 genes specific for nodule bacteria (Fig. 2, *SI Appendix*, Table S7). This INSeq library is made up of undifferentiated bacteria, likely to come from inside, or recently released from, infection threads, and capable of growth in TY (fully differentiated bacteroids are unable to re-grow in this way). Classification of genes as specific for nodule bacteria may be for one of two reasons; firstly, that they are needed for optimum bacterial growth within infection threads or secondly, that they are required for re-growth in TY medium, with implications for importance in bacterial release as nodules senesce.

This list of genes includes a number of flagella-related genes which are considered below. A number of defence and stress-related genes, including RL0392, encoding a putative hemolysin-like protein, RL2102 (*cspA5*) encoding a putative cold shock protein, and pRL110509 and RL2526 encoding putative oxidoreductases, are also specific for nodule bacteria. The gene encoding a resistance-nodule-division (RND) family efflux transporter (pRL120696) is required and its role is likely to be in defence, exporting toxic/unwanted chemicals from bacterial cells. The gene encoding an ABC export system RL4018 (ABC-permease fusion) may also be involved in a similar task. In fact, genes encoding components of several ABC uptake transporters are classified as nodule bacteria-specific including pRL100062 (permease of polyamine/opine/phosphonate transporter (POPT) family), pRL100072 (SBP of polar amino acid transporter (PAAT) family), pRL120071 (SBP of PAAT, which may transport galactosamine, glucosamine given its high homology to SMb21135 (51)), pRL120560 (ABC of carbohydrate uptake transporter 1 (CUT1) family), RL2659 (SBP, class unclassified), and RL3353 (ABC, ferric iron transporter (FeT)). The large number of transporters involved in uptake of amino acids, sucrose, iron and other unknown solutes may be required for optimum recovery from crushed nodules and subsequent growth in TY and their mutation consequently results in loss from the nodule bacteria library.

### Bacteroid genes

There are 24 genes specifically required by bacteroids (Fig. 2, *SI Appendix*, Table S8). Mutants in these genes have reduced fitness specifically in formation or maintenance of bacteroids but are not affected in bacterial progression through infection threads or regrowth from nodules. They include a number of *nif* (pRL100196-5 (*nifAB*), pRL100159 (*nifE*) and pRL100162 (*nifH*)) and *fix* (pRL100197-8 (*fixCX*)) genes which are needed for nitrogenase assembly and microaerobic respiration (52–53). This confirms our prediction that mutants impaired in N_2_ fixation would result in an ES/GD classification in the libraries recovered from nodules.

The dicarboxylate transport system (Dct) genes (RL3425 (*dctB*) and RL3426 (*dctD*)), enabling dicarboxylic uptake by bacteroids to fuel N_2_ fixation (7, 54, 55), are bacteroid-specific. Prior to bacteroid differentiation, rhizobia utilize a wide range of carbon sources and *dct* mutants form well-developed nodules and bacteroids (56). We assume that wild type and *dct* mutant strains maintain similar populations of viable bacteria in nodules (reflected in the nodule bacteria results), while *dct* mutants have severely reduced N_2_ fixation and numbers of bacteroids. RL0037 (*pckA*), encoding phosphoenolpyruvate carboxykinase, which is essential for growth on dicarboxylates (57), was also specifically required in nodule bacteroids (*SI Appendix*, Table S8). This fits with evidence that dicarboxylates drive bacteroid metabolism, but not bacterial growth during nodule infection where multiple carbon source are likely to be available. The cytochrome *bc1* complex is encoded by RL3484-86 (*petABC*), with RL3486 (*petA*), encoding the ubiquinol-cytochrome *c* reductase iron-sulfur subunit required across all symbiosis growth conditions (rhizosphere-progressive), whereas *petB* is classified as nodule-general. In contrast, RL3484 (*petC*), encoding the cytochrome *c*1 precursor, is specifically required in bacteroids. A mutant of RL3484 (*petC*), has been demonstrated to be Fix^−^in *R. leguminosarum* (58). In addition, RL3577 (*guaD*), encoding guanine deaminase, an enzyme that catalyzes the hydrolytic deamination of guanine producing xanthine and ammonia, is also bacteroid-specific (*SI Appendix*, Table S8).

### Mutants with improved bacterial colonization or increased numbers in nodules

When mutation in a gene leads to a bacterial strain with improved fitness for a given condition or environment, its INSeq classification is AD (AD genes at each stage from rhizosphere to nodule (NE or GD in input library), are summarized in *SI Appendix*, Fig. S1 and Table S9). No genes are classified as AD across all four conditions (rhizosphere-progressive) and none are root-progressive. However, there are considerable groups where mutation leads to better recovery from the rhizosphere (84 genes), and/or increased colonization of roots (172 and 33 genes, respectively), as well as leading to an increased population recovered from crushed nodules; nodule-general (11 genes), nodule bacteria-specific (following re-growth, 207 genes) or bacteroid-specific (22 genes) (*SI Appendix*, Fig. S1 and Table S9). A large proportion (approx. 66%) of AD genes are contiguous and clustered, with many in operons. In contrast to the genes required at a given stage, the reason for an AD classification is far more nuanced. Mutation of genes involved in uptake, sensing or metabolism of specific compounds can be advantageous to bacteria in a given environment, increasing their fitness, as energy is not expended on pathways or gene expression that is not needed. For example, in contrast with rapid growth in rich broth *in vitro*, when bacteria are growing slowly, their metabolism will be more restricted and there will be a greater penalty for synthesizing proteins not absolutely required. AD genes are likely to reflect this. In the rhizosphere, AD-classified genes (84) include ABC homoserine transporter pRL80088, Major Facilitator Superfamily (MSF) nitrate transporter NarK (RL1992), ABC nitrate transporter (NitT) family SBP RL1993, and nitrate reductase transcriptional regulator NasT (RL1994), as well as a large region of 26 genes (RL1870-95) including genes encoding sensor regulators, heat shock proteins, amongst others.

Rhizosphere and root AD genes (172, *SI Appendix*, Fig. S1 and Table S9) include ABC transporters involved in uptake of quaternary amines (QAT) pRL100078-83 (*qatV6W6X6*); peptides, PepT-family pRL110053-54 (*optAD*), pRL110281and pRL120330; amino acids, pRL120404 (*braC2*) involved in uptake of branched chain amino acids (59) (HAAT), general L-amino uptake system RL2203-4 (*aapQJ*, polar amino acid (PAAT) family) and pRL80060 (PAAT SBP with homology to a mimosine transporter of *Rhizobium* sp. TAL 1145 (60); carbohydrates RL2895-6 (CUT1); phosphate uptake transporter (PhoT) RL1683 and polyamine/opine/phosphonate uptake (POPT) family RL2009-14. Under these specific experimental conditions, in the rhizosphere and on roots, bacteria can clearly survive without the wide range of compounds that these transporters import. Export systems, RND-family efflux proteins pRL100286-7, and multi-drug efflux transporter RL3414 (*norM*) are also classified AD under these conditions.

There is a small group of nodule-general AD transport genes, including CUT1 uptake system RL0749 (*aglK*) transporting alpha-glucosides and exporter RL3632. This later is part of a cluster of five genes (RL3628-32) which includes a sugar decarboxylase (RL3628) and three glycosyl transferases (RL3629-31).

When we consider nodule bacteria, even slight changes in the terminal differentiation to bacteroids will affect the proportion of bacteria recovered from nodules. Mutation of a gene which leads to delay of differentiation might lead to an AD gene classification. However, as preparation of the nodule bacteria sample includes recovery in TY broth, any conclusions should be tentative as the influence may be to increase fitness for regrowth in a nutrient rich medium. Nodule bacteria AD genes (207) include numerous ABC and other transporter systems; CUT1 family systems pRL100440-pRL100443 contiguous with an inositol degradation protein (pRL100439), pRL12006-7, pRL120557-58 contiguous with a glycosyl hydrolase (pRL120559), and RL4375-8; CUT2 family systems *eryGFE* (pRL120200-2) together with *eryH* (pRL120203) a periplasmic lipoprotein involved in erythritol uptake and/or metabolism, pRL90246-7; HAAT family components pRL120095 and RL3745 (*braC*) (involved in uptake of branched chain amino acids (59)), PepT complete systems pRL110513-7 and pRL90249-53 which could be involved in uptake of NCR peptides as without their uptake, differentiation into bacteroids might be reduced leading to increased bacterial recovery. Tri-partite ATP-independent transporter (TRAP) import system RL4359-61 is likely to be involved in the uptake of methyl pyruvate or pyruvate by homology with *S. meliloti* (51). For uptake of metals the following genes are classified AD in nodule bacteria; Iron chelate uptake transporter (FeCT) components RL2713-5; FeT family RL3350-2; Manganese/Zinc/Iron chelate transporter (MZT) family components RL3885-6 (*sitAB*), likely to import manganese (from homology to *meliloti* transporter (51)) and MgtE-like family cationic transporter RL2550. In addition, AD genes include clusters encoding three extracytoplasmic function (ECF) sigma factors and their associated proteins involved in signal transduction from the external environment; RL1471-7 including *ecfDdgpfD1D2D3*, RL3233-5 including *ecfElppE*, RL3508-12 including *lppFecfFasfF.* These sigma factors control, often large, networks of genes, with their mutation meaning energy is not consumed by unrequired gene expression, thus providing an advantage in the test environment. Genes involved in motility and chemotaxis classified as AD are discussed in detail below.

Bacteroid AD genes (22) include another ECF sigma factor, EcfO (pRL110418), together with nearby gene pRL110416 (*rhaI*) encoding a rhamnose isomerase involved in competition for nodulation in *R. leguminosarum* bv. *trifolii* (61). Contiguous gene pRL110415 (*rhaD*) encodes a dehydrogenase required for nodulation (*SI Appendix*, Table S6), although the contiguous rhamnose transporter (pRL110410-3), which might have been expected to be required for nodulation based on the results with *R. leguminosarum* bv. *trifolii*, is not required at any stage examined (*SI Appendix*, Datasheet 1). This is presumably because there is redundancy in Rlv3841 where another uptake system (most likely a CUT2) is able to import rhamnose, but the metabolism carried out by pRL110415 is needed for successful nodule formation.

### Importance of motility and chemotaxis during symbiosis

The genome of Rlv3841 contains up to 93 genes with putative roles in chemotaxis and motility, arranged in seven clusters of two genes or more, and 28 single genes (*SI Appendix*, Table S10). Genes encoding putative proteins for chemotaxis (*che*, 16 genes), methyl-accepting chemotaxis receptors (*mcp* (24 genes) and *mcr* (3 genes)) and, flagellar biosynthesis and motor function (*fla* (10 genes), and *flg* (16 genes), *flh* (2 genes), *fli* (12 genes), *mot* (5 genes)) have a variety of classifications throughout symbiosis, although the majority seem to fall into three clear categories: either not required for symbiosis (39 genes), required for nodule formation and function (nodule-general, 39 genes, *SI Appendix*, Table S6) or their mutation is advantageous in some way for growth in the rhizosphere and root colonization (12 genes) (*SI Appendix*, Table S10).Whilst ten motility-related genes are encoded on plasmids in Rlv3841 (seven on pRL12, and one each on pRL8, RL10, and pRL11), only chromosomally encoded motility-related genes are required for symbiosis.

Each of the individual genes in *che1* cluster (RL0686-93 (*icpA cheX1Y1A1W1R1B1Y1D1*)) is classified as nodule-general (*SI Appendix*, Table S6, S10), which means that they are all needed for nodule function, and also, perhaps counter intuitively, that the cluster is not required for optimal colonization of the rhizosphere or roots (Fig. 3). This suggests Che1 is either required for chemotaxis specifically to root hairs or early progression into infection threads but is not essential for colonization of the root surface. In contrast, all genes in the *che2* cluster RL4028-36 (*cheX2Y2A2W2R2B2Y2D2*) are classified AD in nodule bacteria (Fig. 3, *SI Appendix*, Table S9, S10). While the reason is not known, we speculate that the *che2* gene cluster is involved in the later stages of bacterial movement through infection threads. A mutation which delays progression from infection threads to differentiated bacteroids would result in their over-representation in the nodule bacteria library, leading to the genes’ classification as AD.

**Fig. 3.**
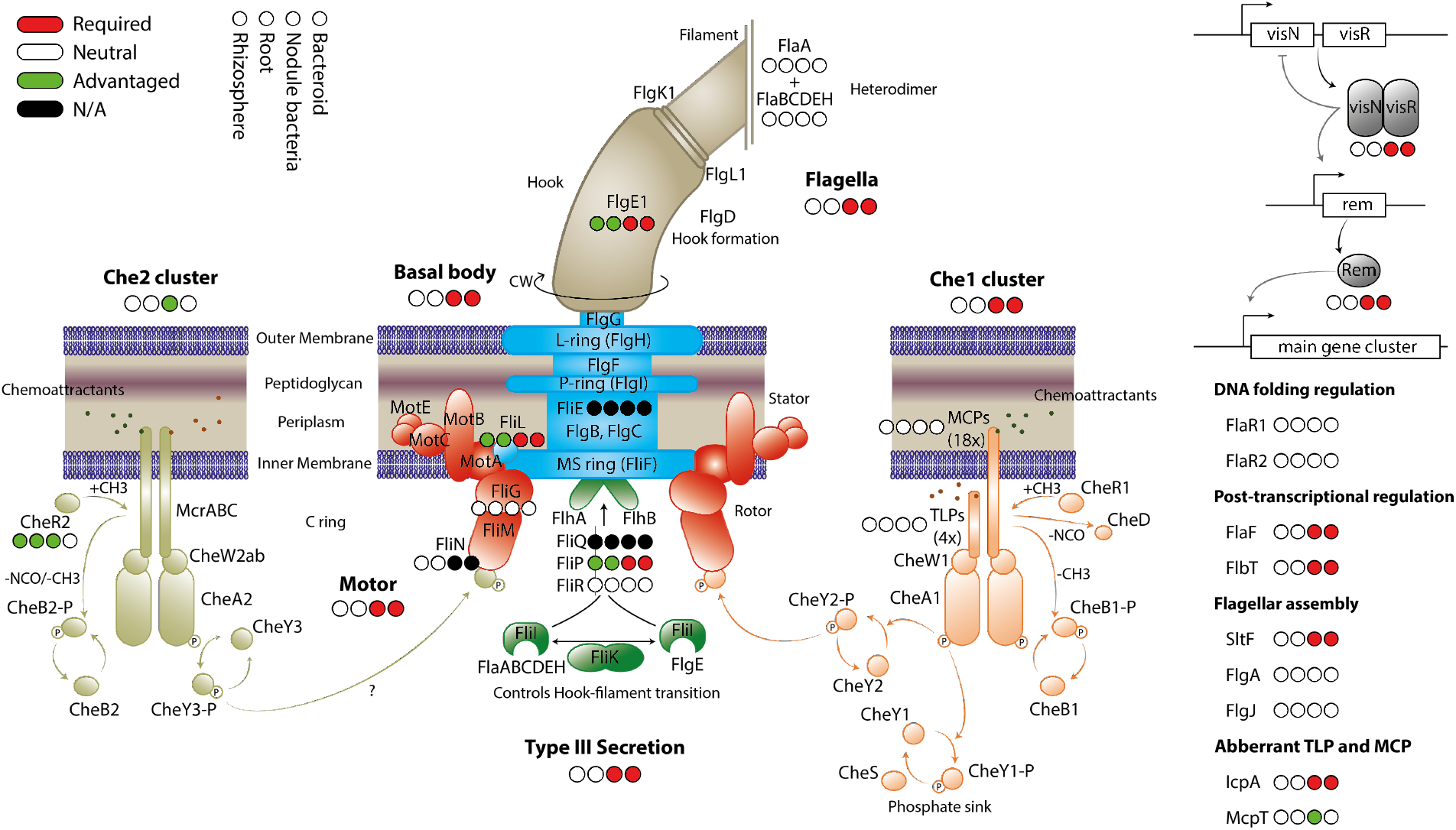
Classification of genes encoding motility and chemotaxis proteins in rhizosphere, root, nodule bacteria and bacteroid INSeq libraries. Genes required at a given stage are shown in red, genes where mutation has no effect (i.e. NE) are shown in white, genes which when mutated are advantageous are in green and those unclassified due to too few TA sites are in black. (INSeq results are shown in *SI Appendix*, Table S10). The proteins are colored by functional group, but the color is arbitrary with respect to INSeq phenotype.

The majority of flagellar assembly and basal body apparatus proteins are important in nodules, both in bacteroids and nodule bacteria (Fig. 3, *SI Appendix*, Table S10). The gene encoding critical motor protein FliG (RL0700 *fliG*) is neutral across all stages of symbiosis, however in contrast, the gene encoding an interacting integral motor protein MotA (RL0703) is required in nodule bacteria and bacteroids (Fig. 3). In addition, the genes encoding master flagella and chemotaxis regulator pair VisNR (RL0696 – RL0697) are required in nodule bacteria and bacteroids (Fig. 3), although the C-terminal GerE-encoding domain of *visN* displays a dramatic increase in transposon insertions, suggesting its deletion does not impair VisN function, and may even enhance it. A number of flagellar biosynthesis-encoding genes required in nodule bacteria are not needed, or their mutation is even advantageous, in the rhizosphere and for root colonization. Non-motile bacteria may become stranded in the rhizosphere or on the root surface. Overall, it suggests that under the experimental conditions, motility and chemotaxis are not essential for colonization of the root surface, but may have a particularly important role in the nodule infection pathway (Fig. 3). While emphasizing a very important role for motility and chemotaxis towards root hairs or, possibly, progression down infection threads, it does not preclude a more general role in the environment. INSeq requires a large inoculum (10^5^ cfu) onto sterile vermiculite-grown plants. However, growth in soil, in competition with a complex microbiome, is likely to impose different motility and chemotaxis selection upon the bacteria, with loss of chemotaxis and motility likely to be catastrophic for long-term survival in soil and root colonization (62).

### Plant rescue of auxotrophic rhizobia

While it is generally accepted that terminally differentiated bacteroids are mainly supplied with dicarboxylic acids, the metabolic status of rhizobia during infection remains largely enigmatic. Root exudates contain a large variety of amino acids, sugars, organic acids, purines, and pyrimidines (63–64), however, it remains unknown what compounds are available in substantial amounts and used by rhizobia. Therefore, we compared INSeq results from Rlv3841 grown in minimal medium (65), with symbiosis to identify auxotrophic mutants rescued by plant-derived compounds (*SI Appendix*, Table S11). We identified the main biosynthetic pathways for 18 of 20 amino acids and key components for purine, pyrimidine, S-adenosyl-methionine, heme, vitamin B12, Coenzyme A, Riboflavin and FMN/FAD, pyridoxal, and tetrahydrofolate biosynthesis (*SI Appendix*, Table S11). These pathways are required throughout all stages tested (i.e. rhizosphere-progressive), with the exception of purine biosynthesis where initial steps to 5-aminoimidazole-4-carboxamide-1-beta-D-ribofuranosyl 5’-monophosphate (AICAR) are only required in nodules and not in the rhizosphere or on roots (*SI Appendix*, Table S11). As the genes encoding the enzymes of these pathways are rhizosphere-progressive, root exudates presumably contain insufficient amounts of these compounds to sustain competitive growth of these mutants. Only AICAR (or an equivalent metabolite) is released at sufficient levels into the rhizosphere and onto root surfaces to enable growth, however not in sufficient amounts for infection thread or nodule development and function. Interestingly several genes, NE in minimal or rich (TY) medium are required during stages of symbiosis. Rlv3841 has at least two glutamine synthases, and of these RL3549 encoding GlnII, is NE in minimal medium and in colonization of roots or rhizosphere but is GD/ES in nodules and nodule recovered bacteria (i.e. nodule-general) (*SI Appendix*, Table S6). This is intriguing and suggests that increased glutamine synthesis may be required during infection. Similarly, there are two redundant copies of *argG* (RL2987 and RL4515). ArgG catalyzes the condensation of citrulline and aspartate to argininosuccinate in the urea cycle as part of arginine biosynthesis. As RL4515 is nodule-general (*SI Appendix*, Table S6) but is NE in minimal medium or during colonization, this suggests an increased role for arginine synthesis or degradation during bacteroid formation or function. Intriguingly, urea is also derived from arginine (see above) and urea catabolism is classified nodule-general. This highlights a potential role for arginine and urea catabolism via allophanate (urea-1-carboxylate) in nodule bacteria and /or bacteroids.

Histidine biosynthesis, requires formation of imidazole-glycerole-3-phosphate from phosphoribulosyl-formimino-AICAR-phosphate, catalyzed by heterodimeric HisFH. Two copies are encoded in the genome, one of which (RL0042/RL0046) is required for growth in the input conditions. Of the second copy, RL0819 is nodule-general (*SI Appendix*, Table S6) and RL0820 is rhizosphere-progressive. This indicates an increased need for these reaction steps or possibly an incompatibility with other enzymes under N_2_-fixing conditions. Glutathione biosynthesis genes (RL0855 (*gshA*), RL0338 (*gshB*), and RL1571 (*pepA*), *SI Appendix*, Table S11) are rhizosphere-progressive, (*SI Appendix*, Table S2), but are dispensable in minimal medium, indicating that when interacting with plants rhizobia must cope with increased oxidative stress (66–67).

### Validation of INSeq predictions

Fifteen genes, identified by INSeq as required in different stages of colonization and symbiosis were mutated by pK19 insertion and their mutant phenotypes determined experimentally (Table 1). Mutants in genes classified as root-progressive (pRL100053 and pRL80032) showed reduced colonization of roots compared to wild type (Fig. 4 A-B), but no significant reduction in survival in the rhizosphere (wild type 2.88 × 10^7^ cfu/ml SEM: 3.84 × 10^6^ (*n*=15) and mutant in pRL100053 1.81 × 10^7^ cfu/ml SEM: 2.57 × 10^6^ (*n*=8); wild type 2.13 × 10^8^ cfu/ml SEM: 3.20 × 10^7^ (*n*=3) and mutant in pRL80032 2.22 × 10^8^ cfu/ml SEM: 3.10 × 10^7^ (*n*=3)). Mutants in genes classified as nodule-general, together with those specific for nodule bacteria and bacteroids (Table 1), were tested experimentally for ability to nodulate and to fix nitrogen in single strain inoculations. Of these 13 mutants, ten nodulated peas at levels similar to wild type (Fig. 4C). The exceptions were mutants in *pssD*, which formed no nodules, and *nifB* and *fixB* which induced approximately twice as many, but small white, nodules as wild type (Fig. 4C). Assessing nitrogenase activity, the *nifB*, *fixB* and *pssD* mutants had no detectable acetylene reduction, while all other mutants showed activity similar (approx. 80-130%) to that of wild type (Table 1, Fig. 4D). These three mutant phenotypes, show clearly that *pssD, nifB and fixB* are required for nodulation and N_2_ fixation. It is also apparent why mutants in these genes have been lost from nodule bacteria and bacteroid libraries (Fig. 4C), as less DNA would be present in the nodules collected from strains that form smaller nodules (*nifB*, *fixB*) or are unable to form nodules at all (*pssD*). The reason for the classification of the other seven genes as nodule-general (Table 1) was unclear until experiments were performed to test their mutant’s ability to compete against wild type to form nodules. Mutant strains (marked with *gusA*) were competed against wild type *celB*-marked bacteria by inoculation in a 1:1 ratio. Six of the seven mutants tested were significantly impaired in competitive nodulation compared with wild type (Fig. 4E). A competitive disadvantage was evident despite the fact that a 1:1 inoculum provides significantly less competition than the INSeq environment where up to 10^5^ strains, with mutations in any one of over six thousand genes (6,429/6,656 classified as NE in input1/input2), compete to form nodules. For these six strains we have shown that the gene mutated in each case is required for effective symbiosis and these INSeq experiments have identified the stage at which it is needed. A single strain (RL1545 mutant), showed no reduction in nodule occupancy in inoculation at this competition ratio and we can speculate that its deficiency (it encodes a dehydrogenase/reductase) may only become apparent under stiffer competition conditions.

**Table 1.**
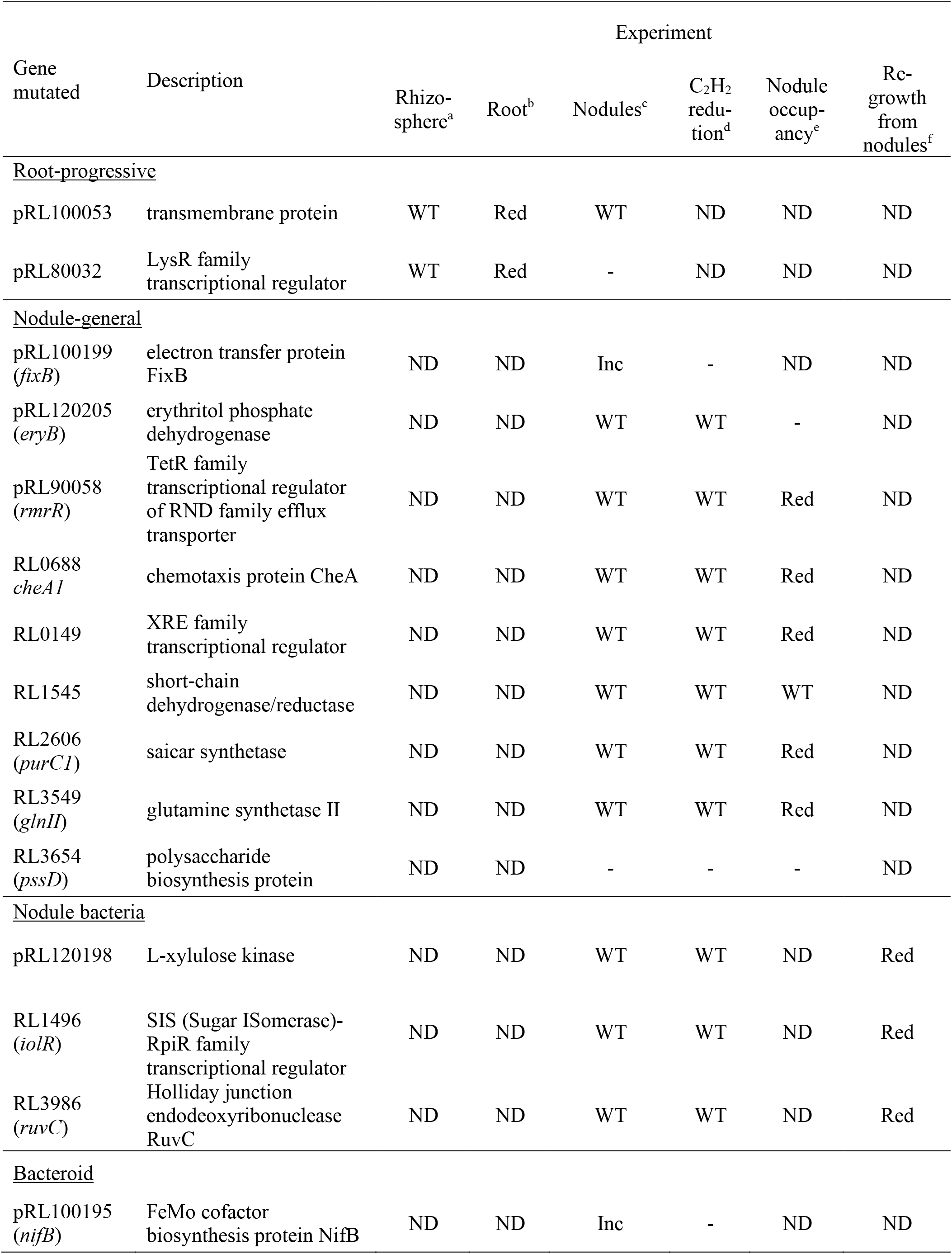
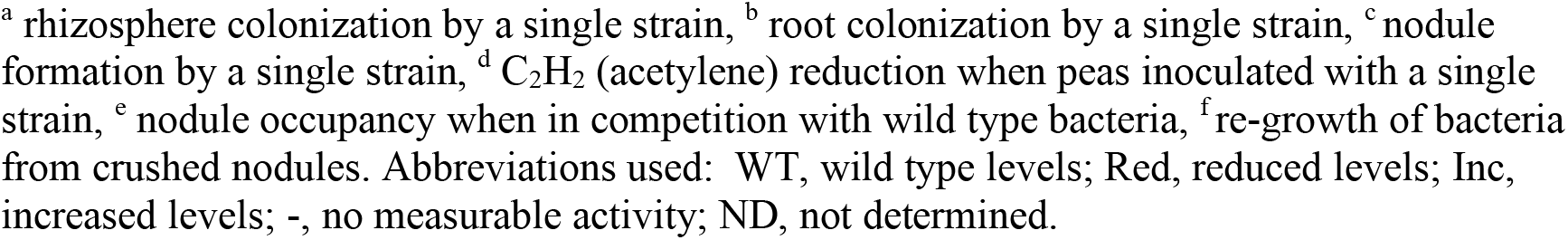
Phenotype of mutants affected at different stages of symbiosis.

| Gene mutated | Description | Experiment |  |  |  |  |  |
| --- | --- | --- | --- | --- | --- | --- | --- |
|  |  | Rhizo-<br>sphere<sup>a</sup> | Root<sup>b</sup> | Nodules<sup>c</sup> | C<sub>2</sub>H<sub>2</sub><br>redu-<br>tion<sup>d</sup> | Nodule<br>occup-<br>ancy<sup>e</sup> | Re-growth<br>from<br>nodules<sup>f</sup> |
| <u>Root-progressive</u> |  |  |  |  |  |  |  |
| pRL100053 | transmembrane protein | WT | Red | WT | ND | ND | ND |
| pRL80032 | LysR family<br>transcriptional regulator | WT | Red | - | ND | ND | ND |
| <u>Nodule-general</u> |  |  |  |  |  |  |  |
| pRL100199<br>(*fixB*) | electron transfer protein<br>FixB | ND | ND | Inc | - | ND | ND |
| pRL120205<br>(*eryB*) | erythritol phosphate<br>dehydrogenase | ND | ND | WT | WT | - | ND |
| pRL90058<br>(*rmrR*) | TetR family<br>transcriptional regulator<br>of RND family efflux<br>transporter | ND | ND | WT | WT | Red | ND |
| RL0688<br>*cheA1* | chemotaxis protein CheA | ND | ND | WT | WT | Red | ND |
| RL0149 | XRE family<br>transcriptional regulator | ND | ND | WT | WT | Red | ND |
| RL1545 | short-chain<br>dehydrogenase/reductase | ND | ND | WT | WT | WT | ND |
| RL2606<br>(*purC1*) | saicar synthetase | ND | ND | WT | WT | Red | ND |
| RL3549<br>(*glnII*) | glutamine synthetase II | ND | ND | WT | WT | Red | ND |
| RL3654<br>(*pssD*) | polysaccharide<br>biosynthesis protein | ND | ND | - | - | - | ND |
| <u>Nodule bacteria</u> |  |  |  |  |  |  |  |
| pRL120198 | L-xylulose kinase | ND | ND | WT | WT | ND | Red |
| RL1496<br>(*iolR*) | SIS (Sugar ISomerase)-<br>RpiR family<br>transcriptional regulator | ND | ND | WT | WT | ND | Red |
| RL3986<br>(*ruvC*) | Holliday junction<br>endodeoxyribonuclease<br>RuvC | ND | ND | WT | WT | ND | Red |
| <u>Bacteroid</u> |  |  |  |  |  |  |  |
| pRL100195<br>(*nifB*) | FeMo cofactor<br>biosynthesis protein NifB | ND | ND | Inc | - | ND | ND |

<sup>a</sup> rhizosphere colonization by a single strain, <sup>b</sup> root colonization by a single strain, <sup>c</sup> nodule formation by a single strain, <sup>d</sup> C<sub>2</sub>H<sub>2</sub> (acetylene) reduction when peas inoculated with a single strain, <sup>e</sup> nodule occupancy when in competition with wild type bacteria, <sup>f</sup> re-growth of bacteria from crushed nodules. Abbreviations used: WT, wild type levels; Red, reduced levels; Inc, increased levels; -, no measurable activity; ND, not determined.

**Fig. 4.**
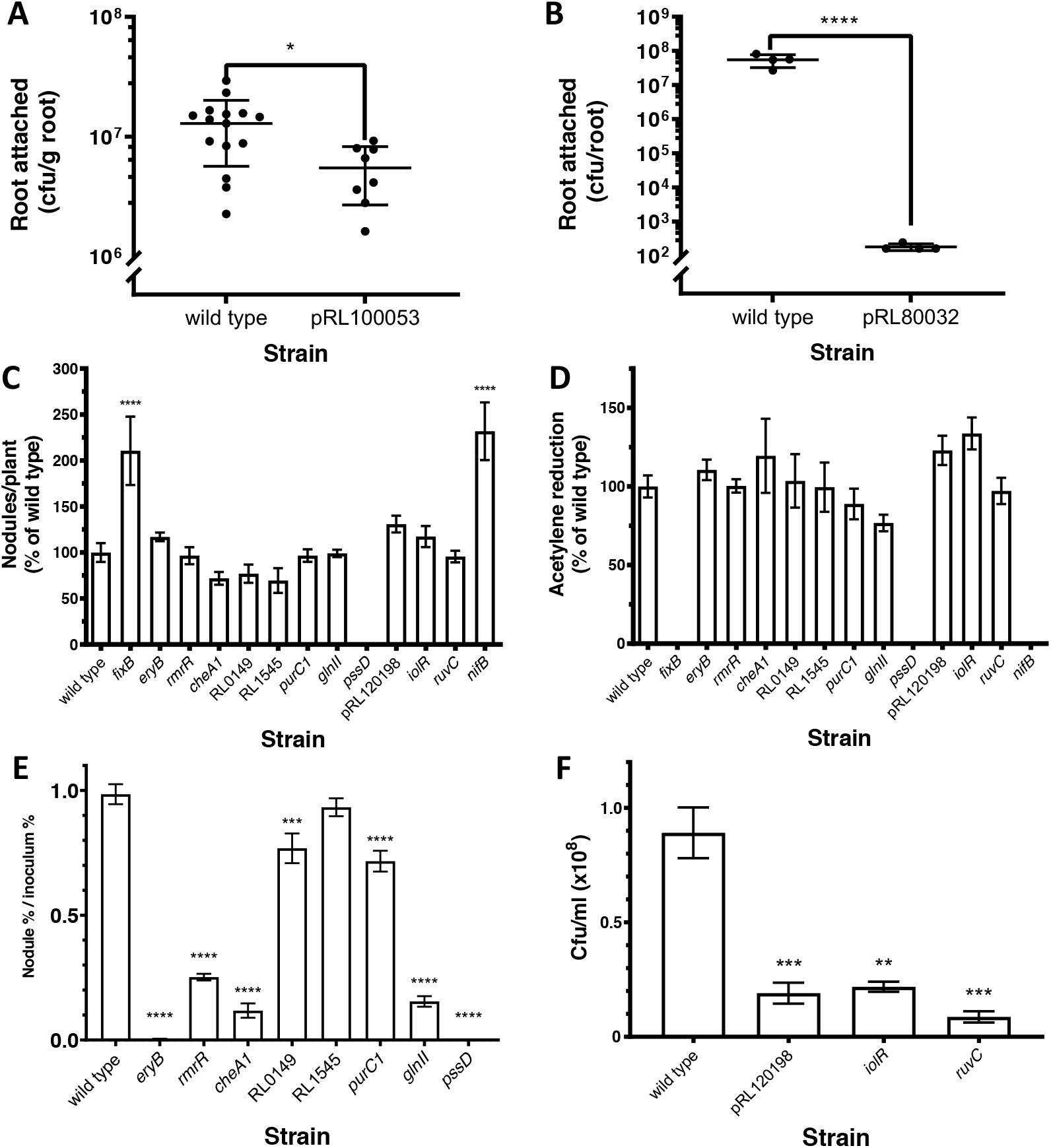
Experimental characterization of mutant phenotypes predicted from INSeq data. (*A*) Root colonization by bacteria (cfu/g root) from a single-strain inoculation (10^5^ cfu) of a 7-d-old pea plant comparing wild type and a mutant in pRL100053 (classified as root-progressive) at 5 dpi. *n* ≥ 8. An unpaired T-test was used to assess statistical significance. *(B*) Root colonization by bacteria (cfu/root) from a single-strain inoculation (10^5^ cfu) of a 7-d-old pea plant comparing wild type and a mutant in pRL80032 (root-progressive) at 5 dpi. *n* ≥ 4. An unpaired T-test was used to assess statistical significance. (*C*) Nodule formation on peas (nodules/plant as a percentage of wild type (Rlv3841GusA, 180 ±19.2 nodules/plant)) from a single-strain inoculation (10^4^ cfu) for mutants in nodule-general genes (9), nodule bacteria genes (3), and a bacteroid gene (Table 1) at 28 dpi. *n* ≥ 3. A Dunnett’s test comparing each mutant to wild type was used to assess statistical significance. (*D*) Acetylene reduction as a percentage of wild type (Rlv3841GusA, 5.18 ± 0.380 µmoles ethylene /plant/hr) of mutants in nodule-general genes (9), nodule bacteria genes (3), and a bacteroid gene (Table 1) at 28 dpi. *n* ≥ 3. A Dunnett’s test comparing each mutant to wild type was used to assess statistical significance. (*E*) Competition for nodule occupancy of mutants with wild type (nodule percentage/inoculum percentage) from 1:1 (mutant:wild type) co-inoculation (total 10^4^ cfu) of pea plants with eight nodule-general mutants harvested at 21 dpi. *n* ≥ 5. Statistical significance was assessed by a mixed effects model comparing the proportion of each strain in inoculum and nodules. (*F*) Bacterial recovery (cfu/ml (×10^8^)) from crushed nodules (10 per plant) and 36 h growth in TY broth at 28°C for wild type and three nodule bacteria mutants. *n* ≥ 3. To assess statistical significance, a Dunnett’s test comparing each mutant strain to wild type was used. * *P* < 0.05, ** *P* < 0.01, *** *P* < 0.001, **** *P* < 0.0001. Error bars show ±SEM. cfu; colony-forming unit.

For genes classified as nodule bacteria-specific we tested re-growth in TY medium since the initial preparation of this library involved a recovery step of growth in TY medium after nodule crushing. Wild type and mutants in three nodule bacteria-specific genes (pRL120198, RL1496 (*iolR*), and RL3986 (*ruvC*)) were grown in TY after extraction from nodules and compared. All mutants had a significantly lower rate of re-growth from crushed nodules than wild type (Fig. 4F). While this explains the low recovery of each of these mutants in this library, we did not identify any genes that increased the ratio of bacteria to bacteroids in nodules in such a small sample set. Future studies using flow cytometry to measure this ratio could be a powerful approach in such analyses.

## Conclusion

Here we present an in-depth investigation of N_2_-fixing bacterial interactions with a host legume during symbiosis by examining bacterial gene requirements at different stages of rhizosphere growth, root colonization, and nodulation. Key observations are summarized in Fig. 5, and emphasize the progressive nature of many mutations affecting stages leading to effective symbiosis. The power of INSeq is that it enables identification of those genes required when in competition with other bacteria. This is important when investigating environmental competition, since a mutation may not be deleterious for subsequent nodule formation or effective symbiosis in the artificial environment of inoculation with a single strain. It is sobering to appreciate that each of the 603 genetic regions required for competitive nodulation is individually just as important for N_2_ fixation as the structural components of nitrogenase. This INSeq dataset provides a powerful resource for better understanding the genetic basis of the crucially important N_2_-fixing rhizobia-legume symbioses.

**Fig. 5.**
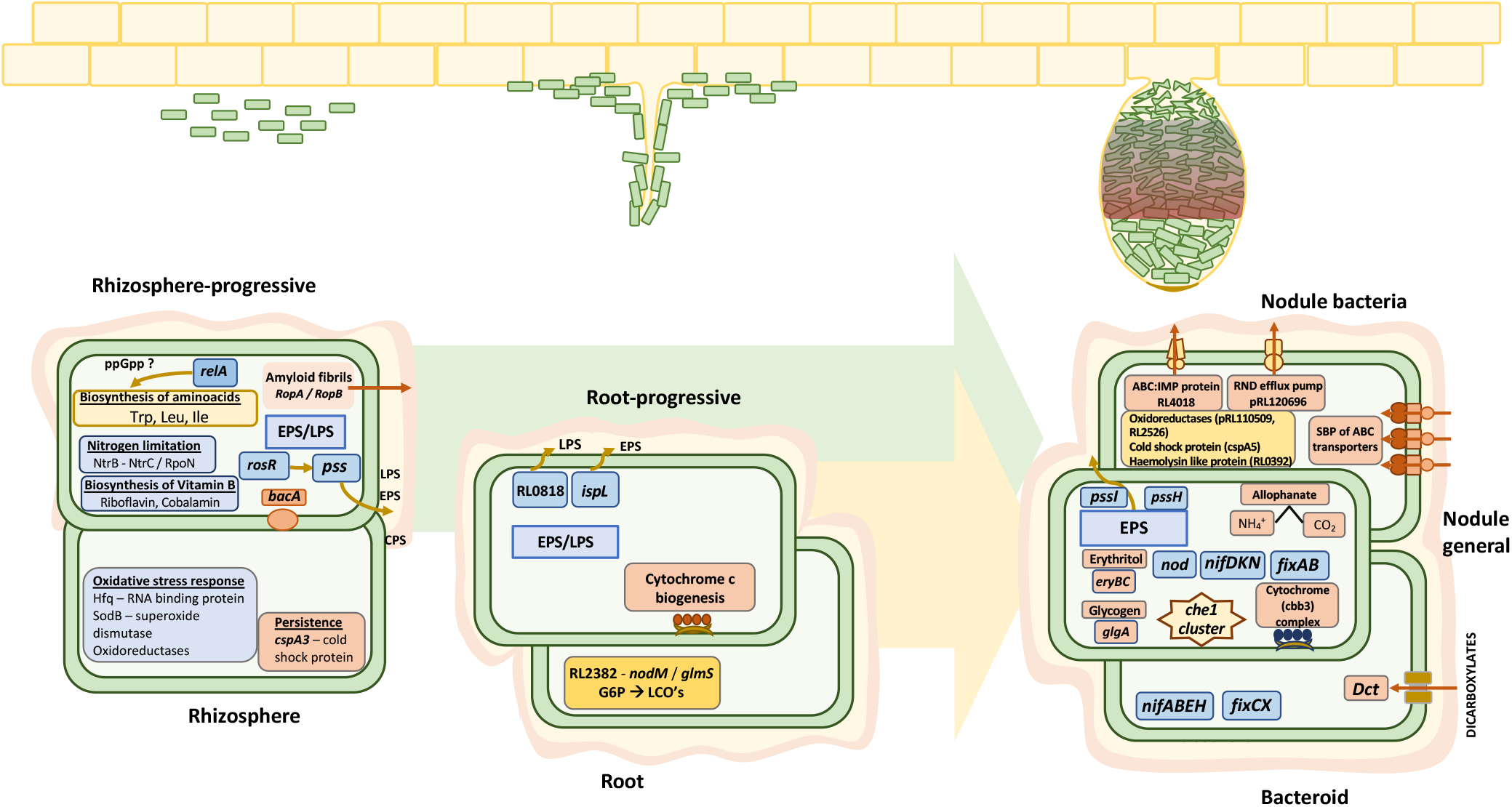
Rlv3841 genes required at different stages of symbiosis with pea from INSeq analysis. Three environments; rhizosphere, root and nodule are shown. Progressive genes alter competition at subsequent stages. Gene requirements at each stage are discussed in the relevant results section, and full gene lists can be found in *SI Appendix*, Tables S1-S8.

## Materials and Methods

### Bacterial and plant growth

Bacterial strains and plasmids shown in *SI Appendix* Table S12 were grown as described previously (58). Mutant strains in *R. leguminosarum* were constructed as described previously (58), except ligation was performed using a BD In-FusionTM cloning kit (Clontech), according to the manufacturer’s instructions (primers listed in *SI Appendix* Table S12). Pea plants (*Pisum sativum* cv. Avola) were grown as described (68).

### INSeq experiments

Construction of the mariner library for sequencing was as described previously (65). Rhizosphere and root colonization samples (5 dpi) were prepared from inoculation of 7-d-old peas with 10^5^ cfu mariner insertion mutants. 10 ml phosphate-buffered saline (PBS) was added to vermiculite, vortexed (1 min) and filtered through 4 layers of muslin. For the root colonized population, the root, snipped from the cotyledons was vortexed (1 min) with 20 ml PBS, ground by pestle and mortar and filtered as above. Both were grown on TY (12 h) and spun to pellet cells before library preparation. For preparation of nodule libraries (28 dpi) from pea seed was inoculated with 10^6^ cfu mariner insertion mutants, a total of ~150,000 nodules were harvested from 1,500 plants and libraries prepared from (i) extract regrown in TY (12 h) before DNA preparation (nodule bacteria), and (ii) DNA extracted from nodules (bacteroids). Libraries were prepared and sequenced as described previously on an Ion Torrent Proton (65). For sequencing data analysis, read quality filtering and mapping (69–70) and conversion of mapped data to wig files (for each replicon) were as described previously (65). Wig files were analyzed by TRANSIT software (71) in two analysis modes (a) whole gene and (b) core region (considering 10-90% of gene, ignoring 10% at 5’- and 3’-ends) with the following parameters; method: HMM (to assign gene status); normalization: trimmed total reads (TTR) and corrected for genome positional bias. Gene status assignment differed by < 0.85% between the two analyses. The 25 gene features that lacked TA sites in their core region were excluded from core region analysis and whole gene analysis was used instead. For disagreements between the two analyses, if a gene was designated ES/DE/AD by whole gene analysis but NE by core region analysis, then final designation was based on the whole gene analysis (i.e. ES/DE/AD). However, when assignment by core region analysis was not NE, then read counts mapped to genes were checked in Integrated Genome Viewer (IGV) (72) and designation was manually assigned.

### Characterization of bacterial mutants

Root colonization was assessed at 5 dpi from inoculation of 7-d-old peas with 10^5^ cfu of bacteria. The root, snipped from the seed, was vortexed (1 min) with PBS (25 ml) before being ground (as above) and resuspended in PBS (10 ml). The rhizosphere population was assessed by adding PBS (30 ml) and vortexing thoroughly. Serial dilution and plating on appropriate antibiotics were used to calculate cfu. Acetylene reduction assays (15) of seed-inoculated peas were performed at 28 dpi, and the same roots, subsequently stained for GusA activity using X-GlcA (73), were photographed to count blue-stained nodules. Competition between two strains to form nodules was assessed at 21 dpi on seed-inoculated peas (total inoculum 10^4^ cfu) by double sequential staining for GusA activity using Magenta-GlcA and CelB activity with X-Gal (73), to count magenta- and blue-stained nodules, respectively. Recovery of bacteria from nodules was assessed by picking the 10 largest nodules from a root, which were surface sterilized (5 min in 2% bleach, followed by 6 washes in sterile distilled water) and crushed in 100 μl PSM buffer (58) in a sterile 1.5-ml Eppendorf with a plastic pestle, before increasing the total volume to 1 ml PSM and vortexing thoroughly. 25 μl was used to inoculate 50 ml of TY broth and OD_600_ followed for 36 h growth, shaking (140 rpm) at 28°C (OD_600_ of 1 = 8 × 10^8^ cfu).

## Supporting information

Data Sheet 1

## Data Availability Statement

The data supporting the findings of the study are available in this article and its SI Appendix.

## Author’s contributions

R.M.W., L.L., B.L.F., H.E.K., S.T.N.A. and V.K.R. made the InSeq libraries, conducted the sequencing, HMM and TRANSIT analysis, made pK19 mutants and all general experimental work. P.S.P., R.M.W. and V.K.R. conceived the project. P.S.P., V.K.R., R.M.W., B.L.F., H.E.K., S.T.N.A., A.K.E. and R.L. interpreted the data and wrote the manuscript.

## Acknowledgments

This work was supported by the Natural Environment Research Council [grant number NE/L501530/1], the Biotechnology and Biological Sciences Research Council [grant numbers BB/M011224/1 and BB/N013387/1] and the Swiss National Science Foundation Postdoc.Mobility to RL [grant number 183901].

