## Supplementary Information for "Lifestyle adaptations of *Rhizobium* from rhizosphere to symbiosis"

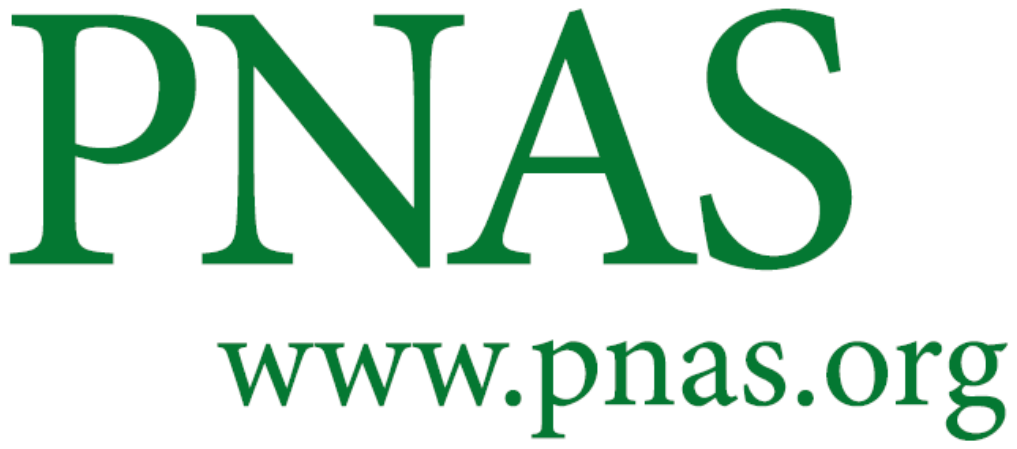

Supplementary Information for

**Lifestyle adaptations of *Rhizobium* from rhizosphere to symbiosis**

**Rachel M. Wheatley, Brandon L. Ford, Li Li, Samuel T. N. Aroney, Hayley E. Knights, Raphael Ledermann, Alison K. East, Vinoy K. Ramachandran**, **and Philip S. Poole**

**Philip S. Poole**

**This PDF file includes:**

Figure S1

Tables S1 to S12

Legends for Datasets S1

SI References

**Other supplementary materials for this manuscript include the following:**

Dataset S1

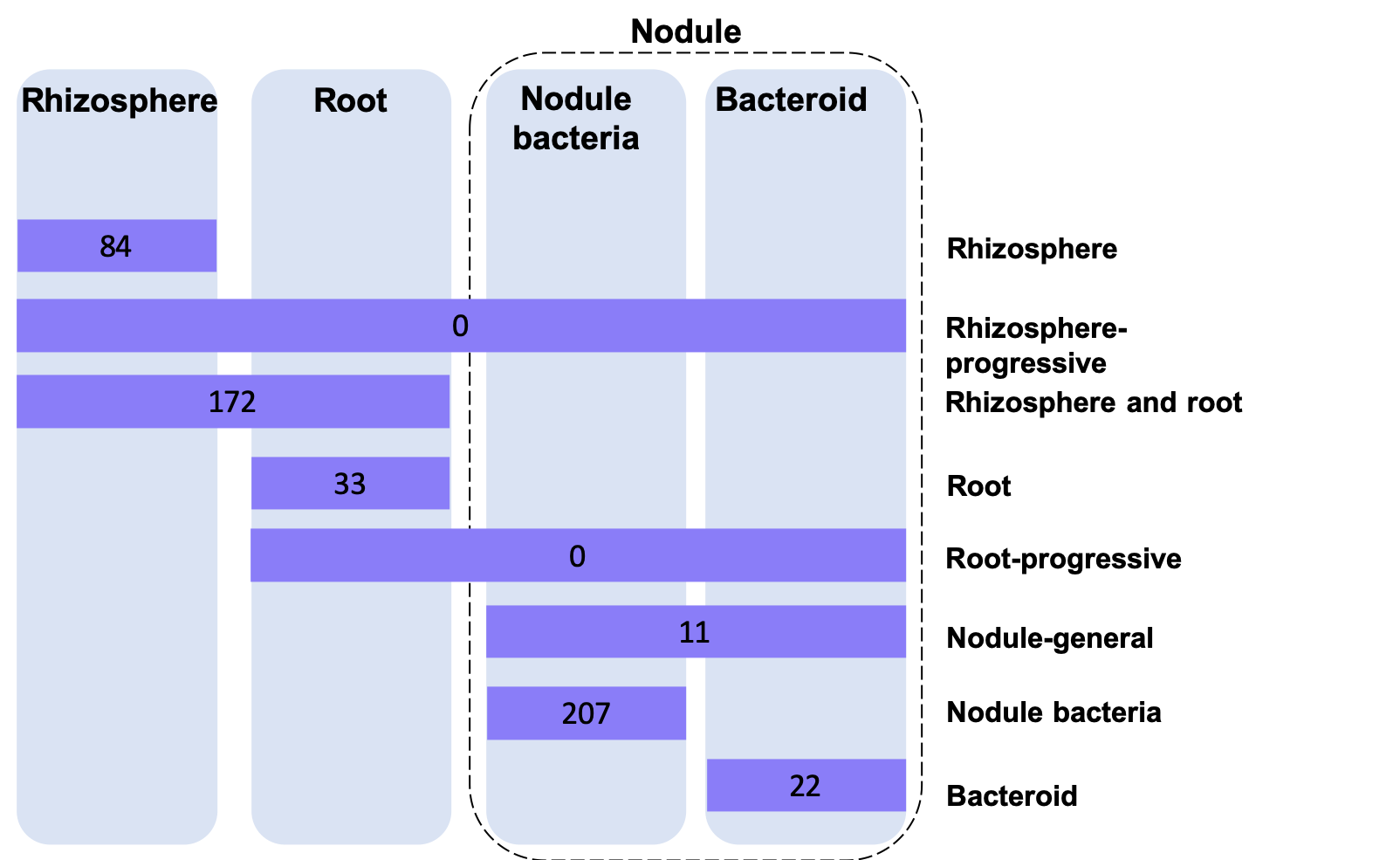

**Fig. S1.** Rlv3841 genetic elements for which mutation confers an advantage during stages of symbiosis with pea. Purple boxes show the number of AD genes at each symbiotic stage, classified as rhizosphere, rhizosphere-progressive, rhizosphere and root, root, root-progressive, nodule-general, nodule bacteria, and bacteroid (*SI Appendix* Table S9). All genes shown were classified NE or DE in the respective input library.

Table S1. Rlv3841 genes (17) classified as rhizosphere-specific.

| **Gene** | **Name** | **Rhizosphere** | **Root colonized** | **Nodule bacteria** | **Bacteroids** | **Description** |
| --- | --- | --- | --- | --- | --- | --- |
| pRL100089 |  | ES | NE | NE | NE | conserved hypothetical protein |
| pRL100144 |  | ES | NE | NE | NE | conserved hypothetical protein |
| pRL100145 |  | ES | NE | NE | NE | putative acyl-CoA dehydrogenase |
| pRL110044 |  | ES | NE | NE | NE | conserved hypothetical protein |
| pRL110210 | *hutC* | DE | NE | NE | NE | putative GntR family transcriptional regulator of histidine utilization operon |
| pRL110539 |  | ES | NE | NE | NE | putative oxidoreductase |
| pRL110564 |  | ES | NE | NE | NE | putative tight adherence protein |
| pRL80090 |  | DE | NE | NE | NE | putative fucose operon protein |
| pRL90136 |  | ES | NE | NE | NE | putative glycosyl transferase |
| pRL90204 |  | ES | NE | NE | NE | putative amidase |
| RL1340 | *sodB* | DE | NE | NE | NE | putative superoxide dismutase |
| RL2284 | *hfq* | ES | NE | NE | NE | putative host factor protein |
| RL2964 | *cspA3* | DE | NE | NE | NE | putative cold shock protein CspA |
| RL3335 |  | ES | NE | NE | NE | putative lysophospholipase |
| RL3474 | *pth* | ES | NE | NE | NE | putative peptidyl-tRNA hydrolase |
| RL4728 |  | ES | NE | NE | NE | conserved hypothetical protein |
| RLt22 |  | DE | NE | NE | NE | tRNA Gln |

HMM classifications ES; essential, DE; growth-defective, NE; neutral.

**Table S2.** Rlv3841 genes (146) classified as rhizosphere-progressive.

| **Gene** | **Name** | **Rhizosphere** | **Root colonized** | **Nodule bacteria** | **Bacteroids** | **Description** |
| --- | --- | --- | --- | --- | --- | --- |
| pRL100386 |  | DE | ES | ES | DE | putative von Willebrand factor type A |
| pRL110389 |  | ES | ES | ES | ES | putative exopolysaccharide production protein |
| pRL110632 | *cobF* | ES | ES | DE | ES | precorrin-6a synthase |
| pRL120618 |  | ES | ES | ES | ES | putative LacI family transcriptional regulator (repressor) |
| pRL70053 |  | DE | DE | DE | DE | putative transmembrane protein |
| pRL90018 | *fixN2* | ES | ES | ES | ES | putative cytochrome oxidase transmembrane component FixN |
| pRL90053 |  | DE | DE | ES | ES | putative O-antigen ligase |
| RL_RF0063 |  | ES | ES | ES | ES | Conserved RNA feature SAM_alpha |
| RL_RF0079 |  | DE | DE | ES | ES | Conserved RNA feature TPP |
| RL_RF0081 |  | DE | ES | ES | ES | Conserved RNA feature SAM_alpha |
| RL0002 |  | ES | ES | ES | ES | putative septum formation protein |
| RL0003 |  | ES | ES | ES | ES | putative shikimate dehydrogenase |
| RL0019 |  | ES | ES | DE | ES | conserved hypothetical protein |
| RL0020 |  | DE | ES | ES | ES | putative N-(5'-phosphoribosyl)anthranilate |
| RL0021 | *trpB* | DE | ES | ES | ES | putative tryptophan synthase beta subunit |
| RL0022 | *trpA* | ES | ES | ES | ES | putative tryptophan synthase alpha subunit |
| RL0026 |  | DE | ES | ES | DE | putative ATP-dependent UvrD family DNA helicase |
| RL0027 |  | DE | ES | ES | DE | conserved hypothetical protein |
| RL0028 |  | DE | ES | ES | DE | putative nucleotidyltransferase protein |
| RL0031 |  | DE | DE | ES | ES | putative S-adenosyl-L-homocysteine hydrolase |
| RL0034 |  | ES | ES | ES | ES | putative HPr kinase |
| RL0052 |  | ES | ES | ES | ES | conserved hypothetical protein |
| RL0108 | *aroA* | DE | DE | ES | ES | putative 3-phosphoshikimate 1-carboxyvinyltransferase (5-enolpyruvylshikimate-3-phosphate synthase) (EPSP synthase) |
| RL0139 |  | DE | DE | ES | ES | putative P-protein |
| RL0179 | *gpmA* | ES | ES | ES | ES | putative 2,3-bisphosphoglycerate-dependent phosphoglycerate mutase |
| RL0180 | *dapB* | ES | ES | ES | ES | putative dihydrodipicolinate reductase |
| RL0254 | *lepA* | DE | DE | ES | ES | putative GTP-binding protein |
| RL0327 |  | ES | ES | ES | ES | putative glutathione S-transferase |
| RL0338 |  | DE | DE | ES | ES | putative glutathione synthetase |
| RL0422 | *rpoN* | ES | DE | ES | DE | putative RNA polymerase sigma-54 factor, involved in nitrogen fixation |
| RL0431 |  | DE | DE | DE | DE | putative plasmid stability protein |
| RL0432 |  | DE | DE | DE | DE | putative plasmid stability protein |
| RL0433 | *fmt* | DE | DE | DE | DE | putative methionyl-tRNA formyltransferase |
| RL0554 | *metZ* | ES | ES | ES | ES | putative O-succinylhomoserine sulfhydrylase |
| RL0588 |  | DE | DE | ES | ES | putative peptidoglycan binding protein |
| RL0751 | *edd* | DE | DE | ES | DE | putative phosphogluconate dehydratase |
| RL0811 |  | DE | DE | ES | ES | putative glycosyltransferase |
| RL0811A |  | DE | DE | ES | ES | putative glycosyltransferase |
| RL0813 |  | DE | DE | ES | ES | putative UDP-glucose-4-epimerase |
| RL0816 |  | DE | DE | ES | ES | conserved hypothetical protein |
| RL0821 |  | DE | DE | ES | ES | putative O-antigen transporter |
| RL0825 | *gmd* | ES | DE | ES | ES | putative GDP-mannose 4,6-dehydratase |
| RL0855 |  | DE | ES | ES | ES | putative gamma-glutamylcysteine synthetase precursor |
| RL0885 |  | DE | ES | ES | DE | putative hydrolase |
| RL0891 |  | ES | ES | DE | DE | putative nitrogen fixation symbiosis related protein |
| RL0920 |  | ES | DE | DE | DE | putative ATP-binding mrp family protein |
| RL0959 |  | ES | ES | ES | ES | putative FAD-dependent oxidoreductase |
| RL1002 | *bioY* | ES | ES | ES | ES | putative transmembrane biotin biosynthesis protein |
| RL1003 |  | ES | ES | ES | ES | putative permease component of ABC transporter Unclass |
| RL1004 |  | ES | ES | ES | ES | putative ATP-binding component of ABC transporter Unclass |
| RL1021 | *ctaC* | DE | DE | ES | ES | putative cytochrome c oxidase polypeptide II precursor |
| RL1022 | *ctaD* | DE | ES | ES | ES | putative cytochrome c oxidase polypeptide I |
| RL1032 | *rnhA* | ES | ES | ES | ES | putative ribonuclease HI |
| RL1105 |  | DE | ES | ES | ES | putative TetR family transcriptional regulator |
| RL1379 | *rosR* | ES | ES | ES | DE | putative MucR/RosR family transcriptional regulator involved in nodulation competitiveness RosR |
| RL1382 |  | ES | ES | ES | ES | putative two-component sensor/regulator; histidine kinase |
| RL1434 | *feuQ* | DE | DE | ES | ES | putative two-component sensor/regulator; histidine kinase |
| RL1503 |  | DE | ES | ES | ES | putative RNA binding protein |
| RL1506 | *relA* | DE | ES | ES | ES | putative stringent response protein |
| RL1523 |  | ES | ES | DE | DE | conserved hypothetical protein |
| RL1524 |  | ES | ES | DE | DE | conserved hypothetical protein |
| RL1525 |  | ES | ES | DE | DE | putative aminomethyltransferase |
| RL1532 |  | ES | ES | ES | DE | putative phosphatidylserine synthase |
| RL1571 | *pepA* | DE | DE | ES | ES | putative aminopeptidase |
| RL1589 |  | ES | DE | ES | DE | putative outer membrane protein RopB |
| RL1597 |  | DE | ES | ES | ES | putative transmembrane protein |
| RL1600 | *ppx* | ES | ES | ES | ES | putative exopolyphosphatase |
| RL1620 | *glyA* | DE | DE | ES | ES | putative serine hydroxymethyltransferase |
| RL1621 | *ribG* | ES | ES | ES | ES | putative riboflavin biosynthesis protein |
| RL1622 | *ribC* | ES | ES | ES | ES | putative riboflavin biosynthesis protein |
| RL1738 | *pyrC* | DE | DE | DE | DE | putative dihydroorotase |
| RL1739 | *pyrB* | DE | DE | DE | DE | putative aspartate carbamoyltransferase |
| RL1803 | *ilvD* | ES | ES | ES | ES | putative dihydroxy-acid dehydratase |
| RL1945 |  | DE | DE | ES | ES | putative vitamin B12-dependent ribonucleotide reductase |
| RL1946 |  | ES | ES | ES | ES | hypothetical protein |
| RL2052 |  | ES | DE | ES | ES | putative peptidase |
| RL2234 | *icdB* | DE | ES | ES | ES | putative citrate synthase |
| RL2239 | *eno* | ES | DE | ES | ES | putative enolase |
| RL2255 | *dus* | ES | DE | ES | DE | putative tRNA-dihydrouridine synthase (nitrogen regulation protein) |
| RL2256 | *ntrB* | DE | DE | ES | DE | putative two-component sensor/regulator; histidine kinase NtrB |
| RL2257 | *ntrC* | DE | DE | ES | DE | two-component sensor/regulator; nitrogen transcriptional regulator NtrC |
| RL2303 | *ccdA* | DE | ES | ES | ES | putative cytochrome c-type biogenesis protein |
| RL2492 | *ppiD* | DE | ES | ES | ES | putative peptidyl-prolyl cis-trans isomerase D |
| RL2493 | *trpD* | DE | DE | ES | ES | putative anthranilate phosphoribosyltransferase |
| RL2494 | *trpC* | DE | DE | ES | ES | putative indole-3-glycerol phosphate synthase |
| RL2567 | *rLuC* | DE | ES | DE | ES | putative ribosomal large subunit pseudouridine synthase C |
| RL2594 | *deaD* | DE | ES | ES | ES | putative cold-shock DEAD-box protein A |
| RL2598 | *rpe* | ES | ES | ES | ES | putative ribulose phosphate 3-epimerase |
| RL2601 | *purB* | ES | ES | ES | ES | putative adenylosuccinate lyase |
| RL2694 | *gor* | DE | ES | ES | ES | putative glutathione reductase |
| RL2775 | *ropA* | DE | DE | ES | DE | putative outer membrane porin protein RopA |
| RL2781 | *cobU* | ES | ES | ES | ES | putative nicotinate-nucleotide--dimethylbenzimidazole phosphoribosyltransferase |
| RL2781A | *cobV* | ES | ES | ES | ES | putative cobalamin synthase |
| RL2829 | *cobO* | ES | ES | ES | ES | putative cob(I)yrinic acid a,c-diamide adenosyltransferase |
| RL3205 | *ilvC* | ES | ES | ES | DE | putative ketol-acid reductoisomerase |
| RL3206 |  | ES | ES | ES | DE | putative TetR family transcriptional regulator |
| RL3245 | *ilvI* | ES | ES | ES | ES | putative acetolactate synthase isozyme III large subunit |
| RL3422 | *greA* | ES | ES | ES | ES | putative transcription elongation factor |
| RL3423 |  | ES | ES | ES | ES | putative lipopolysaccharide core biosynthesis protein |
| RL3454 |  | ES | ES | DE | ES | putative two-component sensor/regulator; transcriptional regulator |
| RL3486 | *petA* | ES | ES | ES | ES | putative ubiquinol-cytochrome c reductase iron-sulfur subunit |
| RL3500 |  | DE | ES | DE | DE | conserved hypothetical protein |
| RL3501 |  | ES | DE | DE | DE | putative transmembrane protein |
| RL3507 |  | ES | ES | ES | ES | putative phosphoesterase |
| RL3513 | *leuA* | ES | ES | ES | ES | putative 2-isopropylmalate synthase |
| RL3521 | *trpE* | DE | DE | ES | ES | putative anthralinate synthase |
| RL3557 | *bacA* | ES | ES | ES | ES | putative transmembrane transporter protein required for bacteroid development |
| RL3651 |  | ES | ES | ES | ES | putative glycosyl transferase |
| RL3664 | *pssN* | ES | ES | ES | ES | putative capsule polysaccharide export protein |
| RL3667 |  | ES | DE | ES | DE | putative UDP-glucose 6-dehydrogenase |
| RL3674 | *noeK* | ES | ES | ES | ES | putative phosphomannomutase |
| RL3765 | *rLuD* | ES | ES | ES | ES | putative ribosomal large subunit pseudouridine synthase D |
| RL3768 | *purA* | ES | ES | ES | ES | putative adenylosuccinate synthetase |
| RL3988 |  | DE | DE | DE | DE | conserved hypothetical protein |
| RL3989 | *ruvA* | DE | DE | DE | DE | putative Holliday junction DNA helicase RuvA |
| RL3990 | *ruvB* | DE | DE | DE | DE | putative Holliday junction DNA helicase RuvB |
| RL4006 | *cbbT* | ES | ES | ES | ES | putative transketolase |
| RL4066 |  | DE | DE | ES | ES | conserved hypothetical protein |
| RL4131 |  | ES | ES | ES | ES | putative TPR repeat family protein |
| RL4283 | *ptsP* | ES | ES | ES | ES | putative phosphoenolpyruvate phosphotransferase |
| RL4291 |  | ES | ES | ES | ES | conserved hypothetical protein |
| RL4296 | *argJ* | ES | DE | ES | ES | putative arginine biosynthesis bifunctional protein |
| RL4330 | *ftsE* | DE | ES | ES | ES | putative cell division ATP-binding protein |
| RL4331 | *ftsX* | DE | ES | ES | ES | putative cell division protein homolog |
| RL4337 | *tyrC* | DE | ES | ES | ES | putative prephenate dehydrogenase |
| RL4338 | *hisC* | DE | ES | ES | ES | putative histidinol-phosphate aminotransferase |
| RL4348 | *cobT* | DE | ES | ES | ES | putative aerobic cobaltochelatase |
| RL4349 | *cobS* | ES | ES | ES | ES | putative aerobic cobaltochelatase |
| RL4399 |  | DE | DE | ES | ES | conserved hypothetical protein YKOF super family |
| RL4400 |  | DE | DE | ES | ES | putative permease component of ABC transporter NitT |
| RL4401 |  | DE | DE | ES | ES | putative ATP-binding component of ABC transporter NitT |
| RL4402 |  | DE | DE | ES | ES | putative SBP of ABC transporter NitT |
| RL4440 |  | DE | DE | ES | ES | conserved hypothetical protein |
| RL4506 | *typA* | DE | DE | ES | ES | putative GTP-binding protein TypA/BipA (tyrosine phosphorylated protein A) |
| RL4537 | *ccmA* | ES | DE | ES | ES | putative ATP-binding component of ABC transporter Export cytochrome c biogenesis |
| RL4544 |  | ES | ES | ES | ES | putative methylase |
| RL4559 |  | ES | ES | ES | ES | putative two-component sensor/regulator; transcriptional regulator |
| RL4583 | *fbpB* | ES | DE | DE | DE | putative permease component of ABC transporter Unclass ferric cations transporter |
| RL4606 | *metA* | ES | ES | ES | ES | putative homoserine O-succinyltransferase |
| RL4691 |  | DE | ES | ES | ES | putative peptidase |
| RL4692 | *ctpA* | DE | ES | ES | ES | putative carboxy-terminal processing protease precursor |
| RL4707 | *leuB* | DE | ES | DE | ES | putative 3-isopropylmalate dehydrogenase |
| RL4716 |  | ES | DE | ES | ES | conserved hypothetical exported protein |
| RL4731 |  | DE | DE | ES | ES | putative racemase |
| RL4732 | *leuS* | DE | DE | ES | ES | putative leucyl-tRNA synthetase |
| RLt11 |  | DE | DE | ES | ES | tRNA Ala |

HMM classifications ES; essential, DE; growth-defective, NE; neutral.

**Table S3.** Rlv3841 genes (7) classified as rhizosphere and root-specific.

| **Gene** | **Name** | **Rhizosphere** | **Root colonized** | **Nodule bacteria** | **Bacteroids** | **Description** |
| --- | --- | --- | --- | --- | --- | --- |
| pRL110107 |  | ES | ES | NE | NE | conserved hypothetical exported protein |
| RL2615 |  | ES | ES | NE | NE | putative glutaredoxin |
| RL2776 |  | ES | ES | NE | NE | putative AsnC family transcriptional regulator |
| RL3005 |  | ES | ES | NE | NE | hypothetical protein |
| RL4065 |  | DE | DE | NE | NE | conserved hypothetical protein |
| RL4162 | *eda* | ES | ES | NE | NE | putative 2-dehydro-3-deoxyphosphogluconate aldolase |
| RLt42 |  | ES | ES | NE | NE | tRNA Met |

HMM classifications ES; essential, DE; growth-defective, NE; neutral.

**Table S4.** Rlv3841 genes (23) classified as root-specific.

| **Gene** | **Name** | **Rhizosphere** | **Root colonized** | **Nodule bacteria** | **Bacteroids** | **Description** |
| --- | --- | --- | --- | --- | --- | --- |
| pRL100162A |  | NE | DE | NE | NE | hypothetical protein |
| pRL100242 |  | NE | ES | NE | NE | conserved hypothetical protein |
| pRL120021 |  | NE | ES | NE | NE | hypothetical protein |
| pRL70001 | *repA* | NE | DE | NE | NE | putative replication protein RepA |
| pRL70002 | *repB* | NE | DE | NE | NE | putative replication protein RepB |
| pRL70003 | *repC* | NE | DE | NE | NE | putative replication protein RepC |
| pRL70004 |  | NE | DE | NE | NE | conserved hypothetical protein |
| pRL70005 |  | NE | DE | NE | NE | conserved hypothetical protein |
| pRL70006 |  | NE | DE | NE | NE | hypothetical protein |
| pRL70007 |  | NE | DE | NE | NE | putative plasmid stabilisation protein |
| pRL70028 |  | NE | DE | NE | NE | conserved hypothetical protein |
| pRL90052 |  | NE | ES | NE | NE | putative polysaccharide biosynthesis O-antigen related protein |
| RL0137 |  | NE | ES | NE | NE | putative XRE family transcriptional regulator |
| RL0138 |  | NE | DE | NE | NE | putative glyoxalase/dioxygenase |
| RL1040 |  | NE | ES | NE | NE | putative LysR family transcriptional regulator |
| RL1371 |  | NE | ES | NE | NE | putative transmembrane protein |
| RL2512 |  | NE | DE | NE | NE | putative transmembrane protein SecG |
| RL3663 | *pssO* | NE | ES | NE | NE | putative polysaccharide transport outer membrane protein |
| RL4147 |  | NE | ES | NE | NE | conserved hypothetical protein |
| RL4148 |  | NE | ES | NE | NE | putative cobalamin synthesis protein |
| RL4322 | *tlpA* | NE | ES | NE | NE | putative thiol:disulfide interchange protein |
| RL4617 | *hslR* | NE | DE | NE | NE | putative heat shock protein 15 |
| RL4618 |  | NE | DE | NE | NE | conserved hypothetical protein |

HMM classifications ES; essential, DE; growth-defective, NE; neutral.

**Table S5**. Rlv3841 genes (33) classified as root-progressive.

| **Gene** | **Name** | **Rhizosphere** | **Root colonized** | **Nodule bacteria** | **Bacteroids** | **Description** |
| --- | --- | --- | --- | --- | --- | --- |
| pRL100053 |  | NE | ES | DE | DE | putative transmembrane protein |
| pRL70027 |  | NE | DE | DE | DE | hypothetical protein |
| pRL70055 |  | NE | DE | ES | DE | hypothetical protein |
| pRL70056 |  | NE | DE | ES | DE | hypothetical protein |
| pRL70108 |  | NE | ES | ES | ES | hypothetical protein |
| pRL80032* |  | NE | ES/DE | ES/DE | ES/DE | putative LysR family transcriptional regulator |
| RL0377 | *hemN* | NE | DE | ES | DE | putative oxygen-independent coprophorphyrinogen III oxidase |
| RL0378 |  | NE | DE | ES | DE | putative HAM1 family protein |
| RL0379 |  | NE | DE | ES | DE | putative glyoxalase |
| RL0380 | *rph* | NE | DE | ES | DE | putative ribonuclease |
| RL0434 | *truA* | NE | DE | DE | DE | putative tRNA pseudouridine synthase A |
| RL0814 |  | NE | DE | ES | ES | conserved hypothetical protein |
| RL0815 |  | NE | DE | ES | ES | putative acetyltransferase |
| RL0818 |  | NE | DE | ES | ES | putative lipopolysaccharide biosynthesis protein |
| RL0820 | *hisH2* | NE | DE | ES | ES | putative imidazole glycerol phosphate synthase subunit (igp synthase glutamine amidotransferase subunit) |
| RL0822 |  | NE | DE | ES | ES | putative DegT family aminotransferase |
| RL0847 | *guaB* | NE | DE | ES | ES | putative inosine-5'-monophosphate dehydrogenase |
| RL0921 |  | NE | DE | DE | DE | putative cationic transport protein, CorA family |
| RL0957 |  | NE | ES | ES | ES | conserved hypothetical protein |
| RL1478 | *amn* | NE | DE | ES | ES | putative AMP nucleosidase |
| RL1553 |  | NE | ES | DE | DE | putative transmembrane protein |
| RL1572 |  | NE | DE | ES | ES | putative DNA polymerase III subunit |
| RL2247 |  | NE | ES | ES | ES | putative transmembrane transglycosylase-associated protein, integral membrane protein |
| RL2248 |  | NE | ES | ES | ES | putative transmembrane transglycosylase-associated protein, integral membrane protein |
| RL2382 | *nodM* | NE | DE | ES | ES | putative glucosamine--fructose-6-phosphate aminotransferase [isomerizing] |
| RL3668 |  | NE | DE | ES | DE | putative serine/threonine protein phosphatase |
| RL3677 | *lspL* | NE | ES | ES | ES | putative UDP-glucuronate 5'-epimerase |
| RL3987 |  | NE | DE | DE | DE | conserved hypothetical protein |
| RL4007 | *gap* | NE | ES | ES | ES | putative glyceraldehyde-3-phosphate dehydrogenase |
| RL4538 | *ccmB* | NE | DE | ES | ES | putative permease component of ABC transporter Export cytochrome c binding export protein |
| RL4539 | *cycZ* | NE | DE | ES | ES | putative permease component of ABC transporter Export heme exporter protein c (cytochrome c-type biogenesis protein) |
| RL4540 | *cycY* | NE | DE | ES | ES | putative thiol:disulfide interchange protein CycY precursor (cytochrome c biogenesis protein) |
| RL4555 |  | NE | DE | ES | ES | putative 3-isopropylmalate dehydratase |

HMM classifications ES; essential, DE; growth-defective, NE; neutral.

*manual curation

**Table S6**. Rlv3841 genes (211) classified as nodule-general.

| **Gene** | **Name** | **Rhizosphere** | **Root colonized** | **Nodule bacteria** | **Bacteroids** | **Description** |
| --- | --- | --- | --- | --- | --- | --- |
| pRL100027 |  | NE | NE | ES | ES | putative restriction modification methylase |
| pRL100051 |  | NE | NE | DE | DE | hypothetical protein |
| pRL100052 | *cspA* | NE | NE | DE | DE | putative cold shock protein CspA |
| pRL100052A |  | NE | NE | DE | DE | pseudogene |
| pRL100132 |  | NE | NE | DE | DE | hypothetical protein |
| pRL100133 |  | NE | NE | DE | DE | putative IclR family transcriptional regulator |
| pRL100134 | *gabD* | NE | NE | DE | DE | orthologue of aatK (blcA) succinate semialdehyde dehydrogenase dehydrogenase gene of A. tumefaciens |
| pRL100158 | *nifN* | NE | NE | DE | DE | putative nitrogenase iron-molybdenum cofactor biosynthesis protein NifN |
| pRL100160 | *nifK* | NE | NE | ES | DE | nitrogenase molybdenum-iron protein beta chain NifK |
| pRL100161 | *nifD* | NE | NE | ES | DE | nitrogenase molybdenum-iron protein alpha chain NifD |
| pRL100167 |  | NE | NE | ES | ES | putative transposase part of insertion sequence |
| pRL100181 | *nodL* | NE | NE | ES | ES | putative nodulation protein NodL |
| pRL100182 | *nodE* | NE | NE | DE | DE | nodulation protein NodE homology with beta-ketoacyl synthases e.g.FabB |
| pRL100183 | *nodF* | NE | NE | DE | DE | nodulation protein NodF acyl carrier protein (ACP) used in Nod factor synthesis |
| pRL100184 | *nodD* | NE | NE | DE | DE | nodulation protein NodD |
| pRL100185 | *nodA* | NE | NE | DE | DE | nodulation protein NodA |
| pRL100186 | *nodB* | NE | NE | DE | DE | putative chitooligosaccharide deacetylase NodB |
| pRL100187 | *nodC* | NE | NE | DE | DE | N-acetylglucosaminyltransferase NodC |
| pRL100188 | *nodI* | NE | NE | DE | DE | putative ATP-binding component of ABC transporter Export of nod factor |
| pRL100189 | *nodJ* | NE | NE | DE | DE | nodulation protein NodJ involved in Nod factor export |
| pRL100199 | *fixB* | NE | NE | DE | DE | electron transfer protein FixB |
| pRL100200 | *fixA* | NE | NE | DE | DE | electron transfer protein FixA |
| pRL100205 | *fixN1* | NE | NE | DE | DE | putative transmembrane cytochrome oxidase subunit |
| pRL100211A |  | NE | NE | DE | DE | putative GntR family transcriptional regulator |
| pRL100212 |  | NE | NE | DE | DE | putative GntR family transcriptional regulator |
| pRL100246 |  | NE | NE | ES | ES | putative aldehyde dehydrogenase |
| pRL100247 |  | NE | NE | ES | ES | conserved hypothetical protein |
| pRL100433 |  | NE | NE | ES | ES | putative allophanate hydrolase subunit 2 |
| pRL100434 |  | NE | NE | ES | ES | putative allophanate hydrolase subunit 1 |
| pRL100435 |  | NE | NE | ES | ES | conserved hypothetical protein |
| pRL100470 |  | NE | NE | ES | ES | hypothetical protein |
| pRL110047 |  | NE | NE | ES | DE | putative permease component of ABC transporter Export |
| pRL110049 |  | NE | NE | ES | ES | putative glycosyltransferase |
| pRL110056 |  | NE | NE | DE | ES | putative O-antigen related protein |
| pRL110415 | *rhaD* | NE | NE | DE | DE | putative short-chain dehydrogenase/oxidoreductase involved in competition for nodulation |
| pRL120205 | *eryB* | NE | NE | ES | DE | putative erythritol phosphate dehydrogenase |
| pRL120206 | *eryC* | NE | NE | ES | DE | putative erythrulose 4-phosphate dehydrogenase |
| pRL120291 |  | NE | NE | DE | DE | putative MarR family transcriptional regulator |
| pRL120292 |  | NE | NE | DE | DE | putative short-chain dehydrogenase/oxidoreductase |
| pRL120397 |  | NE | NE | ES | ES | putative GntR family transcriptional regulator |
| pRL120518 |  | NE | NE | ES | ES | putative TetR family transcriptional regulator |
| pRL120694 |  | NE | NE | DE | DE | putative LacI family transcriptional regulator (repressor) |
| pRL120695 |  | NE | NE | DE | DE | putative TetR family transcriptional regulator of RND family efflux transporter |
| pRL70051 |  | NE | NE | DE | DE | pseudogene |
| pRL70166 |  | NE | NE | DE | DE | conserved hypothetical protein |
| pRL80079 |  | NE | NE | DE | DE | putative DeoR family transcriptional regulator (repressor) |
| pRL80080 |  | NE | NE | DE | DE | putative fructokinase |
| pRL90058 |  | NE | NE | ES | ES | putative TetR family transcriptional regulator of RND family efflux transporter (regulator of rmrAB. Homologous with region in R. etli) |
| pRL90141 |  | NE | NE | DE | DE | hypothetical protein |
| pRL90143 |  | NE | NE | DE | DE | putative transposase |
| pRL90262 |  | NE | NE | ES | ES | putative ATP-binding component of ABC transporter HAAT |
| pRL90263 |  | NE | NE | ES | ES | conserved hypothetical protein |
| pRL90320 |  | NE | NE | DE | DE | hypothetical protein |
| RL0025 |  | NE | NE | ES | DE | putative thioredoxin |
| RL0032 |  | NE | NE | ES | ES | putative phosphocarrier protein HPr for mannose |
| RL0033 |  | NE | NE | ES | ES | putative phosphotransferase system component, mannose PTS component IIA |
| RL0123 | *truB* | NE | NE | ES | ES | putative tRNA pseudouridine synthase B (tRNA pseudouridine 55 synthase) (Psi55 synthase) (pseudouridylate synthase) (uracil hydrolyase) |
| RL0124 | *rbfA* | NE | NE | ES | ES | putative ribosome-binding factor protein |
| RL0133 |  | NE | NE | ES | ES | conserved hypothetical protein |
| RL0143 |  | NE | NE | ES | DE | putative PfkB family sugar kinase |
| RL0148 | *recF* | NE | NE | DE | DE | putative DNA replication and repair protein |
| RL0149 |  | NE | NE | DE | DE | putative XRE family transcriptional regulator, up reg in RL0390 (PraR) mutant MA |
| RL0186 |  | NE | NE | ES | ES | putative permease component of ABC transporter PepT (S. mel SBP homologue SMc02832 induced by taurine, valine, isoleucine) |
| RL0187 |  | NE | NE | ES | ES | putative permease component of ABC transporter PepT (S. mel SBP homologue SMc02832 induced by taurine, valine, isoleucine) |
| RL0226 |  | NE | NE | ES | ES | putative permease component of ABC transporter PepT |
| RL0228 |  | NE | NE | ES | ES | putative SBP of ABC transporter PepT |
| RL0344 |  | NE | NE | ES | ES | putative AsnC family transcriptional regulator |
| RL0397 | *fur* | NE | NE | DE | DE | putative FUR-like transcriptional regulator, iron response regulator |
| RL0398 |  | NE | NE | DE | DE | putative acetyltransferase |
| RL0403 |  | NE | NE | ES | DE | conserved hypothetical protein |
| RL0424 |  | NE | NE | DE | DE | putative sigma-54 modulation protein (has ribosome-associated inhibitory protein (raiA) domain) |
| RL0425 | *ptsN* | NE | NE | DE | DE | putative nitrogen regulatory IIA protein |
| RL0493 | *pyrC* | NE | NE | ES | DE | putative dihydroorotase |
| RL0502 | *frk* | NE | NE | ES | DE | fructokinase |
| RL0546 | *phoU* | NE | NE | ES | ES | putative phosphate uptake regulator PhoU, unknown mechanism to regulate expression of high-affinity ABC systems |
| RL0571 |  | NE | NE | ES | ES | conserved hypothetical protein |
| RL0572 |  | NE | NE | ES | ES | putative dihydroorotate dehydrogenase |
| RL0652 |  | NE | NE | DE | DE | putative LacI family transcriptional regulator (repressor) |
| RL0685 |  | NE | NE | ES | ES | putative chemoreceptor protein |
| RL0686 | *cheX* | NE | NE | ES | ES | putative chemotaxis related CheX protein |
| RL0687 | *cheY* | NE | NE | ES | ES | putative two-component sensor/regulator; chemotaxis transcriptional regulator CheY |
| RL0688 | *cheA* | NE | NE | ES | ES | putative chemotaxis protein CheA |
| RL0689 | *cheW* | NE | NE | ES | ES | putative chemotaxis protein CheW family |
| RL0690 | *cheR* | NE | NE | ES | ES | putative chemotaxis protein methyltransferase |
| RL0691 | *cheB* | NE | NE | ES | ES | putative chemotaxis response regulator protein-glutamate methylesterase |
| RL0692 | *cheY* | NE | NE | ES | ES | putative chemotaxis protein |
| RL0693 | *cheD* | NE | NE | ES | ES | putative chemotaxis related protein |
| RL0695* | *fliF* | NE | NE | DE | ES/DE | putative flagellar M-ring protein |
| RL0699 | *flhB* | NE | NE | ES | ES | putative flagellar biosynthetic protein |
| RL0703* | *motA* | NE | NE | ES/DE | DE | putative chemotaxis motility protein |
| RL0704* | *flgF* | NE | NE | ES/DE | DE | putative flagella-associated protein FlgF |
| RL0712 | *flgI* | NE | NE | DE | DE | putative flagellar basal body P-ring protein precursor FlgI |
| RL0713 |  | NE | NE | DE | DE | putative exported flagella-related protein |
| RL0714 | *flgH* | NE | NE | DE | DE | putative flagellar basal body L-ring protein precursor FlgH |
| RL0723* | *motB* | NE | NE | DE | ES/DE | putative motility protein MotB |
| RL0724* | *motC* | NE | NE | DE | ES/DE | putative motility protein precursor MotC |
| RL0726 |  | NE | NE | DE | DE | conserved hypothetical exported protein |
| RL0727 |  | NE | NE | DE | DE | putative two-component sensor/regulator; transcriptional regulator |
| RL0730* | *flgL* | NE | NE | ES/DE | DE | putative flagellar hook-associated protein FlgK |
| RL0731 | *flaF* | NE | NE | DE | DE | putative flagellar synthesis related protein FlaF |
| RL0732 | *flbT* | NE | NE | DE | DE | putative flagellar biosynthesis repressor protein, binds to and promotes degradation of flagellar mRNA (post-transcriptional regulation) |
| RL0733* | *flDE* | NE | NE | DE | ES/DE | putative flagellar hook formation protein FlgD |
| RL0735* | flhA | NE | NE | DE | ES/DE | putative flagellar biosynthesis protein FlhA |
| RL0776 | *iolA* | NE | NE | ES | DE | malonic semialdehyde oxidative decarboxylase |
| RL0817 |  | NE | NE | ES | DE | putative transmembrane protein |
| RL0819 |  | NE | NE | ES | ES | putative imidazole glycerol phosphate synthase subunit |
| RL0824 |  | NE | NE | ES | ES | putative oxidoreductase |
| RL0826 | *fcl* | NE | NE | ES | ES | putative GDP-L-fucose synthetase |
| RL0892 |  | NE | NE | DE | DE | putative ribosomal large subunit pseudouridine synthase B |
| RL0903 | *gph* | NE | NE | ES | ES | putative phosphoglycolate phosphatase |
| RL0934 | *moaB* | NE | NE | ES | ES | putative molybdenum cofactor biosynthesis protein B |
| RL0938 |  | NE | NE | ES | ES | conserved hypothetical protein |
| RL0939 |  | NE | NE | ES | ES | conserved hypothetical protein |
| RL0947 | *purD* | NE | NE | ES | ES | putative phosphoribosylamine--glycine ligase |
| RL0962 |  | NE | NE | ES | ES | putative ring hydroxylating dioxygenase subunit |
| RL0994 |  | NE | NE | ES | DE | putative hydroxypyruvate reductase |
| RL0995 |  | NE | NE | ES | DE | putative tartrate dehydrogenase |
| RL1383 |  | NE | NE | ES | ES | putative peptidoglycan binding transmembrane protein |
| RL1436 | *cycH* | NE | NE | ES | ES | putative cytochrome c-type biogenesis protein |
| RL1437 | *cycJ* | NE | NE | ES | ES | putative cytochrome c-type biogenesis protein |
| RL1438 | *cycK* | NE | NE | ES | ES | putative cytochrome c-type biogenesis protein |
| RL1439 | *cycL* | NE | NE | ES | ES | putative cytochrome c-type biogenesis protein |
| RL1440 | *degP* | NE | NE | ES | ES | putative serine protease |
| RL1470 |  | NE | NE | DE | ES | putative glycosyl transferase |
| RL1499 | *ropA* | NE | NE | ES | ES | putative outer membrane porin protein RopA |
| RL1508 |  | NE | NE | DE | ES | putative transmembrane protein |
| RL1528 |  | NE | NE | DE | DE | conserved hypothetical protein |
| RL1545 |  | NE | NE | ES | ES | putative short-chain dehydrogenase/reductase |
| RL1546 | *purF* | NE | NE | ES | ES | putative amidophosphoribosyltransferase |
| RL1552 | *rplI* | NE | NE | DE | DE | putative 50S ribosomal protein L9 |
| RL1595 | *purN* | NE | NE | ES | ES | putative 5'-phosphoribosylglycinamide formyltransferase |
| RL1596 | *purM* | NE | NE | ES | ES | putative phosphoribosylformylglycinamidine cyclo-ligase |
| RL1617 |  | NE | NE | DE | DE | putative transmembrane protein |
| RL1618 |  | NE | NE | DE | DE | putative MarR family transcriptional regulator |
| RL1637 |  | NE | NE | DE | DE | conserved hypothetical protein |
| RL1640 | *ihfA* | NE | NE | DE | DE | putative integration host factor alpha-subunit |
| RL1641 |  | NE | NE | DE | DE | putative MerR family transcriptional regulator |
| RL1731 | *rpmG* | NE | NE | DE | DE | putative 50S ribosomal protein L33 |
| RL1732 |  | NE | NE | DE | DE | putative transmembrane protein |
| RL1737 |  | NE | NE | DE | DE | putative transmembrane protein |
| RL2045 | *scpB* | NE | NE | ES | ES | putative chromosome segregation and condensation protein B |
| RL2117A |  | NE | NE | ES | DE | hypothetical protein |
| RL2152 |  | NE | NE | ES | ES | hypothetical protein |
| RL2212 | *clpS* | NE | NE | DE | DE | putative ATP-dependent Clp protease adaptor protein |
| RL2288 | *cysG2* | NE | NE | ES | ES | putative siroheme synthase |
| RL2289 |  | NE | NE | ES | ES | conserved hypothetical protein |
| RL2290 | *cysI* | NE | NE | ES | ES | putative sulfite reductase |
| RL2291 |  | NE | NE | ES | ES | conserved hypothetical protein |
| RL2307 |  | NE | NE | DE | DE | conserved hypothetical protein |
| RL2404 |  | NE | NE | DE | DE | putative peptidyl-prolyl cis-trans isomerase (cyclophilin) |
| RL2405 |  | NE | NE | ES | DE | putative peptidyl-prolyl cis-trans isomerase B (cyclophilin-related protein) |
| RL2406 | *queA* | NE | NE | ES | DE | putative S-adenosylmethionine:tRNA ribosyltransferase-isomerase |
| RL2407 | *tgt* | NE | NE | ES | DE | putative queuine tRNA-ribosyltransferase |
| RL2530 |  | NE | NE | DE | ES | putative nitrogen fixation protein |
| RL2568 |  | NE | NE | DE | ES | putative CrcB family transmembrane protein |
| RL2588 | *tyrS* | NE | NE | ES | ES | putative tyrosyl-tRNA synthetase |
| RL2606 | *purC* | NE | NE | ES | ES | putative phosphoribosylaminoimidazole-succinocarboxamide synthase (saicar synthetase) |
| RL2611 |  | NE | NE | ES | ES | hypothetical protein |
| RL2612 | *purL* | NE | NE | ES | ES | putative phosphoribosylformylglycinamidine synthase II |
| RL2637 | *recA* | NE | NE | ES | ES | putative recombinase |
| RL2828 |  | NE | NE | ES | ES | putative XRE family (HipB) family transcriptional regulator |
| RL2924 |  | NE | NE | ES | ES | putative MarR family transcriptional regulator |
| RL3244 | *ilvH* | NE | NE | ES | ES | putative acetolactate synthase isozyme III |
| RL3259 |  | NE | NE | ES | ES | conserved hypothetical protein |
| RL3260 |  | NE | NE | ES | ES | conserved hypothetical protein |
| RL3333 |  | NE | NE | DE | ES | putative ATP-binding:ATP-binding (ABC:ABC) component of ABC transporter Unclass |
| RL3439 | *lpcB* | NE | NE | DE | ES | putative CMP KDO transferase |
| RL3440 | *gspA* | NE | NE | DE | ES | putative general stress protein A |
| RL3453 |  | NE | NE | DE | ES | putative two-component sensor/regulator; histidine kinase |
| RL3459 | *csaA* | NE | NE | ES | DE | putative chaperone protein |
| RL3460 | *proC* | NE | NE | ES | DE | putative pyrroline-5-carboxylate reductase |
| RL3465 |  | NE | NE | DE | DE | conserved hypothetical protein |
| RL3485 | *petB* | NE | NE | ES | ES | putative cytochrome b |
| RL3539 |  | NE | NE | ES | ES | putative metal-dependent phosphohydrolases with conserved 'HD' motif |
| RL3549 | *glnII* | NE | NE | DE | ES | putative glutamine synthetase II |
| RL3649 | *pssI* | NE | NE | ES | ES | putative glycosyltransferase |
| RL3650 | *pssH* | NE | NE | DE | ES | putative glycosyltransferase |
| RL3764 |  | NE | NE | ES | ES | putative transmembrane protein |
| RL3820 |  | NE | NE | ES | ES | putative exopolysaccharide production protein |
| RL3948 |  | NE | NE | ES | ES | conserved hypothetical protein |
| RL4040 | *thiE* | NE | NE | ES | ES | putative thiamine-phosphate pyrophosphorylase |
| RL4043 |  | NE | NE | ES | ES | conserved hypothetical protein |
| RL4044 | *purE* | NE | NE | ES | ES | putative phosphoribosylaminoimidazole carboxylase catalytic subunit |
| RL4045 | *purK* | NE | NE | ES | ES | putative phosphoribosylaminoimidazole carboxylase ATPase subunit |
| RL4062 |  | NE | NE | ES | ES | putative amidohydrolase |
| RL4073 | *hss* | NE | NE | ES | ES | putative homospermidine synthase |
| RL4117 | *glgA* | NE | NE | ES | ES | putative glycogen synthase |
| RL4183 |  | NE | NE | ES | ES | putative transmembrane protein |
| RL4290 |  | NE | NE | ES | ES | conserved hypothetical protein |
| RL4295 |  | NE | NE | ES | ES | putative acetyltransferase |
| RL4297 |  | NE | NE | ES | ES | putative foldase/peptidyl-prolyl cis-trans isomerase |
| RL4309 |  | NE | NE | ES | ES | putative transmembrane protein |
| RL4333 |  | NE | NE | ES | ES | putative phospholipid/glycerol acyltransferase |
| RL4354 | *xerD* | NE | NE | DE | ES | putative tyrosine recombinase |
| RL4356 |  | NE | NE | DE | DE | conserved hypothetical protein |
| RL4362 |  | NE | NE | DE | DE | putative cobalamin synthesis protein |
| RL4363 | *dacC* | NE | NE | DE | DE | putative penicillin-binding protein precursor |
| RL4386* | *cheW* | NE | NE | DE | ES/DE | putative chemotaxis protein |
| RL4493 | *gpsA* | NE | NE | DE | ES | putative glycerol-3-phosphate dehydrogenase [NAD(P)+] |
| RL4497 |  | NE | NE | DE | DE | putative transmembrane protein |
| RL4498 |  | NE | NE | DE | DE | putative transmembrane protein |
| RL4503 |  | NE | NE | ES | ES | conserved hypothetical protein |
| RL4515 | *argG* | NE | NE | DE | ES | putative argininosuccinate synthase |
| RL4542 | *ispZ* | NE | NE | ES | ES | putative intracellular septation protein |
| RL4599 |  | NE | NE | ES | ES | putative lysyl-tRNA synthetase homolog |
| RL4602 | *nagA* | NE | NE | ES | ES | putative N-acetylglucosamine-6-phosphate deacetylase |
| RL4603 |  | NE | NE | ES | ES | putative aminotransferase |
| RL4631 |  | NE | NE | DE | DE | putative transmembrane transporter protein |
| RL4632 |  | NE | NE | DE | DE | putative DeoR family transcriptional regulator (repressor) |
| RL4638 |  | NE | NE | DE | DE | putative pyruvate carboxylase |
| RLt57 |  | NE | NE | DE | DE | tRNA Thr |

HMM classifications ES; essential, DE; growth-defective, NE; neutral.

*manual curation

**Table S7**. Rlv3841 genes (142) classified as nodule bacteria-specific.

| **Gene** | **Name** | **Rhizosphere** | **Root colonized** | **Nodule bacteria** | **Bacteroids** | **Description** |
| --- | --- | --- | --- | --- | --- | --- |
| pRL100010 |  | NE | NE | DE | NE | conserved hypothetical protein |
| pRL100061 |  | NE | NE | DE | NE | pseudogene, part of an ATP-binding component of ABC transporter PAAT?or POPT (contiguous genes) |
| pRL100062 |  | NE | NE | DE | NE | putative permease component of ABC transporter POPT |
| pRL100072 |  | NE | NE | ES | NE | putative SBP of ABC transporter PAAT |
| pRL100125 |  | NE | NE | DE | NE | putative transposase |
| pRL100126 |  | NE | NE | DE | NE | putative transposase |
| pRL100150 |  | NE | NE | ES | NE | hypothetical protein |
| pRL100168 |  | NE | NE | ES | NE | putative transposase part of insertion sequence |
| pRL100180 | *nodM* | NE | NE | DE | NE | glucosamine--fructose-6-phosphate aminotransferase NodM |
| pRL100204 |  | NE | NE | DE | NE | pseudogene |
| pRL100208 | *fixG* | NE | NE | DE | NE | transmembrane nitrogen fixation cation transport protein FixG |
| pRL100218 |  | NE | NE | DE | NE | putative mobilization protein |
| pRL100220 |  | NE | NE | DE | NE | conserved hypothetical protein |
| pRL100221 |  | NE | NE | DE | NE | conserved hypothetical protein |
| pRL100261 |  | NE | NE | ES | NE | putative LacI family transcriptional regulator (repressor) |
| pRL100292 |  | NE | NE | DE | NE | putative citrate lyase beta chain |
| pRL100293 |  | NE | NE | DE | NE | putative MaoC dehydratase family protein |
| pRL100294 | *mccc1* | NE | NE | DE | NE | putative methylcrotonyl-CoA carboxylase alpha chain |
| pRL100436 | *accC* | NE | NE | ES | NE | putative biotin carboxylase |
| pRL100437 | *accB* | NE | NE | ES | NE | putative biotin carboxyl carrier protein of acetyl-CoA carboxylase |
| pRL110085 |  | NE | NE | DE | NE | putative cycloisomerase |
| pRL110176 |  | NE | NE | DE | NE | conserved hypothetical exported protein |
| pRL110202 |  | NE | NE | DE | NE | conserved hypothetical protein |
| pRL110203 |  | NE | NE | DE | NE | putative urocanate hydratase |
| pRL110283 |  | NE | NE | ES | NE | putative ArsR family transcriptional regulator |
| pRL110509 |  | NE | NE | DE | NE | putative oxidoreductase |
| pRL120070 |  | NE | NE | DE | NE | putative TetR family transcriptional regulator |
| pRL120071 |  | NE | NE | DE | NE | putative SBP of ABC transporter PAAT (S. mel SBP homologue SMb21135 induced by galactosamine, glucosamine) |
| pRL120198 |  | NE | NE | ES | NE | putative L-xylulose kinase |
| pRL120320 |  | NE | NE | ES | NE | putative acetylase |
| pRL120453 |  | NE | NE | ES | NE | putative TIM-barrel fold metal-dependent hydrolase |
| pRL120560 |  | NE | NE | DE | NE | putative ATP-binding component of ABC transporter CUT1 (sucrose?) |
| pRL120578 |  | NE | NE | DE | NE | conserved hypothetical protein |
| pRL120579 |  | NE | NE | DE | NE | putative transmembrane protein |
| pRL120614 |  | NE | NE | DE | NE | putative type III effector Hrp-dependent outers (Hop) (Hrp - Hypersensitive response and pathogenicity) |
| pRL120696 |  | NE | NE | DE | NE | putative RND family efflux transporter (membrane fusion protein) |
| pRL70050 |  | NE | NE | DE | NE | hypothetical protein |
| pRL70090 |  | NE | NE | DE | NE | hypothetical protein |
| pRL70091 | *traI* | NE | NE | DE | NE | putative autoinducer synthesis protein ? Or putative conjugal transfer protein TraI |
| pRL70152 |  | NE | NE | DE | NE | putative crown gall-like type IV secretion system protein |
| pRL70170 |  | NE | NE | DE | NE | conserved hypothetical protein |
| pRL70176 |  | NE | NE | ES | NE | putative transposase-related protein |
| pRL80041 | *hisD* | NE | NE | DE | NE | putative histidinol dehydrogenase |
| pRL80081 |  | NE | NE | DE | NE | putative hydrolase |
| pRL80126 |  | NE | NE | DE | NE | conserved hypothetical protein |
| RL_RF0072 |  | NE | NE | DE | NE | Conserved RNA feature suhB |
| RL_RF0073 |  | NE | NE | DE | NE | Conserved RNA feature suhB |
| RL0006 | *secB* | NE | NE | DE | NE | putative protein-export protein SecB |
| RL0007 |  | NE | NE | DE | NE | putative transmembrane FxsA family protein |
| RL0008 |  | NE | NE | DE | NE | conserved hypothetical protein |
| RL0054 | *glcB* | NE | NE | DE | NE | putative malate synthase |
| RL0104 |  | NE | NE | DE | NE | putative ribonuclease |
| RL0273 |  | NE | NE | DE | NE | putative LysR family transcriptional regulator |
| RL0274 |  | NE | NE | DE | NE | conserved hypothetical protein |
| RL0324 |  | NE | NE | DE | NE | conserved hypothetical protein |
| RL0345 |  | NE | NE | ES | NE | conserved hypothetical protein |
| RL0392 |  | NE | NE | DE | NE | putative hemolysin-like protein |
| RL0734 | *fliQ* | NE | NE | DE | NE | putative flagellar biosynthetic protein FliQ |
| RL0915 | *dgoA* | NE | NE | DE | NE | putative 2-dehydro-3-deoxy-6-phosphogalactonate aldolase |
| RL0916 | *dgoK* | NE | NE | DE | NE | putative 2-dehydro-3-deoxygalactonokinase |
| RL0922 | *kup* | NE | NE | DE | NE | putative potassium uptake transport system protein |
| RL0929 |  | NE | NE | DE | NE | hypothetical protein |
| RL0930 |  | NE | NE | DE | NE | putative ribonuclease HII |
| RL0945 | *aroA* | NE | NE | ES | NE | putative 3-phosphoshikimate 1-carboxyvinyltransferase |
| RL1013 |  | NE | NE | ES | NE | conserved hypothetical protein |
| RL1086 | *ppdK* | NE | NE | DE | NE | pyruvate, phosphate dikinase (pyruvate, orthophosphate dikinase) |
| RL1166 |  | NE | NE | DE | NE | putative ribonuclease-L-PSP family protein |
| RL1167 |  | NE | NE | DE | NE | putative TetR family transcriptional regulator |
| RL1316 | *tam* | NE | NE | DE | NE | putative trans-aconitate 2-methyltransferase |
| RL1496 |  | NE | NE | ES | NE | putative SIS (Sugar ISomerase)-RpiR family transcriptional regulator |
| RL1522 |  | NE | NE | DE | NE | putative transmembrane MscS mechanosensitive ion channel |
| RL1650 |  | NE | NE | DE | NE | putative (DL)-glycerol-3-phosphatase |
| RL1651 | *ubiG* | NE | NE | DE | NE | putative 3-demethylubiquinone-9 3-methyltransferase |
| RL1691 | *hupA* | NE | NE | ES | NE | putative DNA-binding protein HU |
| RL1800 |  | NE | NE | DE | NE | putative transmembrane protein |
| RL2084 |  | NE | NE | DE | NE | putative acetyltransferase |
| RL2087 | *aat* | NE | NE | DE | NE | putative leucyl/phenylalanyl-tRNA--protein transferase |
| RL2102 | *cspA* | NE | NE | ES | NE | putative cold shock protein |
| RL2187 |  | NE | NE | DE | NE | hypothetical protein |
| RL2211 |  | NE | NE | DE | NE | conserved hypothetical protein |
| RL2213 | *clpA* | NE | NE | ES | NE | putative ATP-dependent Clp protease ATP-binding subunit |
| RL2227 | *ecfE* | NE | NE | DE | NE | putative transmembrane protease |
| RL2324 |  | NE | NE | DE | NE | putative ROK family transcriptional regulator |
| RL2368 |  | NE | NE | ES | NE | putative GntR family transcriptional regulator |
| RL2435 |  | NE | NE | DE | NE | putative TolC efflux protein, copper tolerence protein, CopB |
| RL2451 |  | NE | NE | ES | NE | pseudogene |
| RL2473 | *metG* | NE | NE | ES | NE | putative methionyl-tRNA synthetase |
| RL2526 |  | NE | NE | DE | NE | putative oxidoreductase |
| RL2553 |  | NE | NE | DE | NE | conserved hypothetical protein |
| RL2554 |  | NE | NE | DE | NE | hypothetical exported protein |
| RL2569 |  | NE | NE | DE | NE | conserved hypothetical protein |
| RL2607 | *kptA* | NE | NE | DE | NE | putative RNA 2'-phosphotransferase |
| RL2608 | *purQ* | NE | NE | DE | NE | putative phosphoribosylformylglycinamidine synthase I |
| RL2609 |  | NE | NE | DE | NE | conserved hypothetical exported protein |
| RL2643 | *dksA* | NE | NE | DE | NE | putative DnaK suppressor protein |
| RL2644 |  | NE | NE | DE | NE | conserved hypothetical protein |
| RL2658 |  | NE | NE | DE | NE | conserved hypothetical protein |
| RL2659 |  | NE | NE | DE | NE | putative SBP of ABC transporter Unclass |
| RL2716 |  | NE | NE | DE | NE | putative DeoR family transcriptional regulator (repressor) |
| RL2738 |  | NE | NE | DE | NE | putative two-component sensor/regulator; histidine kinase |
| RL2743 |  | NE | NE | ES | NE | putative DeoR family transcriptional regulator (repressor) |
| RL2916 |  | NE | NE | ES | NE | putative siderophore interacting protein |
| RL3232 |  | NE | NE | ES | NE | conserved hypothetical protein with cupin 2 domain |
| RL3236 |  | NE | NE | ES | NE | putative transmembrane anti-sigma factor, AsfE, ECF26 gene organisation (lpp, ecf, asf) |
| RL3254 | *hflK* | NE | NE | DE | NE | putative transmembrane serine protease |
| RL3295 | *recN* | NE | NE | ES | NE | putative DNA repair protein |
| RL3320 |  | NE | NE | DE | NE | putative signalling and peptidoglycan binding protein |
| RL3321 |  | NE | NE | DE | NE | putative DnaJ family chaperone |
| RL3322 | *pfp* | NE | NE | DE | NE | putative pyrophosphate--fructose 6-phosphate 1-phosphotransferase |
| RL3334 | *rnsA* | NE | NE | DE | NE | putative ribonuclease I |
| RL3353 |  | NE | NE | ES | NE | putative ATP-binding component of ABC transporter FeT |
| RL3374A |  | NE | NE | ES | NE | conserved hypothetical protein |
| RL3430 |  | NE | NE | DE | NE | putative ArsR family transcriptional regulator |
| RL3452 |  | NE | NE | DE | NE | conserved hypothetical protein |
| RL3455 |  | NE | NE | DE | NE | putative MarR family transcriptional regulator |
| RL3596 |  | NE | NE | DE | NE | conserved hypothetical protein |
| RL3612 | *ilvD* | NE | NE | DE | NE | putative L-arabinonate dehydratase or 2-keto-3-deoxy-L-arabinoate dehydratase (elevated on MA L-arabinose and galactose Rlv3841) |
| RL3613 |  | NE | NE | DE | NE | putative arabinose dehydrogenase (elevated on MA L-arabinose and galactose Rlv3841) |
| RL3614 |  | NE | NE | DE | NE | putative hydratase, has hydrolase (aligned to COG3802) (elevated on MA L-arabinose and galactose Rlv3841) |
| RL3705 |  | NE | NE | DE | NE | putative anti-anti-sigma factor; two-component sensor/regulator; transcriptional regulator, PhyR, ECF15 gene organisation (histidine kinase, response regulator, asf, ecf) |
| RL3766 | *rpoH* | NE | NE | ES | NE | putative RNA polymerase sigma-32 factor (heat shock) |
| RL3974 |  | NE | NE | DE | NE | putative thioesterase family protein |
| RL3986 | *ruvC* | NE | NE | DE | NE | putative Holliday junction endodeoxyribonuclease RuvC |
| RL4011 | *pgk* | NE | NE | ES | NE | putative phosphoglycerate kinase |
| RL4017 | *rpmE* | NE | NE | DE | NE | putative 50S ribosomal protein L31 |
| RL4018 |  | NE | NE | DE | NE | putative ATP-binding:permease (ABC:IMP) component of ABC transporter Export |
| RL4203 | *talB* | NE | NE | ES | NE | putative transaldolase B |
| RL4210 | *cysZ* | NE | NE | DE | NE | putative cysteine biosynthesis protein |
| RL4336 |  | NE | NE | ES | NE | putative acetyltransferase |
| RL4392 | *fdsB* | NE | NE | DE | NE | putative NAD-dependent formate dehydrogenase beta subunit |
| RL4430 |  | NE | NE | DE | NE | putative 3-oxoacyl-[acyl-carrier-protein] reductase |
| RL4431 |  | NE | NE | DE | NE | conserved hypothetical exported protein |
| RL4490 |  | NE | NE | DE | NE | conserved hypothetical protein |
| RL4491 |  | NE | NE | DE | NE | conserved hypothetical protein |
| RL4492 |  | NE | NE | DE | NE | conserved hypothetical protein |
| RL4516 |  | NE | NE | DE | NE | putative oxidoreductase |
| RL4517 |  | NE | NE | DE | NE | putative LysE family efflux protein |
| RL4535 |  | NE | NE | ES | NE | conserved hypothetical exported protein |
| RL4610 |  | NE | NE | ES | NE | putative thiamine pyrophosphokinase |
| RL4641 |  | NE | NE | DE | NE | pseudogene, transposase |
| RL4670 |  | NE | NE | DE | NE | conserved hypothetical protein |
| RLt48 |  | NE | NE | DE | NE | tRNA Pro |

HMM classifications ES; growth-essential, DE; growth-defective, NE; growth-neutral.

**Table S8**. Rlv3841 genes (24) classified as bacteroid-specific.

| **Gene** | **Name** | **Rhizosphere** | **Root colonized** | **Nodule bacteria** | **Bacteroids** | **Description** |
| --- | --- | --- | --- | --- | --- | --- |
| pRL100159 | *nifE* | NE | NE | NE | DE | putative nitrogenase iron-molybdenum cofactor biosynthesis protein NifE |
| pRL100162 | *nifH* | NE | NE | NE | DE | nitrogenase iron protein NifH |
| pRL100195 | *nifB* | NE | NE | NE | DE | FeMo cofactor biosynthesis protein NifB |
| pRL100196 | *nifA* | NE | NE | NE | DE | transcriptional regulator NifA |
| pRL100197 | *fixX* | NE | NE | NE | DE | ferredoxin-like protein FixX |
| pRL100198 | *fixC* | NE | NE | NE | DE | nitrogen fixation protein FixC |
| pRL100432 |  | NE | NE | NE | ES | putative lactam utilization protein |
| RL0037 | *pckA* | NE | NE | NE | DE | putative phosphoenolpyruvate carboxykinase |
| RL0119 |  | NE | NE | NE | ES | putative ribosomal RNA small subunit methyltransferase |
| RL0401 |  | NE | NE | NE | DE | putative universal stress protein |
| RL0501 |  | NE | NE | NE | DE | putative orotate phosphoribosyltransferase |
| RL0618 |  | NE | NE | NE | DE | conserved hypothetical protein |
| RL0901 |  | NE | NE | NE | DE | conserved hypothetical protein |
| RL1217 |  | NE | NE | NE | ES | putative TetR family transcriptional regulator |
| RL1717 |  | NE | NE | NE | DE | putative lactoylglutathione lyase |
| RL1718 |  | NE | NE | NE | DE | putative transmembrane protein |
| RL1736 | *smf* | NE | NE | NE | DE | putative SMF family DNA protecting protein DprA involved in uptake of DNA and natural bacterial competence |
| RL3425 | *dctB* | NE | NE | NE | DE | putative two-component sensor/regulator; histidine kinase (C4-dicarboxylate transport) |
| RL3426 | *dctD* | NE | NE | NE | DE | putative two-component sensor/regulator; transcriptional regulator C4-dicarboxylate transport (sigma-54) |
| RL3462 |  | NE | NE | NE | DE | conserved hypothetical protein |
| RL3479 |  | NE | NE | NE | ES | putative GTP-dependent nucleic acid-binding protein |
| RL3484 | *petC* | NE | NE | NE | ES | putative cytochrome c1 precursor |
| RL3577 | *guaD* | NE | NE | NE | DE | putative guanine deaminase |
| RL3666 |  | NE | NE | NE | DE | conserved hypothetical protein |

HMM classifications: ES; growth-essential, DE; growth-defective, NE; growth-neutral.

**Table S9.** Rlv3841 genes classified as AD at stages from rhizosphere to symbiosis.

| **Gene** | **Designation** | **Description** | **Riley** |
| --- | --- | --- | --- |
| **Rhizosphere (84 genes)** | | |  |
| pRL100050 |  | putative integrase/recombinase protein | 5.1.2 |
| pRL100055 |  | putative acetolactate synthase subunit | 3.1.21 |
| pRL100285 |  | conserved hypothetical protein | 0.0.1 |
| pRL100357 |  | putative ATP-binding component of ABC transporter CUT2 | 1.5.3 |
| pRL100384 |  | putative esterase/lipase | 3.3.15 |
| pRL110174 |  | putative 3-oxoacyl-[acyl-carrier-protein] reductase | 3.6.0 |
| pRL110209 |  | conserved hypothetical protein | 0.0.2 |
| pRL120438 |  | hypothetical protein | 0.0.0 |
| pRL70118 |  | putative crown gall tumor-like protein | 1.4.1 |
| pRL70119 |  | putative crown-gall tumor-like protein | 1.4.1 |
| pRL7011A |  | pseudogene | 7.2.1 |
| pRL70181 |  | pseudogene | 7.2.1 |
| pRL80059 |  | putative NifS-like cysteine desulfurase/selenocysteine lyase | 3.1.6 |
| pRL80088 |  | putative permease component of ABC transporter CUT2 homoserine transporter (Hynes) | 1.5.3 |
| pRL80089 |  | pseudogene | 7.2.1 |
| pRL80124 | *traBp8* | putative conjugal transfer protein TraB | 5.1.3 |
| pRL90014 | *fixH2* | putative cation transport nitrogen fixation protein | 1.5.2 |
| pRL90137 |  | putative glycosyl transferase | 4.1.4 |
| RL0141 | *cycM* | putative cytochrome c | 3.5.3 |
| RL0142 |  | putative transmembrane permease protein | 1.5.0 |
| RL0255 |  | putative HemeO family (TenA) transcriptional regulator | 6.3.0 |
| RL0526 |  | conserved hypothetical protein | 0.0.2 |
| RL0527 |  | hypothetical protein | 0.0.0 |
| RL0535 | *fixO3* | putative cbb3 cytochrome oxidase subunit FixO | 3.3.22 |
| RL0919 |  | putative LuxR/GerE family transcriptional regulator | 6.3.11 |
| RL0955 |  | putative MipA family outer membrane scaffold protein | 4.1.3 |
| RL1152 |  | putative transmembrane protein | 4.1.1 |
| RL1431 |  | putative transmembrane MFS family permease | 1.5.0 |
| RL1870 | *dksA* | putative DnaK suppressor protein | 1.3.1 |
| RL1871 |  | putative transmembrane cation ATPase transporter | 1.5.2 |
| RL1872 |  | putative universal stress protein | 1.6.1 |
| RL1873 |  | putative pyridoxine oxidase | 3.2.12 |
| RL1874 |  | conserved hypothetical protein | 0.0.1 |
| RL1875 |  | putative transmembrane protein | 4.1.1 |
| RL1876 | *adh* | putative alcohol dehydrogenase | 3.3.15 |
| RL1877 |  | putative protease | 2.1.4 |
| RL1878 |  | putative peptidoglycan binding protein | 4.1.2 |
| RL1879 |  | putative two component sensor/regulator; fused sensor/regulator (nitrogen fixation) | 6.1.3 |
| RL1880 |  | putative FNR/CRP family transcriptional regulator | 6.3.0 |
| RL1881 |  | putative two-component sensor/regulator; transcriptional regulator | 6.1.2 |
| RL1882 |  | conserved hypothetical protein | 0.0.1 |
| RL1883 | *hspF* | putative small heat shock protein | 1.6.1 |
| RL1884 |  | putative phospholipid binding protein | 1.5.5 |
| RL1885 |  | putative beta-lactamase family protein | 7.0.0 |
| RL1886 | *pduW* | putative propionate kinase | 3.3.7 |
| RL1887 |  | putative transmembrane protein | 4.1.1 |
| RL1888 |  | hypothetical exported protein | 0.0.0 |
| RL1889 |  | putative multicopper oxidase | 1.5.2 |
| RL1890 |  | putative transmembrane protein | 4.1.1 |
| RL1891 |  | putative transmembrane protein | 4.1.1 |
| RL1892 |  | putative cation transporting P-type ATPase | 1.5.2 |
| RL1893 |  | putative transmembrane protein (2 membrane spanning domains) with EF hand domain, 58% id with RL2272 | 4.1.1 |
| RL1894 |  | hypothetical protein | 0.0.0 |
| RL1895 |  | hypothetical protein | 0.0.0 |
| RL1923 |  | conserved hypothetical protein | 0.0.1 |
| RL1956 |  | putative transmembrane protein | 4.1.1 |
| RL1957 |  | putative short-chain dehydrogenase/oxidoreductase | 3.3.15 |
| RL1958 |  | putative TetR family transcriptional regulator | 6.3.8 |
| RL1959 | *acpD* | putative acyl carrier protein phosphodiesterase | 3.2.1 |
| RL1960 |  | putative LysR family transcriptional regulator | 6.3.6 |
| RL1961 | *cpdB* | putative 2',3'-cyclic-nucleotide 2'-phosphodiesterase | 3.3.15 |
| RL1983 |  | conserved hypothetical protein | 0.0.2 |
| RL1984 |  | conserved hypothetical protein | 0.0.2 |
| RL1985 | *phr* | putative deoxyribodipyrimidine photo-lyase | 2.2.3 |
| RL1992 | *narK* | putative nitrate transport protein MFS protein | 1.5.4 |
| RL1993 |  | putative SBP of ABC transporter NitT anionic substrate (S. mel SBP homologue SMb21114 induced by N limitation) | 1.5.4 |
| RL1994 | *nasT* | putative two-component sensor/regulator; nitrate reductase transcriptional regulator NasT | 6.1.2 |
| RL1995 |  | putative LacI family transcriptional regulator (repressor) | 6.3.5 |
| RL2064 |  | putative methyltransferase | 3.3.15 |
| RL2065 |  | conserved hypothetical protein | 0.0.2 |
| RL2066 | *sthA* | putative soluble pyridine nucleotide transhydrogenase | 3.2.11 |
| RL2067 | *lldD2* | putative L-lactate dehydrogenase | 3.5.4 |
| RL2068 | *radC* | putative DNA repair protein | 2.2.3 |
| RL2069 | *map1* | putative methionine aminopeptidase | 2.2.10 |
| RL2105 |  | conserved hypothetical protein | 0.0.2 |
| RL2144 |  | conserved hypothetical protein | 0.0.2 |
| RL2145 |  | hypothetical protein | 0.0.0 |
| RL2146 |  | putative transmembrane protein | 4.1.1 |
| RL3231 |  | putative MarR family transcriptional regulator | 6.3.7 |
| RL3394 |  | conserved hypothetical protein | 0.0.2 |
| RL3641 |  | putative transmembrane protein | 4.1.1 |
| RL3642 |  | putative glycosyl hydrolase | 4.1.4 |
| RL3691 |  | conserved hypothetical protein | 0.0.1 |
| RL3692 |  | conserved hypothetical protein | 0.0.1 |
| **Rhizosphere-progressive (0)** | | |  |
| **Rhizosphere and root (172)** | | |  |
| pRL100017 |  | conserved hypothetical protein | 0.0.1 |
| pRL100024 |  | conserved hypothetical protein | 0.0.2 |
| pRL100048 |  | putative phage integrase/tyrosine recombinase | 5.1.2 |
| pRL100049 |  | putative integrase/recombinase protein | 5.1.2 |
| pRL100078 |  | hypothetical protein | 0.0.0 |
| pRL100079 | *qatV6* | putative ATP-binding component of ABC transporter QAT glycine betaine/L-proline transporter | 1.5.1 |
| pRL100080 | *qatW6* | putative permese component of ABC transporter QAT glycine betaine/L-proline transporter | 1.5.1 |
| pRL100081 | *qatX6* | putative SBP of ABC transporter QAT glycine betaine/L-proline transporter | 1.5.1 |
| pRL100082 |  | putative XRE family (HipB) transcriptional regulator | 6.3.0 |
| pRL100083 |  | hypothetical protein | 0.0.0 |
| pRL100201 |  | conserved hypothetical protein | 0.0.2 |
| pRL100286 |  | putative RND family efflux transporter (inner membrane protein) | 1.5.5 |
| pRL100287 |  | putative RND family efflux transporter (membrane fusion protein) | 1.5.5 |
| pRL100355 |  | putative isochorismatase | 3.4.2 |
| pRL100356 |  | putative allophanate hydrolase | 1.4.2 |
| pRL100385 | *ecfM* | putative RNA polymerase ECF sigma factor, family ECF20/ECF01 | 6.2.1 |
| pRL100427 |  | putative permease component of ABC transporter Unclass | 1.5.0 |
| pRL110053 | *optA* | putative SBP of ABC transporter PepT OptA | 1.5.0 |
| pRL110054 | *optD* | putative ATP-binding:ATP-binding (ABC:ABC) component of ABC transporter PepT, OptD | 1.5.0 |
| pRL110106 |  | conserved hypothetical protein | 0.0.1 |
| pRL110149 |  | putative isomerase | 3.3.15 |
| pRL110164 |  | putative gluconate 5-dehydrogenase | 3.3.15 |
| pRL110167 |  | putative permease component of ABC transporter CUT2 | 1.5.3 |
| pRL110173 |  | putative transmembrane protein | 4.1.1 |
| pRL110186 |  | conserved hypothetical protein | 0.0.2 |
| pRL110281 |  | putative SBP of ABC transporter PepT | 1.5.0 |
| pRL110337 |  | putative potassium transport system protein | 1.5.2 |
| pRL110500 |  | conserved hypothetical protein | 0.0.1 |
| pRL110501 |  | putative ribonuclease | 2.1.2 |
| pRL110580 |  | putative insertion sequence/transposase-related protein | 5.1.4 |
| pRL110581 |  | putative transposase-related protein | 5.1.4 |
| pRL120026 |  | putative DeoR family transcriptional regulator (repressor) | 6.3.10 |
| pRL120137 |  | conserved hypothetical protein | 0.0.2 |
| pRL120164 |  | putative SBP of ABC transporter PepT | 1.5.0 |
| pRL120262 |  | putative SBP of ABC transporter FeT | 1.5.0 |
| pRL120288 |  | putative hydrolase | 3.3.15 |
| pRL120309 |  | putative FAD-dependent dehydrogenase | 3.3.15 |
| pRL120310 |  | putative flavodoxin containing oxidoreductase | 3.3.15 |
| pRL120311 |  | putative threonine degradation protein | 3.4.2 |
| pRL120330 |  | putative ATP-binding component of ABC transporter PepT | 1.5.0 |
| pRL120404 | *braC2* | putative SBP of ABC transporter HAAT | 1.5.0 |
| pRL120415 | *dadR* | putative AsnC family transcriptional regulator (alanine catabolic operon regulator) | 6.3.1 |
| pRL120425 |  | putative transmembrane protein | 4.1.1 |
| pRL120571 |  | putative transmembrane tryptophan-rich protein | 4.1.1 |
| pRL120701 |  | putative transmembrane magnesium and cobalt transport protein, CorA family | 1.5.2 |
| pRL120720 |  | conserved hypothetical protein | 0.0.2 |
| pRL120752 |  | putative SBP of ABC transporter PhnT (from homology) | 1.5.0 |
| pRL70117 |  | putative endonuclease | 2.1.1 |
| pRL70158 |  | putative conjugative DNA transfer/component of type IV secretion system | 1.5.5 |
| pRL70159 |  | pseudogene | 7.2.1 |
| pRL70160 |  | conserved hypothetical protein | 0.0.1 |
| pRL70161 |  | putative transposase-related protein | 5.1.4 |
| pRL70162 |  | conserved hypothetical protein | 0.0.1 |
| pRL70163 |  | pseudogene, transposase | 7.2.1 |
| pRL70174 |  | pseudogene, transposase-related protein | 7.2.1 |
| pRL70182 |  | conserved hypothetical exported protein | 0.0.2 |
| pRL70183 |  | conserved hypothetical exported protein | 0.0.2 |
| pRL70184 |  | hypothetical protein | 0.0.0 |
| pRL70185 |  | pseudogene, putative integrase | 7.2.1 |
| pRL70186 |  | pseudogene | 7.2.1 |
| pRL70187 |  | hypothetical protein | 0.0.0 |
| pRL80060 |  | putative SBP of ABC transporter PAAT closest homol mimosine transporter Rhizobium sp. TAL1145 | 1.5.0 |
| pRL80097 |  | hypothetical protein | 0.0.0 |
| pRL80123 | *traFp8* | putative conjugal transfer protein TraF | 5.1.3 |
| pRL90175 | *bdhA2* | putative D-beta-hydroxybutyrate dehydrogenase | 3.3.15 |
| RL0252 |  | putative nodulation protein | 3.3.22 |
| RL0253 | *gfa* | putative glutathione-dependent formaldehyde-activating enzyme | 1.6.1 |
| RL0325 |  | conserved hypothetical exported protein | 0.0.1 |
| RL0326 |  | putative 4Fe-4S ferredoxin protein | 3.5.3 |
| RL0481 |  | putative 3-oxoacyl-[acyl-carrier-protein] reductase | 3.2.1 |
| RL0599 |  | putative transmembrane efflux protein | 1.5.0 |
| RL0715 | *fliL* | putative flagella-related protein FliL | 1.1.1 |
| RL0716 | *fliP* | putative flagellar biosynthetic protein FliP | 1.1.1 |
| RL0717 |  | conserved hypothetical protein (TPR repeat family) | 0.0.2 |
| RL0728 | *flgE1* | putative flagellar hook protein FlgE | 1.1.1 |
| RL0804 |  | conserved hypothetical protein | 0.0.1 |
| RL0836A |  | hypothetical protein | 0.0.0 |
| RL1102 |  | conserved hypothetical protein | 0.0.2 |
| RL1135 |  | putative transmembrane copper resistance protein | 1.5.2 |
| RL1216 |  | conserved hypothetical protein | 0.0.2 |
| RL1407 |  | hypothetical protein | 0.0.0 |
| RL1408 |  | conserved hypothetical protein | 0.0.1 |
| RL1409 |  | conserved hypothetical protein | 0.0.1 |
| RL1444 |  | putative glutathione S-transferase I | 1.6.1 |
| RL1542 |  | conserved hypothetical protein | 0.0.2 |
| RL1655 |  | putative ROK family transcriptional regulator | 6.3.9 |
| RL1683 |  | putative permease component of ABC transporter PhoT | 1.5.0 |
| RL1845 |  | putative exported ErfK/YbiS/YhnG family protein | 0.0.1 |
| RL1846 |  | putative two-component sensor/regulator; transcriptional regulator | 6.1.2 |
| RL1924 |  | conserved hypothetical exported protein (with homology to (neuropeptide) neuromedin U - meaning?) | 0.0.2 |
| RL1925 |  | conserved hypothetical protein with DUF1254 and DUF1214 (COG5361), highest id with pRL70183 (26%) | 0.0.2 |
| RL1926 |  | conserved hypothetical protein with DUF1254 and DUF1214 (COG5361), shows 75% id with pRL70182 | 0.0.2 |
| RL1936 |  | hypothetical protein | 0.0.0 |
| RL1943 |  | conserved hypothetical protein | 0.0.2 |
| RL1944 |  | conserved hypothetical protein | 0.0.2 |
| RL1986 |  | hypothetical protein | 0.0.0 |
| RL1987 |  | conserved hypothetical exported protein | 0.0.2 |
| RL1988 |  | putative glycerol phosphatase | 3.3.8 |
| RL1989 | *nasA* | putative assimilatory nitrate reductase | 3.3.22 |
| RL1990 | *nirD* | putative 2Fe-2S rieske nitrite reductase small subunit | 3.3.22 |
| RL1991 | *nasD* | putative assimilatory siroheme nitrite reductase [NAD(P)H] | 3.3.22 |
| RL2009 |  | putative peptidase | 2.1.4 |
| RL2010 |  | putative ATP-binding component of ABC transporter POPT | 1.5.0 |
| RL2011 |  | putative permease component of ABC transporter POPT | 1.5.0 |
| RL2012 |  | putative permease component of ABC transporter POPT | 1.5.0 |
| RL2013 |  | putative SBP of ABC transporter POPT | 1.5.0 |
| RL2014 |  | putative adenine deaminase | 3.3.17 |
| RL2053 |  | conserved hypothetical protein | 0.0.2 |
| RL2077 |  | putative acetyltransferase | 3.3.15 |
| RL2078 |  | conserved hypothetical protein | 0.0.2 |
| RL2079 |  | conserved hypothetical protein | 0.0.2 |
| RL2080 |  | putative acetyltransferase | 3.3.15 |
| RL2081 |  | putative transmembrane protein | 4.1.1 |
| RL2091 |  | putative outer membrane protein | 4.1.3 |
| RL2092 | *aspB* | putative aspartate aminotransferase | 3.4.2 |
| RL2093 |  | conserved hypothetical protein | 0.0.2 |
| RL2094 | *phaC* | putative poly(3-hydroxyalkanoate) polymerase (PHA synthase) | 2.2.8 |
| RL2095 |  | conserved hypothetical protein | 0.0.2 |
| RL2120 |  | conserved hypothetical protein | 0.0.2 |
| RL2121 |  | putative transmembrane protein | 4.1.1 |
| RL2175 |  | conserved hypothetical protein | 0.0.2 |
| RL2203 | *aapQ* | putative permease component of ABC transporter PAAT (general L-amino acid, Aap system) | 1.5.1 |
| RL2204 | *aapJ* | putative SBP of ABC transporter PAAT (general L-amino acid, Aap system) | 1.5.1 |
| RL2205 | *metC* | putative cystathionine beta-lyase | 3.1.14 |
| RL2260 |  | conserved hypothetical protein | 0.0.1 |
| RL2261 |  | conserved hypothetical exported protein | 0.0.1 |
| RL2443 |  | putative LysR family transcriptional regulator | 6.3.6 |
| RL2489A |  | conserved hypothetical protein | 0.0.2 |
| RL2895 |  | putative SBP of ABC transporter CUT1 | 1.5.3 |
| RL2896 |  | putative permease component of ABC transporter CUT1 | 1.5.3 |
| RL2986 |  | putative AsnC family transcriptional regulator | 6.3.1 |
| RL3087 |  | putative LysR family transcriptional regulator | 6.3.6 |
| RL3090 |  | putative methyltransferase | 3.3.15 |
| RL3104 |  | putative oxidoreductase | 3.3.15 |
| RL3105 |  | putative hydrolase | 3.3.15 |
| RL3183 |  | putative transmembrane protein | 4.1.1 |
| RL3220 |  | putative HxlR family transcriptional regulator (activator) | 6.3.0 |
| RL3221 |  | putative nucleoside-diphosphate-sugar epimerase | 3.3.18 |
| RL3303 |  | putative transmembrane transporter protein | 1.5.0 |
| RL3304 | *galE* | putative UDP-glucose 4-epimerase | 4.1.2 |
| RL3363 |  | putative mannose-6-phosphate isomerase | 4.1.2 |
| RL3364 |  | hypothetical protein | 0.0.0 |
| RL3377 | *cinR* | putative LuxR/GerE family transcriptional regulator (AHL-dependent) | 6.3.11 |
| RL3378 | *cinI* | putative autoinducer synthesis protein, CinS is v small protein, translationally coupled | 1.6.1 |
| RL3379 |  | putative two-component sensor/regulator; transcriptional regulator | 6.1.2 |
| RL3395 |  | putative transmembrane protein | 4.1.1 |
| RL3396 |  | putative AraC family transcriptional regulator (activator) | 6.3.2 |
| RL3397 | *neo* | putative aminoglycoside 3'-phosphotransferase | 1.4.3 |
| RL3398 |  | conserved hypothetical protein | 0.0.1 |
| RL3399 |  | putative proline-rich protein | 0.0.1 |
| RL3403 |  | conserved hypothetical protein | 0.0.2 |
| RL3404 | *tdh* | putative L-threonine 3-dehydrogenase | 3.4.2 |
| RL3405 | *kbl* | putative 2-amino-3-ketobutyrate coenzyme A ligase | 3.4.2 |
| RL3406 |  | putative biotin synthesis protein | 3.2.2 |
| RL3407 | *recQ* | putative ATP-dependent DNA helicase RecQ | 2.2.3 |
| RL3412 |  | conserved hypothetical protein | 0.0.1 |
| RL3413 | *norM* | putative multidrug resistance protein NorM (Na(+)/drug antiporter) (multidrug-efflux transporter) | 1.5.5 |
| RL3414 | *tetA* | putative tetracycline resistance protein | 1.4.3 |
| RL3415 |  | putative oxidoreductase | 3.3.15 |
| RL3416 |  | putative transmembrane protease protein | 2.1.4 |
| RL3417 |  | putative transmembrane protein | 4.1.1 |
| RL3418 |  | putative response regulator of LytR family transcriptional regulator | 6.1.2 |
| RL3469 |  | conserved hypothetical protein | 0.0.1 |
| RL3470 |  | putative transmembrane GGDEF/EAL sensory box protein | 6.6.0 |
| RL3655 | *pssE* | putative glycosyltransferase | 4.1.4 |
| RL3824 |  | putative transmembrane protein | 4.1.1 |
| RL3836 |  | putative transmembrane protein | 4.1.1 |
| RL3863 |  | putative GntR family transcriptional regulator | 6.3.3 |
| RL3889 |  | putative transmembrane protein | 4.1.1 |
| RL3919 |  | hypothetical protein | 0.0.0 |
| RL4186 |  | putative oxidoreductase | 3.3.15 |
| RL4592 |  | putative glycosyl transferase | 4.1.4 |
| **Root (33 genes)** | | |  |
| pRL100202 |  | conserved hypothetical protein | 0.0.2 |
| pRL110338 |  | hypothetical protein | 0.0.0 |
| pRL120025 |  | conserved hypothetical protein | 0.0.1 |
| pRL120412 | *hyuB* | putative hydantoin utilization protein B | 3.1.0 |
| pRL120413 |  | putative 3-oxoacyl-[acyl-carrier-protein] reductase | 3.6.1 |
| pRL120414 |  | putative HpcH/HpaI family aldolase | 3.3.15 |
| pRL120772 |  | putative ATP-binding component of ABC transporter PepT | 1.5.0 |
| pRL70116 |  | putative endonuclease | 2.1.1 |
| pRL90283 |  | putative ATP-binding component of ABC transporter CUT1 (S. mel SBP homologue SMb21652 induced by lactose, lactulose) | 1.5.3 |
| RL0482 |  | putative TetR family transcriptional regulator | 6.3.8 |
| RL0956 |  | putative para-hydroxybenzoate--polyprenyltransferase | 3.2.8 |
| RL1215 |  | putative transmembrane protein | 4.1.1 |
| RL1449 |  | conserved hypothetical protein | 0.0.1 |
| RL1450 |  | hypothetical protein | 0.0.0 |
| RL2176 |  | conserved hypothetical protein | 0.0.2 |
| RL2177 |  | hypothetical protein | 0.0.0 |
| RL2258 |  | hypothetical protein | 0.0.0 |
| RL2259 |  | conserved hypothetical exported protein | 0.0.1 |
| RL2429 | *cyaA2* | putative adenylate cyclase | 3.3.15 |
| RL2430 |  | conserved hypothetical protein | 0.0.2 |
| RL2491 |  | conserved hypothetical exported protein | 0.0.2 |
| RL3182 |  | conserved hypothetical protein | 0.0.2 |
| RL3218 |  | putative TetR family transcriptional regulator | 6.3.8 |
| RL3219 |  | putative MFS family transmembrane transporter | 1.5.0 |
| RL3365 |  | conserved hypothetical protein | 0.0.1 |
| RL3366 |  | putative flavoprotein | 3.5.3 |
| RL3400 |  | conserved hypothetical exported protein | 0.0.1 |
| RL3401 |  | conserved hypothetical protein | 0.0.1 |
| RL3654 | *pssD* | putative polysaccharide biosynthesis protein | 4.1.4 |
| RL3656 |  | putative lipase | 3.4.4 |
| RL4593 |  | putative beta-mannosidase | 4.1.4 |
| RL4594 |  | conserved hypothetical protein | 0.0.2 |
| RL4595 |  | conserved hypothetical protein | 0.0.2 |
| **Root-progressive (0)** | | |  |
| **Nodule-general (11 genes)** | | |  |
| pRL70131 |  | pseudogene | 7.2.1 |
| pRL70132 |  | pseudogene | 7.2.1 |
| RL0749 | *aglK* | putative ATP-binding component of ABC transporter CUT1 alpha-glucoside-transporter | 1.5.3 |
| RL0750 |  | conserved hypothetical protein | 0.0.2 |
| RL3628 |  | putative sugar decarboxylase | 3.3.15 |
| RL3629 |  | putative glycosyltransferase | 4.1.2 |
| RL3630 |  | putative glycosyltransferase | 4.1.4 |
| RL3631 |  | putative glycosyltransferase | 4.1.4 |
| RL3632 |  | putative ATP-binding:permease (ABC:IMP) component of ABC transporter protein Export | 1.5.5 |
| RL3633 | *exoU* | putative UDP-hexose transferase | 4.1.2 |
| RL4139 |  | putative transmembrane GGDEF/EAL sensory box protein | 6.6.0 |
| **Nodule bacteria (207 genes)** | | |  |
| pRL100206 | *fixO1* | putative cytochrome oxidase subunit | 3.5.1 |
| pRL100206A | *fixQ1* | putative component of cytochrome oxidase | 3.5.1 |
| pRL100207 | *fixP1* | putative cytochrome oxidase subunit | 3.5.1 |
| pRL100228 |  | putative SBP of ABC transporter Unclass | 1.5.0 |
| pRL100303 |  | putative 2-oxoisovalerate dehydrogenase | 3.4.2 |
| pRL100439 |  | putative inositol degradation protein | 3.4.4 |
| pRL100440 |  | putative SBP of ABC transporter CUT1 | 1.5.3 |
| pRL100441 |  | putative permease component of ABC transporter CUT1 | 1.5.3 |
| pRL100442 |  | putative permease component of ABC transporter CUT1 | 1.5.3 |
| pRL100443 |  | putative ATP-binding component of ABC transporter CUT1 | 1.5.3 |
| pRL100467 |  | conserved hypothetical protein | 0.0.2 |
| pRL110019 |  | putative transmembrane transporter protein | 1.5.0 |
| pRL110232 |  | putative transmembrane cyclic nucleotide-binding ion channel | 1.5.0 |
| pRL110233 |  | putative permease component of ABC transporter Export | 1.5.0 |
| pRL110234 |  | putative permease component of ABC transporter Export | 1.5.0 |
| pRL110368 |  | putative phosphoesterase/regulator | 6.0.0 |
| pRL110510 |  | conserved hypothetical protein | 0.0.2 |
| pRL110511 |  | putative glycosyl hydrolase | 4.1.4 |
| pRL110512 |  | conserved hypothetical protein | 0.0.1 |
| pRL110513 |  | putative ATP-binding component of ABC transporter PepT | 1.5.0 |
| pRL110514 |  | putative ATP-binding component of ABC transporter PepT | 1.5.0 |
| pRL110515 |  | putative permease component of ABC transporter PepT | 1.5.0 |
| pRL110516 |  | putative permease component of ABC transporter PepT | 1.5.0 |
| pRL110517 |  | putative SBP of ABC transporter PepT | 1.5.0 |
| pRL120004 |  | putative dihydrodipicolinate synthase | 3.1.13 |
| pRL120005 |  | putative GntR family transcriptional regulator | 6.3.3 |
| pRL120006 |  | putative SBP of ABC transporter CUT1 | 1.5.3 |
| pRL120007 |  | putative permease component of ABC transporter CUT1 | 1.5.3 |
| pRL120014 |  | putative ATP-binding component of ABC transporter CUT1 | 1.5.3 |
| pRL120094 | *tfdB* | putative dichlorophenol monooxygenase/hydroxylase | 1.4.2 |
| pRL120095 |  | putative permease component of ABC transporter HAAT | 1.5.1 |
| pRL120138 |  | putative cobalamin synthesis protein (CobW family) | 3.2.3 |
| pRL120139 |  | conserved hypothetical protein | 0.0.2 |
| pRL120199 |  | putative DeoR family transcriptional regulator (repressor) | 6.3.10 |
| pRL120200 | *eryG* | putative SBP of ABC transporter CUT2 erythritol | 1.5.3 |
| pRL120201 | *eryF* | putative permease component of ABC transporter CUT2 erythritol transporter | 1.5.3 |
| pRL120202 | *eryE* | putative ATP-binding:ATP-binding (ABC:ABC) component of ABC transporter CUT2, erythritol | 1.5.3 |
| pRL120203 | *eryH* | putative periplasmic lipoprotein, involved in erythritol uptake/metabolism? | 1.5.3 |
| pRL120207 | *eryD* | putative DeoR family transcriptional regulator (repressor) | 6.3.10 |
| pRL120285 |  | putative ATP-binding:ATP-binding (ABC:ABC) component of ABC transporter CUT2 (S. mel SBP homologue SMa0203 induced by galactose, L-arabinose, fucose) | 1.5.3 |
| pRL120537 |  | putative FAD/NAD/ferredoxin protein | 3.3.15 |
| pRL120538 |  | putative Rieske 2Fe2S dioxygenase | 1.4.2 |
| pRL120557 |  | putative permease component of ABC transporter CUT1 (sucrose?) | 1.5.3 |
| pRL120558 |  | putative permease component of ABC transporter CUT1 (sucrose?) | 1.5.3 |
| pRL120559 |  | putative glycosyl hydrolase, beta-fructosidase | 2.1.3 |
| pRL120617 |  | putative transmembrane MFS family transporter | 1.5.0 |
| pRL120722 |  | putative alpha-amylase | 3.4.3 |
| pRL70072 |  | pseudogene | 7.2.1 |
| pRL70149 |  | putative transglycosylase SLT domain protein | 4.1.2 |
| pRL70150 |  | putative transmembrane protein | 4.1.1 |
| pRL70151 |  | putative transmembrane protein | 4.1.1 |
| pRL90114 |  | putative glycerophosphoryl diester phosphodiesterase | 3.3.8 |
| pRL90115 |  | putative hydrolase | 3.3.15 |
| pRL90116 | *rbtD2* | putative ribitol 2-dehydrogenase | 3.3.21 |
| pRL90117 |  | putative D-ribulokinase/ribitol kinase | 3.3.9 |
| pRL90246 |  | putative ATP-binding component of ABC transporter protein ABC:ABC CUT2 | 1.5.3 |
| pRL90247 |  | putative permease component of ABC transporter CUT2 | 1.5.3 |
| pRL90248 |  | putative xenobiotic monooxygenase | 1.4.2 |
| pRL90249 |  | putative SBP of ABC transporter PepT | 1.5.0 |
| pRL90250 |  | putative permease component of ABC transporter PepT | 1.5.0 |
| pRL90251 |  | putative permease component of ABC transporter PepT | 1.5.0 |
| pRL90252 |  | putative ATP-binding component of ABC transporter PepT | 1.5.0 |
| pRL90253 |  | putative ATP-binding component of ABC transporter PepT | 1.5.0 |
| pRL90298 |  | conserved hypothetical protein | 0.0.2 |
| RL0103 | *gabR* | putative MerR family transcriptional regulator | 6.3.12 |
| RL0105 |  | conserved hypothetical protein | 0.0.2 |
| RL0122 |  | putative oxidoreductase | 3.3.15 |
| RL0176 | *phnH* | putative phosphonate utilisation protein H | 3.3.13 |
| RL0177 | *phnG* | putative phosphonate utilisation protein G | 3.3.13 |
| RL0178 |  | putative GntR family transcriptional regulator | 6.3.3 |
| RL0339 |  | putative Mg2+ chelatase family protein | 7.0.0 |
| RL0455 | *mcpT* | putative methyl-accepting chemotaxis protein | 1.1.1 |
| RL0606 |  | putative enoyl-CoA hydratase | 3.4.4 |
| RL0856 |  | conserved hypothetical protein | 0.0.2 |
| RL0912 |  | putative GGDEF/EAL sensory box protein | 6.6.0 |
| RL0913 |  | putative PRC family protein | 7.0.0 |
| RL0914 |  | conserved hypothetical protein | 0.0.1 |
| RL0944 |  | putative LemA like outer membrane protein | 4.1.4 |
| RL1005 |  | conserved hypothetical protein | 0.0.1 |
| RL1006 | *ribA* | putative riboflavin biosynthesis protein | 3.2.13 |
| RL1087 |  | conserved hypothetical protein | 0.0.1 |
| RL1088 |  | hypothetical protein | 0.0.0 |
| RL1089 |  | conserved hypothetical protein | 0.0.1 |
| RL1090 |  | putative transmembrane protein | 4.1.1 |
| RL1097 | *hsdSch* | putative type I restriction enzyme specificity subunit | 2.2.3 |
| RL1113 |  | putative LuxR/GerE family transcriptional regulator | 6.3.11 |
| RL1115 |  | hypothetical protein | 0.0.0 |
| RL1314 |  | hypothetical exported protein | 0.0.0 |
| RL1315 | *zwf2* | putative glucose-6-phosphate 1-dehydrogenase | 3.5.6 |
| RL1471 |  | putative transmembrane protein | 4.1.1 |
| RL1472 |  | putative transmembrane protein | 4.1.1 |
| RL1473 | *ecfD* | putative RNA polymerase ECF sigma factor (domain COG4941) ECF42 gene organisation | 6.2.1 |
| RL1474 | *dgpfD1* | conserved hypothetical protein with DGPF domain (anti-sigma factor), ECF42 gene organisation with 3 dgfp genes | 6.2.2 |
| RL1475 | *dgpfD2* | conserved hypothetical protein with DGPF domain (anti-sigma factor), ECF42 gene organisation with 3 dgfp genes | 6.2.2 |
| RL1476 | *dgpfD3* | conserved hypothetical protein with DGPF domain (anti-sigma factor), ECF42 gene organisation with 3 dgfp genes | 6.2.2 |
| RL1477 |  | putative transmembrane protein | 4.1.1 |
| RL1570 |  | putative transmembrane protein | 4.1.1 |
| RL1636 |  | conserved hypothetical protein | 0.0.2 |
| RL2083 |  | putative acetyltransferase | 3.3.15 |
| RL2208 |  | putative hydrolase | 3.3.15 |
| RL2320 |  | putative MarR family transcriptional regulator | 6.3.7 |
| RL2321 |  | putative transmembrane protein | 4.1.1 |
| RL2322 |  | conserved hypothetical protein | 0.0.2 |
| RL2323 |  | putative GFO/IDH/MocA dehydrogenase | 3.3.15 |
| RL2400 |  | putative MarC (multiple antibiotic resistance) family transmembrane protein, they may be transporters | 4.1.1 |
| RL2450 |  | putative permease component of ABC transporter CUT2 | 1.5.3 |
| RL2524 |  | conserved hypothetical protein | 0.0.2 |
| RL2525 |  | conserved hypothetical protein | 0.0.2 |
| RL2543 |  | conserved hypothetical protein | 0.0.2 |
| RL2544 | *sdaA* | putative L-serine dehydratase | 3.3.4 |
| RL2545 |  | conserved hypothetical protein | 0.0.1 |
| RL2546 |  | putative transmembrane acyltransferase | 3.3.15 |
| RL2547 | *rpiB* | putative ribose-5-phosphate isomerase B | 3.3.9 |
| RL2548 |  | putative transmembrane protein | 4.1.1 |
| RL2549 |  | putative transmembrane protein | 4.1.1 |
| RL2550 |  | putative MerR family transcriptional regulator | 6.3.12 |
| RL2551 |  | putative transmembrane cationic transporter, MgtE family | 1.5.2 |
| RL2552 | *def2* | putative peptide deformylase | 2.2.10 |
| RL2610 |  | putative GntR family transcriptional regulator | 6.3.3 |
| RL2642 |  | conserved hypothetical protein | 0.0.2 |
| RL2713 |  | putative SBP of ABC transporter FeCT ferrisiderophore binding | 1.5.2 |
| RL2714 |  | putative permease component of ABC transporter FeCT iron transporter | 1.5.2 |
| RL2715 |  | putative ATP-binding component of ABC transporter FeCT iron transporter | 1.5.2 |
| RL2736 | *maiA* | putative maleylacetoacetate isomerase | 3.4.2 |
| RL2737 | *ohr1* | putative organic hydroperoxide resistance protein | 1.6.1 |
| RL2782 |  | putative transmembrane protein | 4.1.1 |
| RL2783 |  | putative transmembrane protein | 4.1.1 |
| RL2784 |  | putative LysR family transcriptional regulator | 6.3.6 |
| RL2785 |  | putative short-chain dehydrogenase/reductase | 3.3.15 |
| RL2818 | *fnrN* | putative FNR/CRP family transcriptional regulator, 100% id to VF39 FnrN | 6.3.0 |
| RL2819 |  | hypothetical protein | 0.0.0 |
| RL2825 |  | putative inosine-uridine preferring nucleoside hydrolase | 3.3.17 |
| RL2826 | *cobE* | putative cobalamin synthesis protein | 3.2.3 |
| RL2827 |  | conserved hypothetical protein | 0.0.2 |
| RL2899 |  | putative ATP-binding component of ABC transporter CUT1 | 1.5.3 |
| RL2910 |  | putative TetR family transcriptional regulator | 6.3.8 |
| RL2935 |  | conserved hypothetical protein | 0.0.2 |
| RL2995 |  | conserved hypothetical protein | 0.0.1 |
| RL2996 |  | conserved hypothetical protein | 0.0.1 |
| RL3044 | *ctaD2* | putative cytochrome c oxidase subunit I | 3.5.1 |
| RL3053 |  | conserved hypothetical protein | 0.0.2 |
| RL3054 |  | conserved hypothetical protein | 0.0.2 |
| RL3095 | *arcB1* | putative ornithine cyclodeaminase | 3.4.2 |
| RL3160 |  | conserved hypothetical protein | 0.0.1 |
| RL3233 |  | putative glyoxalase/bleomycin resistance protein/dioxygenase | 3.3.15 |
| RL3234 | *lppE* | putative lipoprotein LppE, ECF operon with ECF26 gene organisation (lpp, ecf, asf) | 4.1.3 |
| RL3235 | *ecfE* | putative RNA polymerase ECF sigma factor, EcfE, ECF26 gene organisation (lpp, ecf, asf) | 6.2.1 |
| RL3268 | *flaF2* | putative flagellin protein | 1.1.1 |
| RL3350 |  | putative SBP of ABC transporter FeT | 1.5.0 |
| RL3351 |  | putative permease component of ABC transporter FeT | 1.5.0 |
| RL3352 |  | putative permease component of ABC transporter FeT | 1.5.0 |
| RL3420 |  | putative helicase/glycosylase | 2.2.3 |
| RL3421 |  | conserved hypothetical protein | 0.0.2 |
| RL3466 |  | putative Xaa-Pro dipeptidase | 2.1.4 |
| RL3467 |  | conserved hypothetical protein | 0.0.1 |
| RL3488 |  | putative ATP-binding:ATP-binding (ABC:ABC) component of ABC transporter Export | 1.5.5 |
| RL3492 |  | putative transmembrane transporter protein | 1.5.0 |
| RL3493 |  | conserved hypothetical protein | 0.0.2 |
| RL3498 |  | conserved hypothetical protein | 0.0.1 |
| RL3499 |  | conserved hypothetical protein | 0.0.2 |
| RL3508 | *lppF* | conserved hypothetical exported protein, tetratricopeptide TPR_2 repeat protein, lipoprotein?, LppF, ECF26 gene organisation (lpp, ecf, asf) | 4.1.3 |
| RL3509 | *ecfF* | putative RNA polymerase ECF sigma factor, EcfF, ECF26 gene organisation (lpp, ecf, asf) | 6.2.1 |
| RL3510 | *asfF* | putative transmembrane anti-sigma factor, with COG5662, AsfF, ECF26 gene organisation (lpp, ecf, asf) | 6.2.2 |
| RL3511 |  | putative dehydrogenase/oxidoreductase | 3.3.15 |
| RL3512 |  | conserved hypothetical protein | 0.0.1 |
| RL3550 |  | putative glycosylltransferase | 4.1.4 |
| RL3551 |  | putative exported lipase/esterase | 3.4.4 |
| RL3565 |  | putative sensory box HD domain protein | 6.0.0 |
| RL3595 |  | putative LacI family transcriptional regulator (repressor) | 6.3.5 |
| RL3611 |  | putative two-component sensor/regulator; transcriptional regulator | 6.1.2 |
| RL3742 |  | putative class A non-specific acid phosphatase (usually in periplasm, active at low pH) has autotransporter domain | 3.3.15 |
| RL3743 |  | putative two-component sensor/regulator; histidine kinase | 6.1.1 |
| RL3744 |  | putative LuxR/GerE family transcriptional regulator | 6.3.11 |
| RL3745 | *braC* | putative SBP of ABC transporter HAAT | 1.5.1 |
| RL3775 |  | putative RND family efflux transporter (membrane fusion protein) | 1.5.5 |
| RL3776 |  | putative lipase | 3.4.4 |
| RL3829 | *exoY* | putative exopolysaccharide production regulator | 6.0.0 |
| RL3830 |  | putative transmembrane protein | 4.1.1 |
| RL3885 | *sitB* | putative ATP-binding component of ABC transporter MZT (S. mel SBP homologue SMc02509 induced by manganese limitation) | 1.5.2 |
| RL3886 | *sitC* | putative permease component of ABC transporter MZT (S. mel SBP homologue SMc02509 induced by manganese limitation) | 1.5.2 |
| RL4026 |  | putative TetR family transcriptional regulator | 6.3.8 |
| RL4027 |  | putative short-chain dehydrogenase/reductase | 3.3.15 |
| RL4028 | *cheB2* | putative two-component sensor/regulator; chemotaxis transcriptional regulator, glutamate methylesterase | 6.1.2 |
| RL4030 | *cheW3* | putative chemotaxis protein | 1.1.1 |
| RL4031 | *mcrA* | putative sensory transducer methyl-accepting chemotaxis protein | 1.1.1 |
| RL4032 | *mcrB* | putative sensory transducer methyl-acccepting chemotaxis protein | 1.1.1 |
| RL4033 | *mcrC* | putative methyl accepting chemotaxis protein | 1.1.1 |
| RL4034 | *cheW2* | putative chemotaxis protein | 1.1.1 |
| RL4035 | *cheA2* | putative two-component sensor/regulator; histidine kinase chemotaxis protein CheA | 6.1.1 |
| RL4036 | *cheY3* | putative two-component sensor/regulator; chemotaxis transcriptional regulator CheY | 6.1.2 |
| RL4100 |  | putative transmembrane protein | 4.1.1 |
| RL4101 |  | putative transmembrane protein | 4.1.1 |
| RL4102 |  | putative transmembrane protein | 4.1.1 |
| RL4103 |  | conserved hypothetical exported protein | 0.0.1 |
| RL4104 |  | putative SBP of ABC transporter Unclass | 1.5.0 |
| RL4105 |  | putative ornithine carbamoyltransferase | 3.1.2 |
| RL4279 | *clpB* | putative chaperone protein ClpB (heat-shock protein) | 1.3.1 |
| RL4359 |  | putative small permease (DctQ-like) component of TRAP transporter (S. mel SBP homologue SMb21353 induced by pyruvic acid, methyl pyruvic acid ) | 1.5.0 |
| RL4360 |  | putative large permease (DctM-like) component of TRAP transporter (S. mel SBP homologue SMb21353 induced by pyruvic acid, methyl pyruvic acid ) | 1.5.0 |
| RL4361 |  | putative GntR family transcriptional regulator | 6.3.3 |
| RL4375 |  | putative permease component of ABC transporter CUT1 | 1.5.3 |
| RL4376 |  | putative permease component of ABC transporter CUT1 | 1.5.3 |
| RL4377 |  | putative SBP of ABC transporter CUT1 | 1.5.3 |
| RL4378 |  | putative LacI family transcriptional regulator (repressor) | 6.3.5 |
| RL4411 |  | putative transmembrane protein | 4.1.1 |
| RL4533 |  | putative acetyltransferase | 3.3.15 |
| RL4534 |  | conserved hypothetical protein | 0.0.2 |
| **Bacteroid (22 genes)** | | |  |
| pRL110416 | *rhaI* | putative rhamnose isomerase involved in competition for nodulation | 3.3.15 |
| pRL110417 |  | putative carboxymuconolactone decarboxylase (CMD) EC:4.1.1.44 domain, involved in protocatechuate catabolism | 1.4.2 |
| pRL110418 | *ecfO* | putative RNA polymerase ECF sigma factor, atypical ECF41 gene organisation | 6.2.1 |
| RL0001 |  | conserved hypothetical protein | 0.0.1 |
| RL0423 |  | putative transmembrane protein | 4.1.1 |
| RL1529 |  | putative Nudix family protein (phosphohydrolases) | 3.3.18 |
| RL1530 |  | putative transmembrane protein | 4.1.1 |
| RL1531 |  | conserved hypothetical protein | 0.0.2 |
| RL1722 |  | conserved hypothetical protein | 0.0.2 |
| RL2036 |  | putative outer membrane transport protein | 4.1.3 |
| RL2219 |  | conserved hypothetical exported protein | 0.0.2 |
| RL2626 |  | putative transmembrane protein | 4.1.1 |
| RL2627 | *murI* | putative glutamate racemase | 4.1.3 |
| RL2628 |  | putative rRNA methylase family protein | 2.2.11 |
| RL2629 |  | putative transmembrane protein | 4.1.1 |
| RL2630 |  | conserved hypothetical exported protein | 0.0.2 |
| RL3146 |  | putative LysR family transcriptional regulator | 6.3.6 |
| RL3147 |  | putative transmembrane transporter protein | 1.5.0 |
| RL3148 |  | putative ArsR family transcriptional regulator | 6.3.13 |
| RL3576 |  | membrane-bound urate hydroxylase PuuD (putative transmembrane protein) | 4.1.1 |
| RL3819 |  | putative FNR/CRP family transcriptional regulator | 6.3.0 |
| RL4246 |  | putative GFO/IDH/MocA family oxidoreductase | 3.3.15 |

Gene clusters are colored (color is arbitrary and has no significance).

**Table S10.** INSeq classification of genes involved in motility and chemotaxis.

| Details | Name | Gene | Clusters | Input1 | Rhizosphere | Root | Input2 | Nodule bacteria | Bacteroids | CONCLUSION |
| --- | --- | --- | --- | --- | --- | --- | --- | --- | --- | --- |
| methyl-accepting chemotaxis protein | *mcpG* | pRL100403 |  | NE | NE | NE | NE | NE | NE |  |
| flagellar filament | *flaE* | pRL110518 |  | NE | NE | NE | NE | NE | NE |  |
| methyl-accepting chemotaxis protein | *mcpR* | pRL120056 |  | NE | NE | NE | NE | NE | NE |  |
| flagellar hook-filament junction | *flgL2* | pRL120062 | flg2 | NE | NE | NE | NE | NE | NE |  |
| flagellar hook-filament junction | *flgK2* | pRL120063 | flg2 | NE | NE | NE | NE | NE | NE |  |
| flagellar hook complex | *flgE2* | pRL120064 | flg2 | NE | NE | NE | NE | NE | NE |  |
| methyl-accepting chemotaxis protein | *mcpY2* | pRL120068 |  | NE | NE | NE | NE | NE | NE |  |
| methyl-accepting chemotaxis protein | *mcpC* | pRL120312 |  | NE | NE | NE | NE | NE | NE |  |
| methyl-accepting chemotaxis protein | *mcpB* | pRL120683 |  | NE | NE | NE | NE | NE | NE |  |
| methyl-accepting chemotaxis protein | *mcpS* | pRL80031 |  | NE | NE | NE | NE | NE | NE |  |
| methyl-accepting chemotaxis protein | *mcpX* | RL0426 |  | NE | NE | NE | NE | NE | NE |  |
| methyl-accepting chemotaxis protein | *hemAT (previously mcpH)* | RL0429 |  | NE | NE | NE | NE | NE | NE |  |
| methyl-accepting chemotaxis protein | *mcpT* | RL0455 |  | NE | NE | NE | NE | AD | NE | Advantaged nodule-bacteria-specific |
| methyl-accepting chemotaxis protein | *mcpE* | RL0564 |  | NE | NE | NE | NE | NE | NE |  |
| methyl-accepting chemotaxis protein | *icpA (previously hemAT)* | RL0685 | che1 | NE | NE | NE | NE | ES | ES | Nodule-specific |
| chemotaxis response-regulator system | *cheX1/cheS* | RL0686 | che1 | NE | NE | NE | NE | ES | ES | Nodule-specific |
| chemotaxis response-regulator system | *cheY1* | RL0687 | che1 | NE | NE | NE | NE | ES | ES | Nodule-specific |
| chemotaxis response-regulator system | *cheA1* | RL0688 | che1 | NE | NE | NE | NE | ES | ES | Nodule-specific |
| chemotaxis response-regulator system | *cheW1* | RL0689 | che1 | NE | NE | NE | NE | ES | ES | Nodule-specific |
| MCP-associated proteins | *cheR1* | RL0690 | che1 | NE | NE | NE | NE | ES | ES | Nodule-specific |
| MCP-associated proteins | *cheB1* | RL0691 | che1 | NE | NE | NE | NE | ES | ES | Nodule-specific |
| chemotaxis response-regulator system | *cheY2* | RL0692 | che1 | NE | NE | NE | NE | ES | ES | Nodule-specific |
| MCP-associated proteins | *cheD* | RL0693 | che1 | NE | NE | NE | NE | ES | ES | Nodule-specific |
|  |  | RL0694 |  | NE | NE | NE | NE | NE | NE |  |
| flagellar basal body complex | *fliF* | RL0695* | flg1 | NE | NE | NE | NE | DE | ES/DE | Nodule-specific |
| transcriptional regulation | *visN* | RL0696* | flg1 | AD | NE | NE | NE | ES/DE | ES/DE | Nodule-specific |
| transcriptional regulation | *visR* | RL0697* | flg1 | NE | NE | NE | NE | ES/DE | ES/DE | Nodule-specific |
|  |  | RL0698 | flg1 | NE | NE | NE | NE | NE | NE |  |
| flagellar export apparatus | *flhB* | RL0699 | flg1 | NE | NE | NE | NE | ES | ES | Nodule-specific |
| flagellar motor switch (rotor) complex | *fliG* | RL0700 | flg1 | NE | NE | NE | NE | NE | NE |  |
| flagellar motor switch (rotor) complex | *fliN* | RL0701* | flg1 | NE | NE | NE | N/A | N/A | N/A |  |
| flagellar motor switch (rotor) complex | *fliM* | RL0702* | flg1 | NE | NE | NE | NE | ES/DE | ES/DE | Nodule-specific |
| flagellar motor stator complex | *motA* | RL0703* | flg1 | NE | NE | NE | NE | ES/DE | ES | Nodule-specific |
| basal body proximal rod | *flgF* | RL0704* | flg1 | NE | NE | NE | NE | ES/DE | ES | Nodule-specific |
| flagellar export apparatus | *fliI* | RL0705* | flg1 | NE | NE | NE | NE | ES/DE | ES/DE | Nodule-specific |
|  |  | RL0706 | flg1 | NE | NE | NE | NE | NE | NE |  |
| basal body proximal rod | *flgB* | RL0707* | flg1 | NE | NE | NE | NE | ES/DE | ES/DE | Nodule-specific |
| basal body proximal rod | *flgC* | RL0708* | flg1 | NE | NE | NE | NE | ES/DE | ES/DE | Nodule-specific |
| basal body proximal rod | *fliE* | RL0709* | flg1 | N/A | N/A | N/A | N/A | N/A | N/A |  |
| flagellar basal body complex | *flgG* | RL0710* | flg1 | NE | NE | NE | NE | ES/DE | ES/DE | Nodule-specific |
| flagellar assembly apparatus | *flgA* | RL0711 | flg1 | NE | NE | NE | NE | NE | NE |  |
| flagellar basal body complex | *flgI* | RL0712 | flg1 | NE | NE | NE | NE | ES | ES | Nodule-specific |
| flagellar motor stator complex | *motE* | RL0713 | flg1 | NE | NE | NE | NE | DE | DE | Nodule-specific |
| flagellar basal body complex | *flgH* | RL0714 | flg1 | NE | NE | NE | NE | DE | DE | Nodule-specific |
| flagellar basal body complex | *fliL* | RL0715 | flg1 | NE | AD | AD | NE | DE | DE | Nodule-specific & Advantaged rhiz and root-colonised |
| flagellar export apparatus | *fliP* | RL0716* | flg1 | NE | AD | AD | NE | DE | ES/DE | Nodule-specific & Advantaged rhiz and root-colonised |
|  |  | RL0717 | flg1 | NE | AD | AD | NE | NE | NE | Advantaged rhiz and root-colonised |
| flagellar filament | *flaA* | RL0718 | flg1 | NE | NE | NE | NE | NE | NE |  |
| flagellar filament | *flaB* | RL0719 | flg1 | NE | NE | NE | NE | NE | NE |  |
| flagellar filament | *flaC* | RL0720 | flg1 | NE | NE | NE | NE | NE | NE |  |
| flagellar filament | *flaD* | RL0721 | flg1 | NE | NE | NE | NE | NE | NE |  |
|  |  | RL0722 | flg1 | NE | NE | NE | NE | NE | NE |  |
| flagellar motor stator complex | *motB* | RL0723* | flg1 | NE | NE | NE | NE | DE | ES/DE | Nodule-specific |
| flagellar motor stator complex | *motC* | RL0724* | flg1 | NE | NE | NE | NE | DE | ES/DE | Nodule-specific |
| flagellar hook complex | *fliK (previously motD)* | RL0725* | flg1 | NE | NE | NE | NE | ES/DE | ES/DE | Nodule-specific |
| flagellar assembly apparatus | *sltF* | RL0726 | flg1 | NE | NE | NE | NE | DE | DE | Nodule-specific |
| transcriptional regulation | *rem* | RL0727 | flg1 | NE | NE | NE | NE | DE | DE | Nodule-specific |
| flagellar hook complex | *flgE1* | RL0728* | flg1 | NE | AD | AD | NE | DE | ES/DE | Nodule-specific & Advantaged rhiz and root-colonized |
| flagellar hook-filament junction | *flgK1* | RL0729* | flg1 | NE | NE | NE | NE | ES/DE | ES/DE | Nodule-specific |
| flagellar hook-filament junction | *flgL1* | RL0730* | flg1 | NE | NE | NE | NE | ES/DE | DE | Nodule-specific |
| biosynthesis regulation | *flaF/flaF1* | RL0731 | flg1 | NE | NE | NE | NE | DE | DE | Nodule-specific |
| biosynthesis regulation | *flbT* | RL0732 | flg1 | NE | NE | NE | NE | DE | DE | Nodule-specific |
| flagellar assembly apparatus | *flgD* | RL0733* | flg1 | NE | NE | NE | NE | DE | ES/DE | Nodule-specific |
| flagellar export apparatus | *fliQ* | RL0734 | flg1 | N/A | N/A | N/A | N/A | N/A | N/A |  |
| flagellar export apparatus | *flhA* | RL0735* | flg1 | NE | NE | NE | NE | DE | ES/DE | Nodule-specific |
| flagellar export apparatus | *fliR* | RL0736 | flg1 | NE | NE | NE | NE | NE | NE |  |
| methyl-accepting chemotaxis protein | *mcpZ* | RL0757 |  | NE | NE | NE | NE | NE | NE |  |
| methyl-accepting chemotaxis protein | *mcpI* | RL0758 |  | NE | NE | NE | NE | NE | NE |  |
| methyl-accepting chemotaxis protein | *mcpJ* | RL0949 |  | NE | NE | NE | NE | NE | NE |  |
| methyl-accepting chemotaxis protein | *mcpK* | RL0972 |  | NE | NE | NE | NE | NE | NE |  |
| methyl-accepting chemotaxis protein | *mcpL* | RL1065 |  | NE | NE | NE | NE | NE | NE |  |
| methyl-accepting chemotaxis protein | *mcpM* | RL1318 |  | NE | NE | NE | NE | NE | NE |  |
| methyl-accepting chemotaxis protein | MCP putative | RL1386 |  | NE | NE | NE | NE | NE | NE |  |
| methyl-accepting chemotaxis protein | MCP putative | RL1447 |  | NE | NE | NE | NE | NE | NE |  |
| transcriptional regulation | *flaR1* | RL2250 |  | NE | NE | NE | NE | NE | NE |  |
| transcriptional regulation | *flaR2* | RL2251 |  | NE | NE | NE | NE | NE | NE |  |
| basal body proximal rod | *flgC* hypothetical | RL2575* |  | NE | NE | NE | NE | ES/DE | ES/DE | Nodule-specific |
| methyl-accepting chemotaxis protein | *mcpD* | RL2683 |  | NE | NE | NE | NE | NE | NE |  |
| methyl-accepting chemotaxis protein | *mcpN* | RL2931 |  | NE | NE | NE | NE | NE | NE |  |
| flagellar filament | *flaH/flaF2* | RL3268 |  | NE | NE | NE | NE | AD | NE | Advantaged nodule-bacteria-specific |
| chemotaxis response-regulator system | *cheW3* | RL3289 |  | NE | NE | NE | NE | NE | NE |  |
| flagellar assembly apparatus | *flgJ* | RL3927 |  | AD | NE | NE | NE | NE | NE |  |
| methyl-accepting chemotaxis protein | *mcpO* | RL3985 |  | NE | NE | NE | NE | NE | NE |  |
| flagellar motor stator complex | *motB* putative | RL4023 |  | NE | NE | NE | NE | NE | NE |  |
| MCP-associated proteins | *cheB2* | RL4028 | che2 | NE | NE | NE | NE | AD | NE | Advantaged nodule-bacteria-specific |
| MCP-associated proteins | *cheR2* | RL4029 | che2 | NE | AD | AD | NE | AD | NE | Advantaged nodule-bacteria, rhiz and root-colonised |
| chemotaxis response-regulator system | *cheW2b* | RL4030 | che2 | NE | NE | NE | NE | AD | NE | Advantaged nodule-bacteria-specific |
| methyl-accepting chemotaxis protein | *mcrA* | RL4031 | che2 | NE | NE | NE | NE | AD | NE | Advantaged nodule-bacteria-specific |
| methyl-accepting chemotaxis protein | *mcrB* | RL4032 | che2 | NE | NE | NE | NE | AD | NE | Advantaged nodule-bacteria-specific |
| methyl-accepting chemotaxis protein | *mcrC* | RL4033 | che2 | NE | NE | NE | NE | AD | NE | Advantaged nodule-bacteria-specific |
| chemotaxis response-regulator system | *cheW2a* | RL4034 | che2 | NE | NE | NE | NE | AD | NE | Advantaged nodule-bacteria-specific |
| chemotaxis response-regulator system | *cheA2* | RL4035 | che2 | NE | NE | NE | NE | AD | NE | Advantaged nodule-bacteria-specific |
| chemotaxis response-regulator system | *cheY3* | RL4036 | che2 | NE | NE | NE | NE | AD | NE | Advantaged nodule-bacteria-specific |
| methyl-accepting chemotaxis protein | *mcpP* | RL4277 |  | NE | NE | NE | NE | NE | NE |  |
| chemotaxis response-regulator system | *cheW4** | RL4386 |  | NE | NE | NE | NE | DE | ES/DE | Nodule-specific |
| methyl-accepting chemotaxis protein | *mcpQ* | RL4387 |  | NE | NE | NE | NE | NE | NE |  |
| flagellar motor switch (rotor) complex | *fliY* | RL4634 |  | NE | NE | NE | NE | NE | NE |  |
| flagellar filament | *flaG* | RL4729 |  | NE | NE | NE | NE | NE | NE |  |

*** manually curated**

**Table S11.** INSeq-identified metabolic phenotypes in Rlv3841.

| **EC^a^** | **Gene^b^** | **Minimal Medium^c^** | **Rhizo-sphere** | **Root colonized** | **Nodule bacteria** | **Bacteroids** | **Remarks** |
| --- | --- | --- | --- | --- | --- | --- | --- |
| **Amino acid biosynthesis** | | | | | | | |
| **Glutamate biosynthesis** | | | | | | | |
| 1.4.1.13 | RL4084 (*gltD*)  RL4085 (*gltB*) | ES  ES | ES  ES | ES ES | ES ES | ES ES |  |
| **Glutamine biosynthesis** | | | | | | | |
| 6.3.1.2 | RL0755,  RL1466,  RL2392 (*glnA*, GSI),  RL3346,  RL3549 (*glnII*, GSII),  pRL110554 (*glnT*, GSIII) |  |  |  | DE | ES |  |
| **Aspartate biosynthesis** | | | | | | | |
| 2.6.1.1 | RL3443 (*aatA*),  pRL100431 (*aatA2*),  pRL120409 (*aatA3*), | ES | ES | ES | ES | ES |  |
| **Asparagine biosynthesis** | | | | | | | |
| 6.3.5.6 | RL0225^1^,  pRL110427^1^,  pRL120136^1^,  RL2076 (*gatA*)^1,2^  RL2082 (*gatB*)^2^  RL2075 (*gatC*)^2^ | ES  ES  ES | ES  ES  ES | ES  DE  ES | ES  ES  ES | ES  ES  ES | ^1^All subunit A  ^2^ES in TY |
| 6.3.5.4 | pRL100163 |  |  |  |  |  |  |
| **Alanine biosynthesis** | | | | | | | |
| 1.4.1.1 | RL1966 (*aldA*) |  |  |  |  |  |  |
| **Histidine biosynthesis** | | | | | | | |
| 2.4.2.17 | RL0878 (*hisZ*)^1^  RL0879 (*hisG*) | ES  ES | ES  ES | ES  ES | ES  ES | ES  ES | ^1^Regulatory subunit |
| 6.3.1.31 | RL0041 | ES | ES | ES | ES | ES |  |
| 6.5.4.19 | RL2532 (*hisI*) | ES | DE | ES | DE | ES |  |
| 5.3.1.16 | RL0043 (*hisA*) | ES | ES | ES | ES | ES |  |
| 4.3.2.10 | RL0042 (*hisF*)  RL0819 (*hisF2*),  RL0046 (*hisH*)^1^  RL0820 (*hisH2*) | ES  ES | ES  ES | ES  ES  DE | ES  ES  ES  ES | ES  ES  ES  ES | ^1^probably^d^ ES in TY |
| 4.2.1.19 | RL0048 (*hisB*)^1^ | ES | ES | ES | ES | ES | ^1^probably^d^ ES in TY |
| 2.6.1.9 | RL4338 (*hisC1*)  RL4629 (*hisC2*)  RL1482 (*hisC3*) | DE | DE | ES | ES | ES |  |
| 3.1.3.15 | Not annotated | | | | | | |
| 1.1.1.23 | RL0613 (*hisD*) | ES | ES | ES | DE | DE |  |
| **Lysine biosynthesis** | | | | | | | |
| 2.7.2.4 | RL4284 (*aspK*)^1,2^ | ES ^3^ | ES | ES | ES | ES | ^1^Original annotation is *aspC*, reannotated as *aspK* (aspartate kinase) to avoid confusion with the asparate aminotransferase gene *aspC* (RL4471), an aromatic amino acid transaminase  ^2^probably^d^ ES in TY  ^3^DE in glucose |
| 1.2.1.11 | RL4715 (*asd*) | ES | DE | ES | ES | ES |  |
| 4.3.3.7 | RL1502 (*mosA*)^1^,  RL1811 (*dapA1*),  Rl3247,  RL3594,  RL4423,  RL4484 (*dapA2*),  pRL90100,  pRL110028,  pRL120004,  pRL120328,  pRL120528,  pRL120776 | DE | ES | ES | ES  AD | ES | ^1^Also involved in rhizopine biosynthesis |
| 1.17.1.8 | RL0180 (*dapB1*),  RL3002 (*dapB2*) | ES | ES | ES | ES | ES |  |
| 2.3.1.117 | RL0437 (*dapD*) | ES | ES | ES | ES | ES |  |
| 2.6.1.17 | RL0549 (*argD*)^1,2^ | ES | ES | ES | ES | ES | ^1^Annotated as *gabT2*  ^2^Also involved in Arg biosynthesis |
| 3.5.1.18 | RL0436 (*dapE*) | ES | ES | ES | ES | ES |  |
| 5.1.1.7 | RL4545 (*dapF*)^1^ | ES | ES | ES | ES | ES | ^1^ES in TY |
| 4.1.1.20 | RL4325 (*lysA*)^1^ | ES ^2^ | DE | DE | ES | ES | ^1^probably^d^ ES in TY  ^2^DE in glucose |
| **Arginine biosynthesis** | | | | | | | |
| 2.3.1.1 | RL4296 (*argJ*)^1^ | DE | ES | DE | ES | ES | ^1^Also catalysing reversible ornithine acetylation. Probably missing actual *argA* annotation. |
| 2.7.2.8 | RL0445 (*argB*) |  |  |  |  |  |  |
| 1.2.1.38 | RL1668 (*argC*) |  | ES | ES | ES |  |  |
| 2.6.1.11 | RL0549 (*argD*)^1,2^ | ES | ES | ES | ES | ES | ^1^Annotated as *gabT2*  ^2^Also involved in Lys biosynthesis |
| 2.3.1.35  or  3.5.1.16 | RL4296 (*argJ*)  pRL120633 (*argE*) | DE | ES | DE | ES | ES |  |
| 2.1.3.3 | RL0550 (*argF*) | ES | ES | ES | ES | ES |  |
| 6.3.4.5 | RL2987 (*argG1*),  RL4515 (*argG2*) |  |  |  | DE | ES |  |
| 4.3.2.1 | RL4323 (*argH*) | ES | ES | ES | ES | ES |  |
| **Proline biosynthesis** | | | | | | | |
| 2.7.2.11 | RL4682 (*proB*) | ES | ES | ES | ES | ES |  |
| 1.2.1.41 | RL4683 (*proA*) | ES | ES | ES | ES | ES |  |
| 1.5.1.2 | RL3460 (*proC1*),  pRL120421 (*proC2*) | ES ^1^ |  |  | ES | DE | ^1^DE in succinate |
| **Aromatic amino acid biosynthesis** | | | | | | | |
| Chorismate biosynthesis | | | | | | | |
| 2.5.1.54 | RL2686 (*aroG*),  RL2692 |  |  |  |  |  |  |
| 4.2.3.4 | RL4352 (*aroB*) | ES | ES | ES | DE | ES |  |
| 4.2.1.10 | RL2090^1^,  pRL110076 | ES | ES | ES | ES | ES | ^1^ES in TY |
| 1.1.1.25 | RL0003,  RL2845 (*aroE*),  RL2847  pRL110081 | ES | ES | ES | ES | ES |  |
| 2.7.1.71 | RL4353 (*aroK*)^1^ | ES | ES | ES | DE | ES | ^1^probably^d^ ES in TY |
| 2.5.1.19 | RL0108 (*aroA1*),  RL0945 (*aroA2*) | DE | DE | DE | ES  ES | ES |  |
| 4.2.3.5 | RL1007 (*aroC*) | ES | ES | ES | ES |  |  |
| Tryptophane biosynthesis | | | | | | | |
| 4.1.3.27 | RL3521 (*trpE*) | ES | DE | DE | ES | ES |  |
| 2.4.2.18 | RL2493 (*trpD*) | ES | DE | ES | ES | ES |  |
| 5.2.1.24 | RL0020 | ES | DE | ES | ES | ES |  |
| 4.1.1.48 | RL2494 (*trpC*) | ES | DE | DE | ES | ES |  |
| 4.2.1.20 | RL0022 (*trpA*)  RL0021 (*trpB*) | ES  ES | ES  DE | ES  ES | ES  ES | ES  ES |  |
| Tyrosine and Phenylalanine biosynthesis  (The whole biosynthesis pathways are unclear, two routes may exist. Both start at prephenate and involve one transamination reaction and either a hydrolysation/decarboxylation (Phe) or oxidation/decarboxylation (Tyr) reaction. If the transamination reaction takes place first, arogenate is formed and then converted into either Phe (RL0139) or Tyr (TyrC). The same enzymes would also act on prephenate, forming either Phenylpyruvate (RL0139) or 4-hydroxiphenylpyruvate (TyrC) which then would be transaminated to Phe or Tyr, respectively. Besides aromatic amino acid transaminases, also other aspartate-accepting transaminases such as *aatA1-3* or *hisC1-3* could perform the transamination. | | | | | | | |
| 5.4.99.5 | RL4548^1^ | ES | ES | ES | ES | ES | ^1^probably^d^ ES in TY |
| 2.6.1.57 | RL4471 (*aspC*),  pRL100056 |  |  |  | DE |  | Aromatic amino acid transaminases |
| 4.2.1.51 | RL0139 | ES | DE | DE | ES | ES | Phe biosynthesis |
| 1.14.16.1 | RL1860 (*phhA*) |  |  |  |  |  | Phe to Tyr conversion |
| 1.3.1.12/43 | RL4337 (*tyrC*) | DE | DE | ES | ES | ES | Tyr biosynthesis |
| **Serine and Glycine biosynthesis** | | | | | | | |
| 1.1.1.95 | RL3960,  pRL120588 | ES ^1^ | ES | ES | ES | ES | ^1^NE in glucose |
| 2.6.1.52 | RL3961 (*serC*)^1^ | ES | ES | ES | ES | ES | ^1^probably^d^ ES in TY |
| 3.1.3.3 | RL3250 (*serB*) | ES | ES | ES | ES | ES | Ser biosynthesis |
| 2.1.2.1 | RL1620 (*glyA*) | ES | DE | DE | ES | ES | Gly biosynthesis |
| Threonine Glycine interconversion | | | | | | | |
| 4.1.2.48 | RL4086 (*ltaA*) |  |  |  |  |  |  |
| **Threonine biosynthesis** | | | | | | | |
| 2.7.2.4 | RL4284 (*aspK*)^1^ | ES ^2^ | ES | ES | ES | ES | ^1^cf. Lys biosynthesis  ^2^DE in glucose |
| 1.2.1.11 | RL4715 (*asd*) | ES | DE | ES | ES | ES |  |
| 1.1.1.3 | RL2097 (*hom1*),  pRL80071 (*hom2*) | ES | ES | ES | DE | ES |  |
| 2.7.1.39 | RL1031 (*thrB*)^1^ | ES | ES | ES | ES | ES | ^1^probably^d^ ES in TY |
| 4.2.3.1 | RL1062 (*thrC*)^1^ | ES | ES | ES | ES | ES | ^1^ES in TY |
| **Leucine, Isoleucine, and Valine biosynthesis** | | | | | | | |
| 2.2.1.6 | RL3244 (*ilvH*)  RL3245 (*ilvI*),  RL3338 (*ilvG*)^1^,  pRL100055^1^,  pRL120271 (*budB*)^1^ | ES | ES  AD | ES | ES  ES  ES | ES  ES | ^1^Orphan large subunits only |
| 1.1.1.86 | RL3205 (*ilvC*) | ES | ES | ES | ES | DE |  |
| 4.2.1.9 | RL1803 (*ilvD1*),  RL4421 (*ilvD5*),  pRL120238 (*ilvD*) | ES ^1^ | ES | ES | ES | ES | ^1^DE in glucose |
| 2.3.3.13 | RL3513 (*leuA1*),  RL1538 (*leuA2*) | ES ^1^ | ES | ES | ES | ES | Val specific  ^1^DE in glucose |
| 4.2.1.33 | RL4555  RL4705 (*leuD*) | ES ^1^ |  | DE | ES | ES | Val and Ile biosynthesis  ^1^NE in glucose |
| 1.1.1.85 | RL4707 (*leuB*) | DE | DE | ES | DE | ES | Val and Ile biosynthesis |
| 2.6.1.42 | RL1326 (*ilvE*)  RL3200 (*ilvE1*) |  |  |  |  |  | Leu, Ile, and Val biosynthesis |
| **Cysteine biosynthesis** | | | | | | | |
| Sulfite reduction | | | | | | | |
| 1.8.1.2 | RL2274  RL2290 (*cysI*) | ES ^1^ |  |  | ES | ES | ^1^DE in glucose |
| Starting from Serine | | | | | | | |
| 2.3.1.30 | RL2209 (*cysE1*),  RL4152 (*cysE2*) | DE |  | DE | ES |  |  |
| 2.5.1.47 | RL0340 (*cysK*),  RL1979 (*cysB*) |  |  |  |  |  |  |
| Or Starting from 3-Phosphoglycerate, via O-Phosphoserine | | | | | | | |
| 1.1.1.95 | RL3960,  pRL120588 | ES ^1^ | ES | ES | ES | ES | ^1^NE in glucose |
| 2.6.1.52 | RL3961 (*serC*)^1^ | ES | ES | ES | ES | ES | ^1^probably^d^ ES in TY |
| 2.5.1.47 | RL0340 (*cysK*),  RL1979 (*cysB*) |  |  |  |  |  |  |
| **Methionine biosynthesis** | | | | | | | |
| Homoserine biosynthesis (shared with Threonine biosynthesis) | | | | | | | |
| 2.7.2.4 | RL4284 (*aspK*)^1^ | ES ^2^ | ES | ES | ES | ES | ^1^cf. Lys biosynthesis  ^2^DE in glucose |
| 1.2.1.11 | RL4715 (*asd*) | ES | DE | ES | ES | ES |  |
| 1.1.1.3 | RL2097 (*hom1*),  pRL80071 (*hom2*) | ES | ES | ES | DE | ES |  |
| Homocysteine biosynthesis via Succinyl-homoserine | | | | | | | |
| 2.3.1.46 | RL4606 (*metA*),  pRL100137 (*metX*) | ES | ES | ES | ES | ES |  |
| 2.5.1.- | RL0554 (*metZ*) | ES | ES | ES | ES | ES |  |
| Or instead via O-Acetyl-homoserine | | | | | | | |
| 2.3.1.31 | RL4606 (*metA*) | ES | ES | ES | ES | ES |  |
| 2.5.1.49 | RL1679 (*cysD*) |  |  |  |  |  |  |
| Homocysteine to Methionine conversion | | | | | | | |
| 2.1.1.13  or  2.1.1.10 | RL3362 (*metH*)  RL1357 | ES | ES | ES | ES | ES |  |
| **Purine and Pyrimidine biosynthesis** | | | | | | | |
| **Purine biosynthesis** | | | | | | | |
| 2.4.2.14 | RL1546 (*purF*) | ES |  |  | ES | ES |  |
| 6.3.4.13 | RL0947 (*purD*) | DE |  |  | ES | ES |  |
| 2.1.2.2 | RL1595 (*purN*) | DE |  |  | ES | ES |  |
| 6.3.5.3 | RL2608 (*purQ*)  RL2612 (*purL*) | ES |  |  | DE  ES | ES |  |
| 6.3.3.1 | RL1596 (*purM*) | DE |  |  | ES | ES |  |
| 6.3.4.18 | RL4045 (*purK*) | ES |  |  | ES | ES |  |
| 4.4.99.18 | RL4044 (*purE*) | ES |  |  | ES | ES |  |
| 6.3.2.6 | RL2606 (*purC1*),  RL3872 (*purC2*) | ES  DE |  |  | ES | ES |  |
| 4.3.2.2 | RL2601 (*purB*) | ES | ES | ES | ES | ES |  |
| 2.1.2.3  and  3.5.4.10 | RL4722 (*purH*) | ES | DE | DE | ES | DE | IMP biosynthesis |
| 6.3.4.4 | RL3768 (*purA*) | ES | ES | ES | ES | ES |  |
| 4.3.2.2 | RL2601 (*purB*) | ES | ES | ES | ES | ES | AMP biosynthesis |
| 2.7.4.3 | RL1795 (*adk1*)^1^,  pRL100122 (*adk2*) | ES | ES | ES | ES | ES | ADP biosynthesis  ^1^ES in TY |
| 1.17.4.1 | RL1945^1^,  RL4259 (*nrdE*)  RL4258 (*nrdF*) | DE | DE | DE | ES | ES | dADP biosynthesis  ^1^Orphan alpha chain |
| 1.1.1.205 | RL0847 (*guaB*) | ES |  | DE | ES | ES |  |
| 6.3.5.2 | RL0251,  RL0315 (*guaA*),  RL4500 | DE | ES | ES | ES | ES | GMP biosynthesis |
| 2.7.4.8 | RL1563 (*gmk*)^1^ |  |  | ES |  |  | GDP biosynthesis  ^1^ES in TY |
| 1.17.4.1 | RL1945^1^,  RL4259 (*nrdE*)  RL4258 (*nrdF*) | DE | DE | DE | ES | ES | dGDP biosynthesis  ^1^Orphan alpha chain |
| **Pyrimidine biosynthesis** | | | | | | | |
| 6.3.5.5 | RL3411 (*carA*)^1^  RL3419 (*carB*) | ES  ES | ES  ES | ES  ES | ES  ES | ES  ES | Carbamoyl-phosphate biosynthesis  ^1^probably^d^ ES in TY |
| 2.1.3.2 | RL1739 (*pyrB*) | ES | DE | DE | DE | DE |  |
| 3.5.2.3 | RL0493 (*pyrC*),  RL1738 (*pyrC2*),  pRL110419,  pRL120121 | ES  ES | DE | DE | ES  DE | DE  DE |  |
| 1.3.5.2 | RL0572 | ES |  |  | ES | ES |  |
| 2.4.2.10 | RL0501 |  |  |  |  | DE |  |
| 4.1.1.23 | RL0331 (*pyrF*) |  |  |  |  |  | UMP biosynthesis |
| 2.7.4.22 | RL2223 (*pyrH*) | ES | ES | ES | DE | ES |  |
| 2.7.4.6 | RL1580 (*ndk*) |  |  |  |  |  |  |
| 6.3.4.2 | RL2511 (*pyrG*)^1^ | ES | ES | DE | ES | ES | CTP biosynthesis  ^1^ES in TY |
| 3.5.4.13 | RL0555 |  |  |  |  |  |  |
| 2.7.4.9 | RL2476 (*tmk*)^1^ | ES | ES | ES | ES | ES | ^1^ES in TY |
| or | | | | | | | |
| 3.1.3.5 | RL2050 (*surE*),  RL4075 |  | AD |  |  | AD |  |
| 3.5.4.5 | RL0200 (*cdd*) |  |  |  |  |  |  |
| 2.7.1.21 | RL3532 (*tdk*) |  |  |  |  |  |  |
| 2.1.1.45 | RL3256 (*thyA*)^1^ | ES ^2^ | ES | ES | ES | ES | dTMP biosynthesis  ^1^ES in TY  ^2^DE in glucose |
| **Cofactor biosynthesis** | | | | | | | |
| **S-Adenosyl-methionine biosynthesis/regeneration** | | | | | | | |
| 2.5.1.6 | RL0389 (*metK*)^1^ | ES | ES | ES | ES | ES | ^1^probably^d^ ES in TY |
| 2.1.1.37 | RL1932 |  |  |  |  |  |  |
| 3.3.1.1 | RL0031,  RL0860 | ES ^1^ | DE | DE | ES | ES | ^1^DE in glucose |
| 2.1.1.13  or  2.1.1.10 | RL3362 (*metH*)  RL1357 | ES | ES | ES | ES | ES |  |
| **Heme and Vitamin B12 coenzyme biosynthesis** | | | | | | | |
| Uroporphyrinogen III biosynthesis | | | | | | | |
| 2.3.1.37 | RL4379 (*hemA1*)^1^,  pRL90008 (*hemA*)^2^ | ES  DE^3^ | ES | ES | ES | ES | ^1^ES in TY  ^2^probably^d^ ES in TY  ^3^NE in glucose |
| 4.2.1.24 | RL1616 (*hemB*) | ES | ES | ES | DE | DE |  |
| 2.5.1.61 | RL4495 (*hemC*) | ES | ES | ES | DE | ES |  |
| 4.2.1.75 | RL4496 (*hemD*) | ES | ES | ES | DE | ES |  |
| Heme biosynthesis | | | | | | | |
| 4.1.1.37 | RL4742 (*hemE*)^1^ | ES | ES | ES | ES | ES | ^1^ES in TY |
| 1.3.3.3  or  1.3.98.3 | RL3494 (*hemF*),  pRL90023 (*hemN*) | ES | ES | ES | ES | ES |  |
| 1.3.3.4  or  1.3.5.3 | Not annotated | | | | | | |
| 4.99.1.1 | RL4076 (*hemH*)^1^ | ES | ES | ES | DE | ES | ^1^probably^d^ ES in TY |
| 2.5.1.141 | RL1023 (*ctaB*) | DE ^1^ | DE |  | ES | ES | ^1^NE in succinate |
| - | RL1666 (*cox15*) |  |  |  |  |  | HemA biosynthesis |
| 1.14.99.58 | RL3713 |  |  |  |  |  | Biliverdin-IX biosynthesis |
| Vitamin B12 coenzyme biosynthesis | | | | | | | |
| 2.1.1.107 | RL2288 (*cysG2*),  RL2824 (*cobA*) | ES ^1^  ES | ES | ES | ES  ES | ES  ES | ^1^DE in glucose |
| 2.1.1.130 | pRL110629 (*cobI*) | ES | ES | ES | ES | DE |  |
| 1.14.13.83 | pRL110631 (*cobG*) | ES | ES | ES | DE | DE |  |
| 2.1.1.131 | RL2826 (*cobE*),  pRL110628 (*cobJ*) | ES | ES | ES | AD  ES | DE |  |
| 2.1.1.133 | pRL110625 (*cobM*) | ES | ES | ES | ES | DE |  |
| 2.1.1.152 | pRL110632 (*cobF*) | ES | ES | ES | DE | ES |  |
| 1.3.1.106 | pRL110627 (*cobK*) | ES | ES | ES | ES | DE |  |
| 2.1.1.132 | pRL110626 (*cobL*) | ES | ES | ES | ES | DE |  |
| 5.4.99.61 | pRL110630 (*cobH*) | ES | ES | ES | ES | DE |  |
| 6.3.5.9 | RL2823 (*cobB*) | ES | ES | ES | ES | ES |  |
| 6.6.1.2 | RL2830 (*cobN*)  RL4348 (*cobT*)  RL4349 (*cobS*) | ES  ES  ES | ES  DE  ES | ES  ES  ES | ES  ES  ES | ES  ES  ES |  |
| 1.16.8.1 | Not annotated | | | | | | |
| 2.5.1.17 | RL2829 (*cobO*) | ES | ES | ES | ES | ES |  |
| 6.3.5.10 | RL2836 (*cobQ*) | ES | ES | ES | ES | ES |  |
| 6.3.1.10 | RL2821 (*cobD*)  RL2822 (*cobC*) | ES  ES | ES  ES | ES  ES | ES  ES | ES  ES |  |
| 2.7.1.156 | RL2832 (*cobP*) | ES | ES | ES | DE | DE |  |
| 2.7.8.26 | RL2781A (*cobV*) | ES | ES | ES | ES | ES |  |
| **Pantothenate^e^ and Coenzyme A biosynthesis** | | | | | | | |
| 2.1.2.11 | pRL110619,  pRL120360 (*panB*) |  |  |  |  |  |  |
| 1.1.1.169 | RL1813 |  |  |  |  |  |  |
| 6.3.2.1 | pRL120359 (*panC*) |  |  |  |  |  | Pantothenate biosynthesis |
| 2.7.1.33 | RL0040 (*coaA*) | ES | ES | ES | ES | ES |  |
| 6.3.2.5  and  4.1.1.36 | RL0357 (*coaBC*)^1^ | ES | ES | ES | ES |  | ^1^probably^d^ ES in TY |
| 2.7.7.3  or  3.6.1.9 | RL2403^1^  RL2286 |  |  |  | DE | DE | ^1^probably^d^ ES in TY |
| 2.7.1.24 | RL0004^1^ | ES | ES | ES | ES | ES | ^1^ES in TY |
| **Riboflavin and FMN/FAD biosynthesis** | | | | | | | |
| 3.5.4.25 | RL1006 (*ribA*) |  |  |  | AD |  |  |
| 3.5.4.26  and  1.1.1.193 | RL1621 (*ribG*) | ES | ES | ES | ES | ES |  |
| 3.1.3.104 | Not annotated | | | | | | |
| 4.1.99.12 | RL1006 (*ribA*),  RL2726 (*ribA1*) |  |  |  | AD |  |  |
| 2.5.1.78 | RL1632 (*ribH1*)^1^,  RL3153 (*ribH2*) |  | ES |  | DE | DE | ^1^probably^d^ ES in TY |
| 2.5.1.9 | RL1622 (*ribC*) | ES | ES | ES | ES | ES | Riboflavin biosynthesis |
| 2.7.1.26  and  2.7.7.2 | RL0886 (*ribF*)^1^ | ES | DE | ES | ES | DE | FMN and FAD biosynthesis  ^1^probably^d^ ES in TY |
| **Thiamine^f^ biosynthesis**  (Thiazol biosynthesis is not annotated and hence omitted) | | | | | | | |
| 4.1.99.17 | pRL100148 (*thiC*) |  |  |  |  |  |  |
| 2.7.4.7 | pRL110441 (*thiD*) |  |  |  |  |  |  |
| 2.5.1.3 | RL4040 (*thiE2*),  pRL110442 (*thiE*) |  |  |  | ES | ES |  |
| 3.1.3.1  or  3.1.3.100 | RL4713  pRL90033 |  |  |  |  |  |  |
| **Biotin biosynthesis is not annotated** | | | | | | | |
| **Pyridoxal biosynthesis is only partially annotated** | | | | | | | |
| 1.1.1.262 | RL1565 (*pdxA1*)^1^ | ES | ES | ES | ES | ES | ^1^probably^d^ ES in TY |
| 2.6.99.2 | RL3222 (*pdxJ*) |  | ES | ES | ES |  |  |
| 1.4.3.5 | RL1014 (*pdxH*),  RL1184 | ES | ES | ES | ES | ES |  |
| 2.7.1.35 | RL4719 (*pdxK*) |  |  |  |  |  |  |
| 1.1.1.65 | RL0474,  RL3415,  pRL100446 |  | AD | AD |  |  |  |
| **Glutathione biosynthesis and salvage** | | | | | | | |
| 6.3.2.2 | RL0855 (*gshA*) |  | DE | ES | ES | ES |  |
| 6.3.2.3 | RL0338 (*gshB*) |  | DE | DE | ES | ES |  |
| 4.3.2.7 | RL4334,  pRL100236 |  |  |  |  |  |  |
| 3.5.2.9 | RL2446 (*hyuA1*),  RL2462,  pRL100432,  pRL120154 |  |  |  |  | ES |  |
| 3.4.19.13 | RL4050 (*ggt1*),  RL4578 (*ggt2*),  pRL120335 |  |  |  |  |  |  |
| 3.4.11.1  or  3.4.11.2 | RL0222,  RL1571 (*pepA*)  RL1446 (*pepN*) | DE ^1^ | DE | DE | ES | ES | ^1^NE in glucose |
| **Tetrahydrofolate biosynthesis** | | | | | | | |
| 3.5.4.16 | RL2531 (*folE*)  pRL120570 | ES | DE | ES | DE | ES |  |
| 3.1.3.1 | RL2417,  RL4713 |  |  |  |  |  |  |
| 4.1.2.25 | RL2649 (*folB*) | ES | ES | ES | ES | ES |  |
| 2.7.6.3 | RL2648 | ES | ES | ES | ES | ES |  |
| 2.5.1.15 | RL2650 (*folC1*) | ES | ES | ES | ES | ES |  |
| 6.3.2.12/17 | RL0024 (*folC2*)^1^ | ES | ES | ES | ES | ES | ^1^probably^d^ ES in TY |
| 1.5.1.3 | RL3255 (*folA*)^1^ | ES ^2^ | ES | ES | DE | ES | ^1^ES in TY  ^2^DE in glucose |
| **Molybdopterin cofactor biosynthesis** | | | | | | | |
| 4.1.99.22 | RL2711 (*moaA2*)  pRL80034 (*moaA1*) |  |  |  |  |  |  |
| 4.6.1.17 | RL2495 (*moaC*) |  |  |  |  |  |  |
| 2.8.1.12 | RL1584 (*moaE*)  RL1585 (*moaD*) | ES |  |  |  |  |  |
| 2.7.7.75 | RL0934 (*moaB*) |  |  |  | ES | ES |  |
| 2.10.1.1 | RL2496 (*moeA*) |  |  |  |  |  |  |
| 2.7.7.76 | pRL120298 |  |  |  |  |  |  |
| 2.7.7.77 | RL2729 (*mobA*)  RL2730 (*mobB*) |  |  |  |  |  |  |

^a^Listed in the order of reaction steps wherever possible. Immediate subsequent steps are separated by dashed lines.

^b^Homologues are separated by commas to distinguish from subunits.

^c^Minimal medium supplemented with NH_4_ and either glucose or succinate, 21% O_2._

^d^In one of two experiments.

^e^Minimal medium is supplemented with pantothenate.

^f^Minimal medium is supplemented with thiamin.

HMM classifications: ES; essential, GD; growth-defective, NE; neutral, AD; advantaged. Empty cell is NE.

**Table S12.** Strains, plasmids, and primers.

| **Strain, plasmid, or primer** | **Description/ Sequence (5'to 3')** | **Source** |
| --- | --- | --- |
| Strain |  |  |
| Rlv3841 | Streptomycin-resistant derivative of *R. leguminosarum* bv. *viciae* strain 300 | (1) |
| 3841celB | Rlv3841 with chromosomal *celB* under constitutive Ptac promoter | (2) |
| 3841gusA | Rlv3841 with chromosomal *gusA* under constitutive Ptac promoter | (2) |
| DH5α | *Escherichia coli* used for cloning, F- 80*dlacZ* M15 (*lacZYA-argF*) *U169 recA1 endA1hsdR17*(rk-, mk+) *phoAsupE44* -*thi-1 gyrA96 relA1* | Bioline |
| OPS0132 | Rlv3841 mutated in pRL80032 (PCR product of oxp1072/oxp1073 cloned in pK19mob conjugated into Rlv3841) | This work |
| OPS1598 | 3841gusA mutated in RL0688 (*cheA1*) (pOPS0820 conjugated into 3841gusA) | This work |
| OPS1599 | 3841gusA mutated in pRL100199 (*fixB*) (pOPS0822 conjugated into 3841gusA) | This work |
| OPS1600 | 3841gusA mutated in pRL100195 (*nifB*) (pOPS0823 conjugated into 3841gusA) | This work |
| OPS1602 | 3841gusA mutated in pRL90058 (*rmrR*) (pOPS0825 conjugated into 3841gusA) | This work |
| OPS1603 | 3841gusA mutated in pRL120205 (*eryB*) (pOPS0831 conjugated into 3841gusA) | This work |
| OPS1604 | 3841gusA mutated in RL0149 (pOPS0833 conjugated into 3841gusA) | This work |
| OPS1606 | 3841gusA mutated in RL1545 (pOPS0836 conjugated into 3841gusA) | This work |
| OPS1607 | 3841gusA mutated in RL2606 (*purC1*) (pOPS0837 conjugated into 3841gusA) | This work |
| OPS1608 | 3841gusA mutated in RL3549 (*glnII*) (pOPS0838 conjugated into 3841gusA) | This work |
| OPS1609 | 3841gusA mutated in RL3654 (*pssD*) (pOPS0839 conjugated into 3841gusA) | This work |
| OPS1610 | 3841gusA mutated in pRL120198 (pOPS0846 conjugated into 3841gusA) | This work |
| OPS1611 | 3841gusA mutated in RL1496 (*iolR*) (pOPS0849 conjugated into 3841gusA) | This work |
| OPS1612 | 3841gusA mutated in RL3986 (*ruvC*) (pOPS0851 conjugated into 3841gusA) | This work |
| OPS2053 | Rlv3841 mutated in pRL10053 (pOPS0478 conjugated into Rlv3841) | This work |
| Plasmid |  |  |
| pK19mob | pK19-based mobilizable plasmid for single cross-over homologous recombination in *Rhizobium* to make mutants, kanamycin/neomycin-resistant | (3) |
| pOPS0478 | Fragment of pRL10053 (PCR product oxp1751/oxp1752) BD cloned into *Hin*dIII site of pK19mob, kanamycin/neomycin-resistant | This work |
| pOPS0820 | Fragment of RL0688 (*cheA1*) (PCR product oxp2200/oxp2201) BD cloned into *Hin*dIII site of pK19mob, kanamycin/neomycin-resistant | This work |
| pOPS0822 | Fragment of pRL100199 (*fixB*) (PCR product oxp2204/oxp2205) BD cloned into *Hin*dIII site of pK19mob, kanamycin/neomycin-resistant | This work |
| pOPS0823 | Fragment of pRL100195 (*nifB*) (PCR product oxp2206/oxp2207) BD cloned into *Hin*dIII site of pK19mob, kanamycin/neomycin-resistant | This work |
| pOPS0825 | Fragment of pRL90058 (*rmrR*) (PCR product oxp2210/oxp2211) BD cloned into *Hin*dIII site of pK19mob, kanamycin/neomycin-resistant | This work |
| pOPS0831 | Fragment of pRL120205 (*eryB*) (PCR product oxp2222/oxp2223) BD cloned into *Hin*dIII site of pK19mob, kanamycin/neomycin-resistant | This work |
| pOPS0833 | Fragment of RL0149 (PCR product oxp2226/oxp2227) BD cloned into *Hin*dIII site of pK19mob, kanamycin/neomycin-resistant | This work |
| pOPS0836 | Fragment of RL1545 (PCR product oxp2232/oxp2233) BD cloned into *Hin*dIII site of pK19mob, kanamycin/neomycin-resistant | This work |
| pOPS0837 | Fragment of RL2606 (*purC1*) (PCR product oxp2234/oxp2235) BD cloned into *Hin*dIII site of pK19mob, kanamycin/neomycin-resistant | This work |
| pOPS0838 | Fragment of RL3549 (*glnII*) (PCR product oxp2236/oxp2237) BD cloned into *Hin*dIII site of pK19mob, kanamycin/neomycin-resistant | This work |
| pOPS0839 | Fragment of RL3654 (*pssD*) (PCR product oxp2238/oxp2239) BD cloned into *Hin*dIII site of pK19mob, kanamycin/neomycin-resistant | This work |
| pOPS0846 | Fragment of pRL120198 (PCR product oxp2252/oxp2253) BD cloned into *Hin*dIII site of pK19mob, kanamycin/neomycin-resistant | This work |
| pOPS0849 | Fragment of RL1496 (*iolR*) (PCR product oxp2258/oxp2259) BD cloned into *Hin*dIII site of pK19mob, kanamycin/neomycin-resistant | This work |
| pOPS0851 | Fragment of RL3986 (*ruvC*) (PCR product oxp2262/oxp2263) BD cloned into *Hin*dIII site of pK19mob, kanamycin/neomycin-resistant | This work |
| pRK2013 | *E. coli* helper plasmid for tri-parental matings | (4) |
| pSAM_Rl | Plasmid carrying mariner transposon for INSeq library preparation, kanamycin/neomycin- and ampicillin-resistant | (5) |
| Primer |  |  |
| oxp1072 | TGATTACGCCAAGCTAATCCGATATTTTCTCGA  GATTGCC | This work |
| oxp1073 | GCAGGCATGCAAGCTTCGATAGAAGCTTGCTG ATTATCTG | This work |
| oxp1751 | TGATTACGCCAAGCTATGGTTGCCATCAAGC | This work |
| oxp1752 | GCAGGCATGCAAGCTTCTTTGAAGCGATCACGGGC | This work |
| oxp2200 | TGATTACGCCAAGCTATGACGCCACCCTGCTG | This work |
| oxp2201 | GCAGGCATGCAAGCTGCAGGCTCATGGTGAAGAC | This work |
| oxp2204 | TGATTACGCCAAGCTCACGTCTGGGTCTTCATGGA | This work |
| oxp2205 | GCAGGCATGCAAGCTCCGGACTGACCAATTTGTCG | This work |
| oxp2206 | TGATTACGCCAAGCTAGGTGCTTGCCGTCGC | This work |
| oxp2207 | GCAGGCATGCAAGCTATCCCCTTTAGTGAGACTGCA | This work |
| oxp2210 | TGATTACGCCAAGCTTGCCGAAGAAAAACCCGC | This work |
| oxp2211 | GCAGGCATGCAAGCTTTGGACGACCTTGGTGAAGT | This work |
| oxp2222 | TGATTACGCCAAGCTAACGGCGCGGGGATAG | This work |
| oxp2223 | GCAGGCATGCAAGCTCAGCGGCGCCTTCTCC | This work |
| oxp2226 | TGATTACGCCAAGCTGCAACGTCAGACAGTTTCGC | This work |
| oxp2227 | GCAGGCATGCAAGCTCCGTCATGAGGGTGA | This work |
| oxp2232 | TGATTACGCCAAGCTCGGTATCGGCTATTTCACGG | This work |
| oxp2233 | GCAGGCATGCAAGCTACGAGCTTGTTCTCCCGG | This work |
| oxp2234 | TGATTACGCCAAGCTACGATGCCACTGCCTTCA | This work |
| oxp2235 | GCAGGCATGCAAGCTCTTCGGAATAGGCTTCGAG | This work |
| oxp2236 | TGATTACGCCAAGCTGGATCCTCGACGCAGCAG | This work |
| oxp2237 | GCAGGCATGCAAGCTTGACGAAGGAGTGCGGC | This work |
| oxp2238 | TGATTACGCCAAGCTAGTTCTCGCTGCCTCGTC | This work |
| oxp2239 | GCAGGCATGCAAGCTAGCGTCGCAATATGGCCG | This work |
| oxp2252 | TGATTACGCCAAGCTCGATCGAGCTGACGGGAC | This work |
| oxp2253 | GCAGGCATGCAAGCTTCTTTGCGATGGAGGCCG | This work |
| oxp2258 | TGATTACGCCAAGCTCTCCGATGTCCAGCCGTC | This work |
| oxp2259 | GCAGGCATGCAAGCTCACCTCGAACCAGTGCGT | This work |
| oxp2262 | TGATTACGCCAAGCTTCCGGAACCGTGACCTCT | This work |
| oxp2263 | GCAGGCATGCAAGCTCGGCGTCGTTACCCTTGA | This work |

**Legend for Dataset S1** (separate file). Complete Rlv3841 INSeq results for input1, rhizosphere, root, input2, nodule bacteria and bacteroid libraries. HMM classifications: ES; essential, GD; growth-defective, NE; neutral, AD; advantaged. N/A means there are no TA sites in the gene.
